# Assessing human genome-wide variation in the Massim region of Papua New Guinea and implications for the Kula trading tradition

**DOI:** 10.1101/2022.02.06.479284

**Authors:** Dang Liu, Benjamin M. Peter, Wulf Schiefenhövel, Manfred Kayser, Mark Stoneking

## Abstract

The Massim, a cultural region encompassing the southeastern tip of mainland Papua New Guinea (PNG) and nearby islands, is famous for the Kula trading network, in which different valuables circulate in different directions among the islands. To explore Massim genetic history, we generated genome-wide data from across the region and found variable levels of Papuan-related (indigenous) ancestry, including a novel ancestry associated with Rossel Island, and Austronesian-related ancestry that arrived later. We find genetic evidence for different patterns of contact across PNG, the Massim, and the Bismarck and Solomon Archipelagoes for Austronesian-related vs. Papuan-related ancestry, with more geographic restriction for the latter. Moreover, Kula-practicing groups share more genetic similarity than do other groups, and this likely predates the origin of Kula, suggesting that high between-group contact facilitated the formation of Kula. Our study provides the first comprehensive genome-wide assessment of Massim inhabitants and new insights into the fascinating Kula system.

## Introduction

New Guinea, including its eastern part known as Papua New Guinea (PNG), has a long history of human occupation and harbors extensive ethnolinguistic diversity. Modern humans first colonized Sahul (the connected Australia – New Guinea land mass) at least 47 thousand years ago (kya), and perhaps as long ago as 65 kya (*1–4*), while ~3 kya the Austronesian expansion/settlement brought a second wave of human migration into the region (*5, 6*). This long-term isolation of human occupation and different episodes of human migration promoted diverse regional culture developments, including an independent invention of farming in the New Guinea highlands at least 4-5 kya (*7, 8*) and the Lapita culture associated with the expansion of Austronesians from Southeast Asia into the Pacific (*6, 9*).

Geographically, the Massim encompasses the southeastern tip of mainland PNG and nearby offshore islands (Fig. 1) and is considered by anthropologists to be a culturally-defined region (*10, 11*). The Massim region is well-known for the Kula Ring tradition (or Kula), a network trading system in which two types of unique necklaces circulate among the islands in opposite directions, facilitating inter-island social and economic relationships (*12–15*). The Kula connects the islands of the northern and the western Massim (including the eastern tip of Mainland PNG) and also includes Misima, the most northern island of the southern Massim, while other islands of the southern Massim are not involved in the Kula (Fig. 1). Due to seasonal wind conditions, traders on Kula voyages often spend long periods of time on different islands they do not originate from (*12, 14*), which might further facilitate interactions, including genetic ones (*16*), between the groups participating in Kula.

**Fig. 1.**
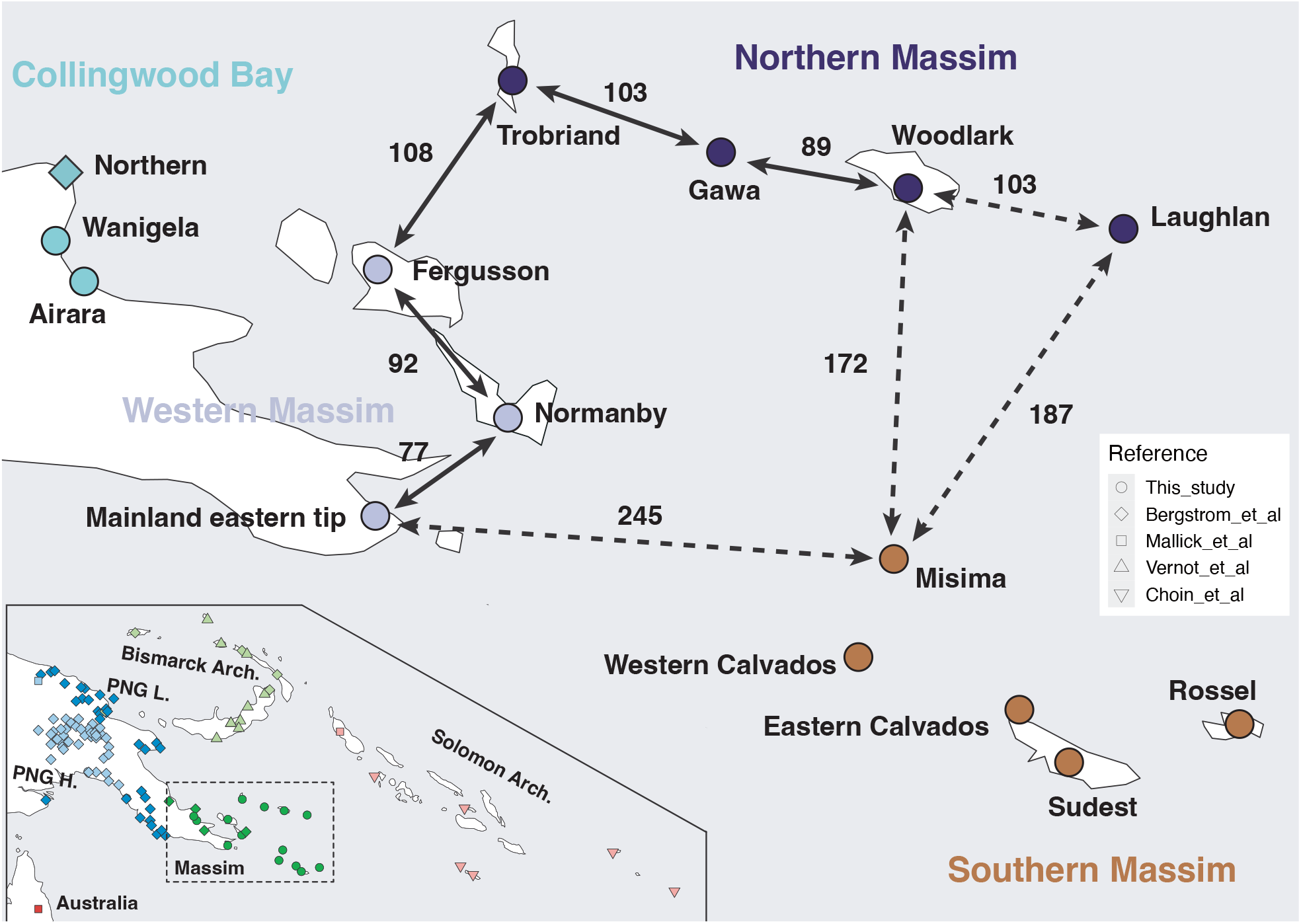
Map showing the location of the newly-studied Massim groups and groups in the reference dataset. The main map shows the location of Massim groups, color-coded according to sub-region, while the inset map shows the location of reference groups. Dot shapes indicate the corresponding publication. The solid arrows connect groups frequently involved in a Kula relationship, while the dashed arrows connect groups that marginally involved in a Kula relationship. The numbers indicate distances (in kilometers) between islands.

Since the initial description of Kula by Malinowski in 1922 (*12*), the region has attracted the attention of anthropologists and archaeologists, and chronologies of Massim have been proposed (*17, 18*). The earliest evidence for human occupation of the Massim is dated to 17.3 kya, on Paneati Island in the Louisiades Archipelago (*19*); undated obsidian and stone artifacts that are similar to those dated to the Late Pleistocene or Early Holocene elsewhere in New Guinea further support human occupation around this time, when lowered sea levels connected much of the Massim to the PNG mainland (*20, 21*). After settlement by Lapita people ~2.5-3 kya, the northern and southern Massim exhibited separate cultural developments, until ~500-200 years ago with the formation of Kula (*17*). Linguistically, almost all Massim inhabitants speak languages belonging to the Austronesian language family, except for a few groups living on mainland PNG who speak Papuan languages belonging to the Trans-New Guinea language family (*22*), and the Rossel Islanders on the most eastern tip of the southern Massim, who speak a Papuan language that has been classified either as a language isolate (*23, 24*) or as belonging to the Yele-West New Britain language family, linking Rossel linguistically with the Bismarck Archipelago (*22, 25*).

While the Massim region has been well studied by archaeologists, anthropologists, and linguists, genetic variation in the Massim remains largely unexplored, even though it occupies a key position in connecting the northern and southern coasts of New Guinea as well as connecting New Guinea with the neighboring Solomon Islands and other regions of Oceania. The most comprehensive human genetic study of the Massim to date analyzed mitochondrial DNA (mtDNA) and Y chromosome variation, and found regional genetic-geographic population structure for mtDNA but not for the Y chromosome (*26*). This was interpreted as a potential signature of the Kula, as the travel between islands to perform the trading of the goods is mostly mediated by males, which could reduce inter-island genetic differences for the Y chromosome but not for mtDNA. However, studies of genome-wide data provide much richer information concerning admixture and population history, as shown by two recent genome-wide studies of PNG (*27, 28*), but genome-wide studies of the Massim are lacking as of yet. Here, we report the results of comprehensive genome-wide analyses of the Massim region with resulting insights into the genetic history of Oceania in general and implications for the Kula tradition in particular.

## Results

### Overall genetic variation of Austronesian and Papuan ancestries in the Massim

We generated genome-wide single nucleotide polymorphism (SNP) array data for 192 individuals from 15 groups spanning the entire Massim region including northern, southern and western Massim as well as the mainland parts from Collingwood Bay (Fig. 1). We merged our data with published array and WGS data encompassing East Asia and Oceania and additionally included Africans and Europeans as more distant comparative groups (Fig. 1; fig. S1; table S1). To obtain an overview of the genetic variation and genetic-geographic population structure revealed with our compiled genomic dataset, we first performed principal component analysis (PCA). In a PCA plot focusing on the East Asian and Oceanian groups (Fig. 2; fig. S2), we observed a striking cline with East Asians at one pole and PNG highlanders at the other; the Massim groups fall in between together with groups from the Central Province of PNG, the Bismarck Archipelago (in short, Bismarcks), and the Solomon Archipelago (i.e. Bougainville and the Solomon Islands; in short, Solomons). Groups from the PNG lowlands are placed close to PNG highlanders, except for the Central group, which comprises more Austronesian speakers. Bellona, Renell and Tikopia are Polynesian outliers in the Solomons, i.e. groups that migrated back to the Solomons from Polynesia, which explains their placement further towards East Asians than any other Near Oceanian group studied. The northern Massim groups are closer to the East Asian pole, while the southern Massim group from Rossel is closest to the PNG highlander pole, and other southern Massim as well as all western Massim and Collingwood Bay groups fall in between. Notably, there is a division in the PCA within southern Massim, with Rossel being closest to the PNG mainland and highland groups and Sudest, which geographically is located next to Rossel, being close to Rossel, while the other southern Massim groups fall together with northern and western Massim groups and those from Collingwood Bay.

**Fig. 2.**
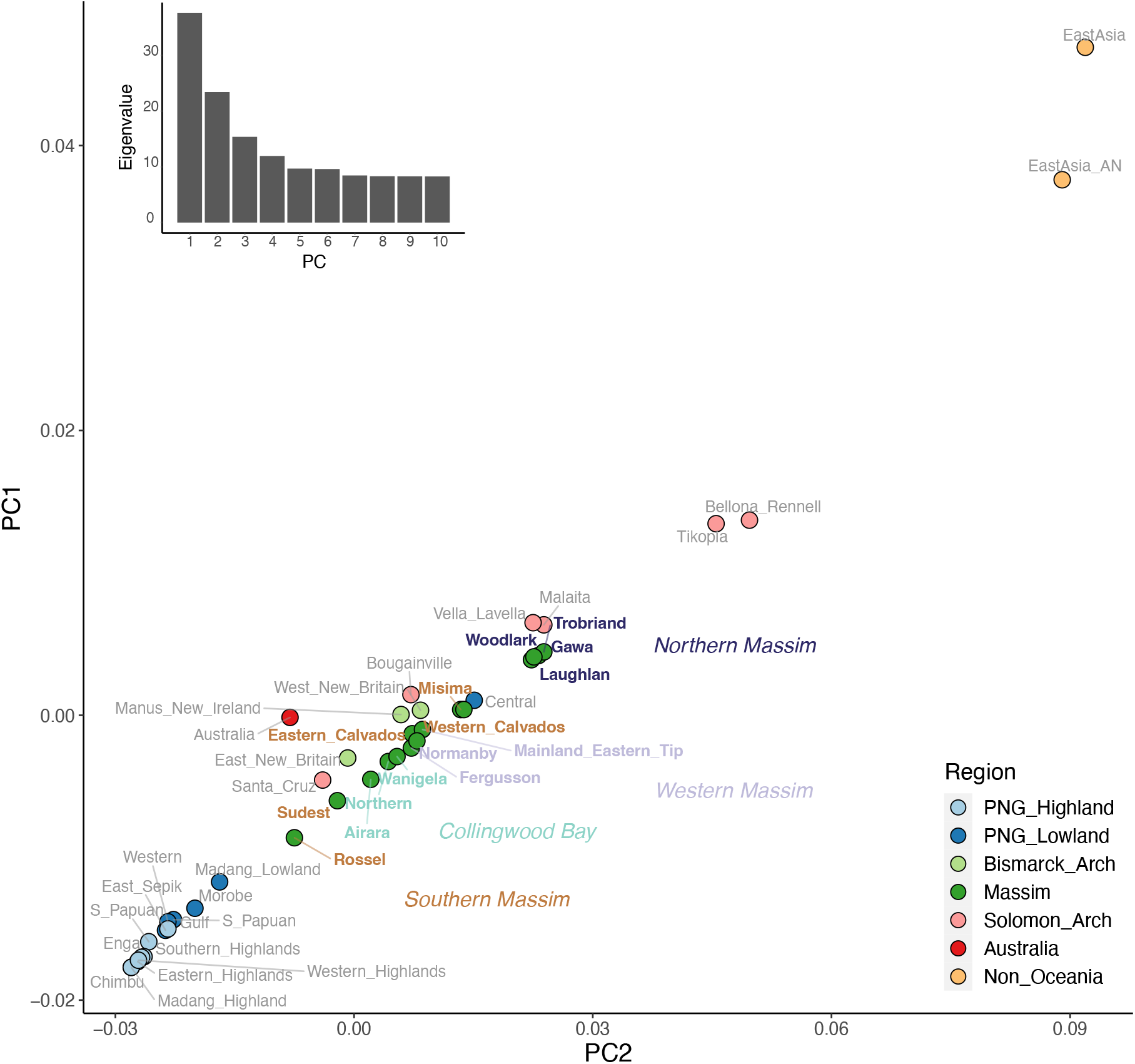
Principal component analyses (PCA) of the East Asian and Oceanian groups. Plot of PC1 vs. PC2 of the median position of the East Asian and Oceanian groups from fig. S2, colored according to region; Massim regions are further indicated. The eigenvalues from PC1 to PC10 are shown on the top left.

Since the PCA suggests mixed East Asian-Papuan ancestry in the Massim groups, we explored this further in an ADMIXTURE analysis. At K=2, one component (designed in pink) is most enriched in the East Asian Austronesian groups while the other component (blue) is enriched in the Papuan highlanders (Fig. 3); all other Oceanian groups (including the Massim) exhibit a mixture of these two genetic ancestry components, albeit in different proportions. The northern Massim individuals, who speak Austronesian languages, have the highest amount of Austronesian-related ancestry (~52% pink), while the Papuan-speaking Rossel islanders from southern Massim have the lowest amount (~20% pink), and hence the highest amount of Papuan (~80% blue) ancestry (table S2). However, several Austronesian-speaking groups in the Massim, including those from Sudest, the western Massim, and Collingwood Bay, also have more Papuan than Austronesian ancestry, suggesting substantial contact between Papuan and Austronesian groups.

**Fig. 3.**
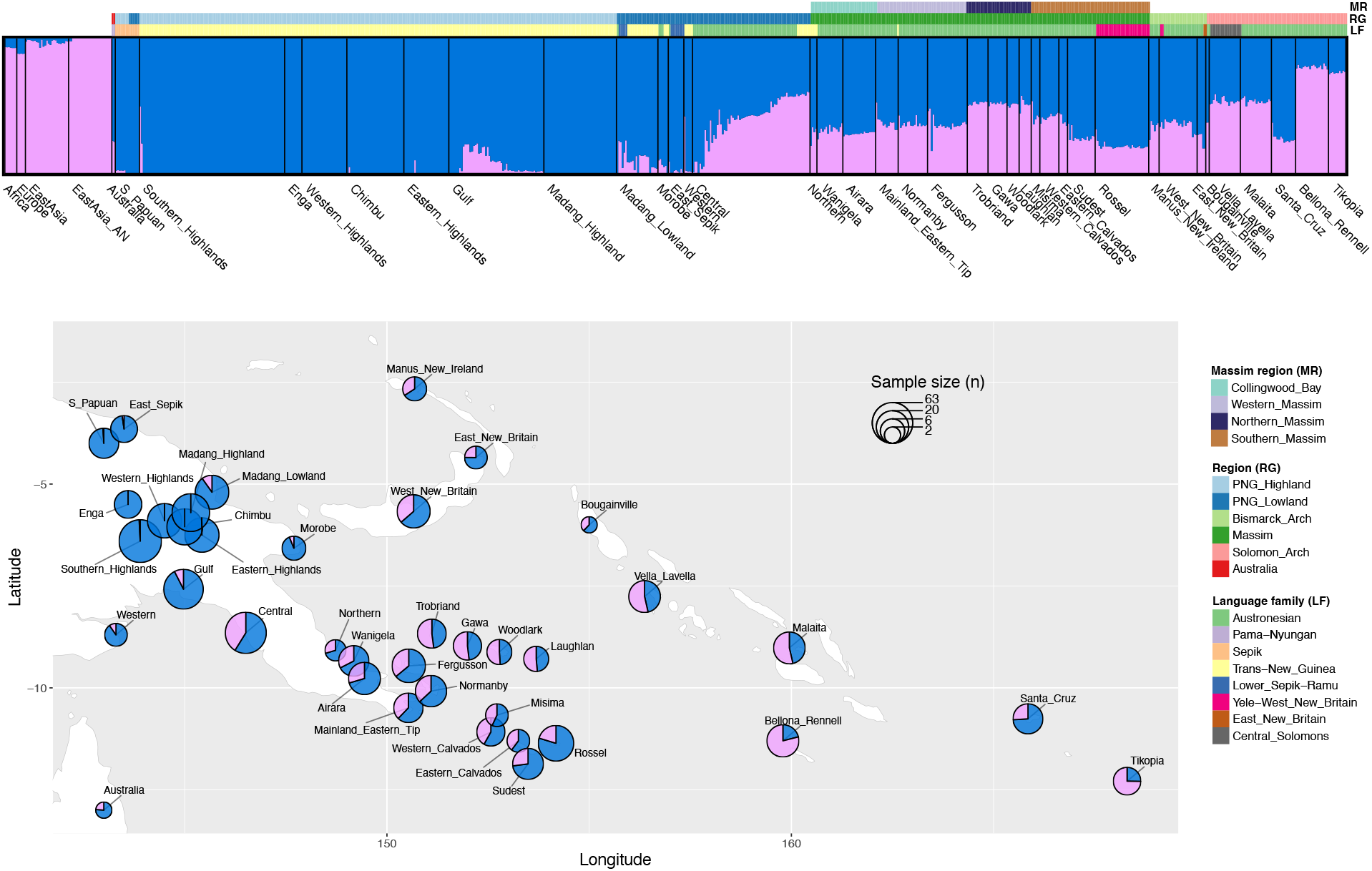
K=2 ADMIXTURE analyses focusing on Oceanian structure. ADMIXTURE results for K=2 at the top, plotted on a map at the bottom. Color bars on the top indicate the region (RG), language family (LF) for the Oceanian groups and Massim region (MR) for the Massim groups. The size of the circle for each group is proportional to the sample size.

This large range in variation of Austronesian (~20-52%) vs. Papuan genomic ancestry in the Massim exceeds that of nearby, larger island regions: Austronesian ancestry varies between ~25-36% in the Bismarcks and between ~38-54% in the Solomons (excluding Santa Cruz and Polynesian outliers) (Fig. 3). We therefore investigated the Austronesian vs. Papuan ancestry in the Massim in more detail via outgroup f3 statistics. A plot of the f3 values measuring shared drift between Oceanian populations and East Asian Austronesians, vs. the f3 value measuring shared drift between Oceanian populations and PNG highlanders (Fig. 4), gives results similar to the PCA plot (Fig 2), namely that Massim groups harbor both Austronesian and Papuan ancestry with different proportions depending on their locations within the Massim region. As before, northern Massim groups have more Austronesian ancestry while Rossel Islanders from southern Massim have more Papuan ancestry. As in the PCA plot, the Massim groups are distributed along a linear cline between PNG highlanders with high amounts of Papuan ancestry at one pole, and Polynesian outliers in the Solomons with high amounts of Austronesian ancestry at the other pole. Moreover, the shape of the f3 plot suggests a single major admixture episode rather than continuous gene flow or a tree-like history (*29*).

**Fig. 4.**
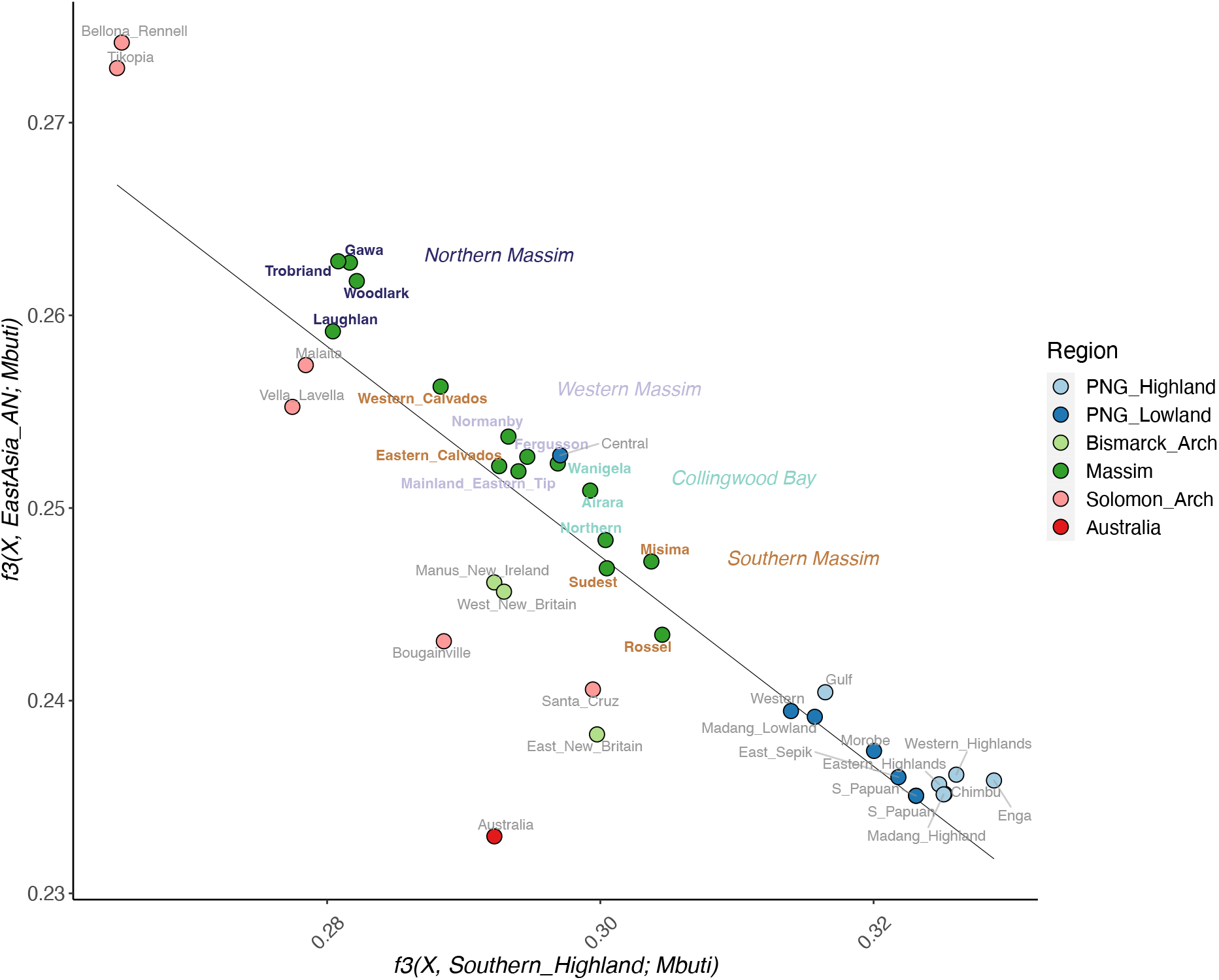
F3 statistics measuring shared genetic drift between Oceanians and either East Asian Austronesians vs. between Oceanians and PNG highlanders. The value of f3(X, Southern province highlanders; Mbuti) is on the x-axis, and the value of f3(X, East Asian Austronesians; Mbuti) is on the y-axis. X denotes the Oceanian groups, colored according to region, with Massim regions further highlighted. The linear regression line was computed using all values shown on the plot.

In addition to the drift/allele sharing analyses, a haplotype-based method (GLOBETROTTER) also suggests a single pulse of admixture for all the Massim groups except for Rossel (fig. S3); Rossel exhibits an unclear admixture signal probably due to the low amount of Austronesian ancestry. Moreover, it is likely that the incoming population was already admixed (fig. S3) i.e., had both Austronesian and Papuan ancestry. Consistently, the percentage of modeled Austronesian-related ancestry is highest in the northern Massim groups (~42%), and the Papuan-related ancestry is highest in the Rossel group from southern Massim (~73%; table S2). GLOBETROTTER also infers admixture dates of ~1-3 kya for the Massim groups, and similar dates are inferred with another method, ALDER (fig. S4). The inferred dates are not correlated with the amount of Austronesian ancestry (fig. S5), suggesting that the higher amounts of Austronesian ancestry in some Massim groups cannot be explained by a longer period of contact, but instead must reflect other social circumstances.

### Austronesian and Papuan ancestries specific to the Massim

In the ADMIXTURE analysis, the lowest cross-validation error (and hence, the best-supported number of ancestry components) occurred at K=8 (fig. S6). Six ancestry components were found in the Oceanians (Fig. 5); the other two were restricted to Africans, and to Europeans and East Asians, respectively. The ancestry components found in Oceanians include:

1. a light green component at highest frequency in Chimbu and Madang highlanders and also enriched in Western and Eastern highlanders and S_Papuans (S for SGDP (*30*)) from the East Sepik; this component is present at uniformly low frequencies across the Massim.
2. a blue component at highest frequency in Southern PNG highlanders and Enga and enriched in other mainland PNG groups; this component is present up to medium frequency in the Massim, Bismarcks and Santa Cruz.
3. a pink component enriched in Central PNG that exists in low frequency in a few other lowland PNG groups, the Solomons (including the Polynesian outlier Tikopia, but not the other Polynesian outlier, Bellona_Rennell), and the Bismarcks; and is the major component in all Massim groups except Rossel and neighboring Sudest.
4. a black component that is at highest frequency in the Rossel group, second highest in the neighboring group from Sudest, and at lower frequency in several other groups from the western and southern Massim.
5. a deep green component enriched in Bougainville, the Bismarcks and the Solomons (including the Polynesian outlier Tikopia, but not the other Polynesian outlier, Bellona_Rennell); this component is present at very low frequency in some PNG mainland groups but notably absent from the Massim.
6. a peach component that is the only component in the Polynesian outlier Bellona_Rennell, and is found in high frequency in Tikopia and in low frequency in other Solomons and in Bismarck groups; this component is not found in the Massim.

**Fig. 5.**
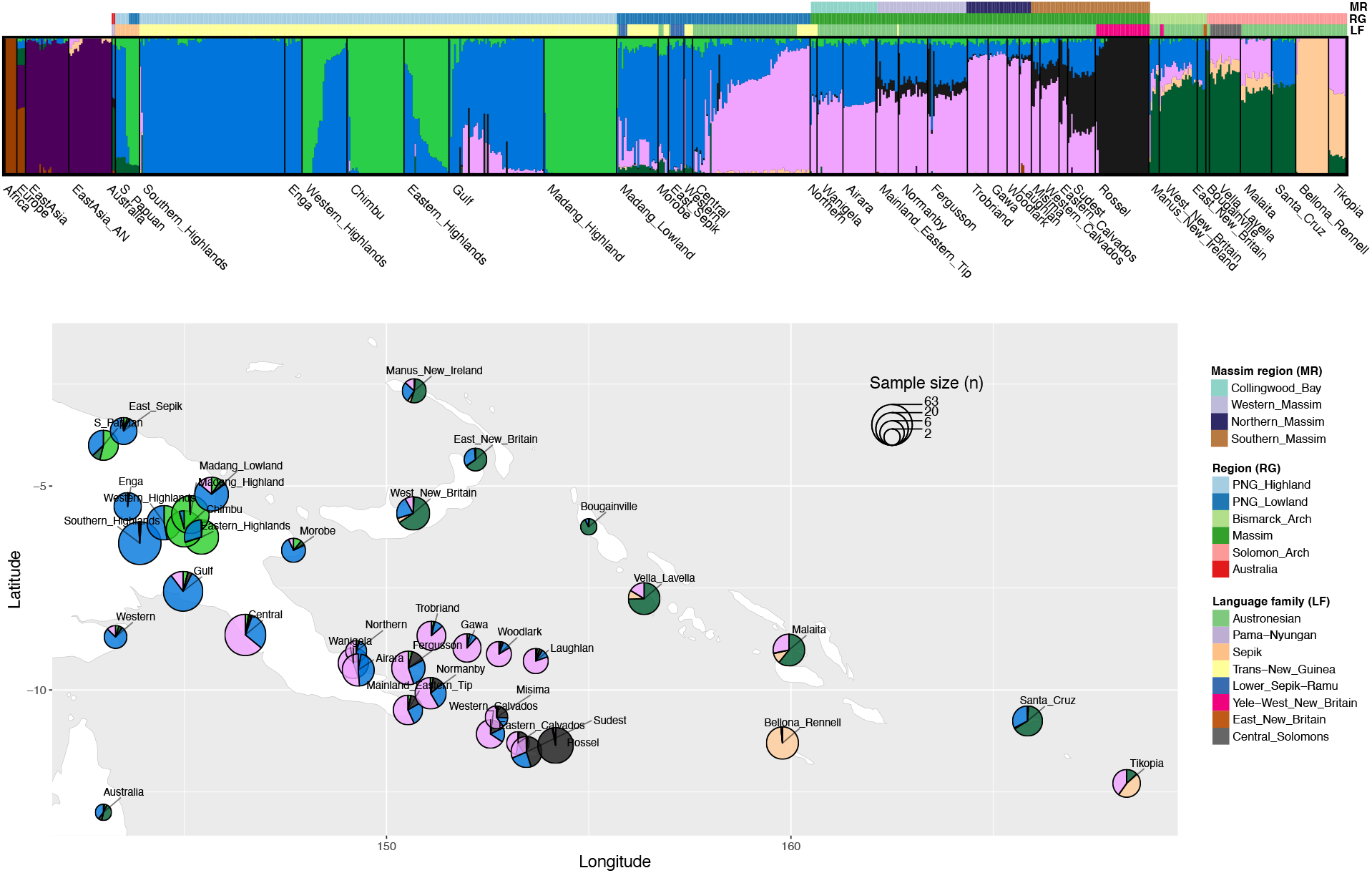
K=8 ADMIXTURE analyses focusing on Oceanian structure. ADMIXTURE results for K=8 on the top, plotted on a map at the bottom. Color bars on the top indicate the region (RG), language family (LF) for the Oceanian groups and Massim region (MR) for the Massim groups. The size of the circle for each group on the map is proportional to the sample size.

The blue and pink components detected at K=8 (Fig. 5) overlap largely, but not completely, with the results for the Oceanian groups observed at K=2 (Fig. 3). All of the groups exhibiting any of the four components newly detected with K=8 (light green, black, dark green, and peach) in the Oceanians have more Papuan than Austronesian ancestry detected at K=2 (Fig. 3) except for the Polynesian outliers (peach), which suggests substantial variation in Papuan ancestry as well as a drifted Austronesian ancestry in the Polynesian outliers. Hence, we further investigated the Papuan and Austronesian ancestry-specific variation in the Massim.

We first calculated f4 statistics of the form f4(Test groups, East Asian Austronesians; Chimbu/Bougainville/Rossel, Southern Highlanders) to test for different affinities of each test group (all other Oceanian groups) with Chimbu, Bougainville, or Rossel-related ancestry when compared to Southern Highlanders. To control for different amounts of Austronesian-related ancestry in the test groups, we plotted these against an f4 statistic of the form f4(Test groups, Southern Highlanders; East Asian Austronesians, Australians), which as shown previously provides a measure of Austronesian-related ancestry in the test groups (*31*). Groups exhibiting significant deviations from the regression line in each plot would indicate differential Papuan ancestries in those groups. The results for the comparison of Chimbu vs. Southern Highlanders shows no significant deviations from the regression line for any Oceanian group (Fig. 6A), indicating that all of the groups tested have equal affinities with Chimbu and Southern Highlanders ancestry. In the comparison between Bougainville and Southern Highlanders (Fig. 6B), several of the groups from the Bismarcks and the Solomons exhibit stronger affinities with Bougainville, as noted previously (*32*), while several PNG groups exhibit stronger affinities with the Southern Highlanders; however, none of the Massim groups show any preferential affinities. In the comparison between Rossel and Southern Highlanders (Fig. 6C), again several PNG groups exhibit stronger affinities with Southern Highlanders, along with the Polynesian Outliers from the Solomons. Interestingly, there is variation in genomic ancestry among the Massim groups: southern Massim groups share more ancestry with Rossel; Collingwood Bay groups (from mainland PNG) share more ancestry with the Southern Highlanders; and the remaining Massim groups do not share excess ancestry with either Rossell or Southern Highlanders. These results strongly support the existence of more than one distinct Papuan-related ancestry in the Massim groups (one related to Rossel, and at least one more related to other Papuan groups).

**Fig. 6.**
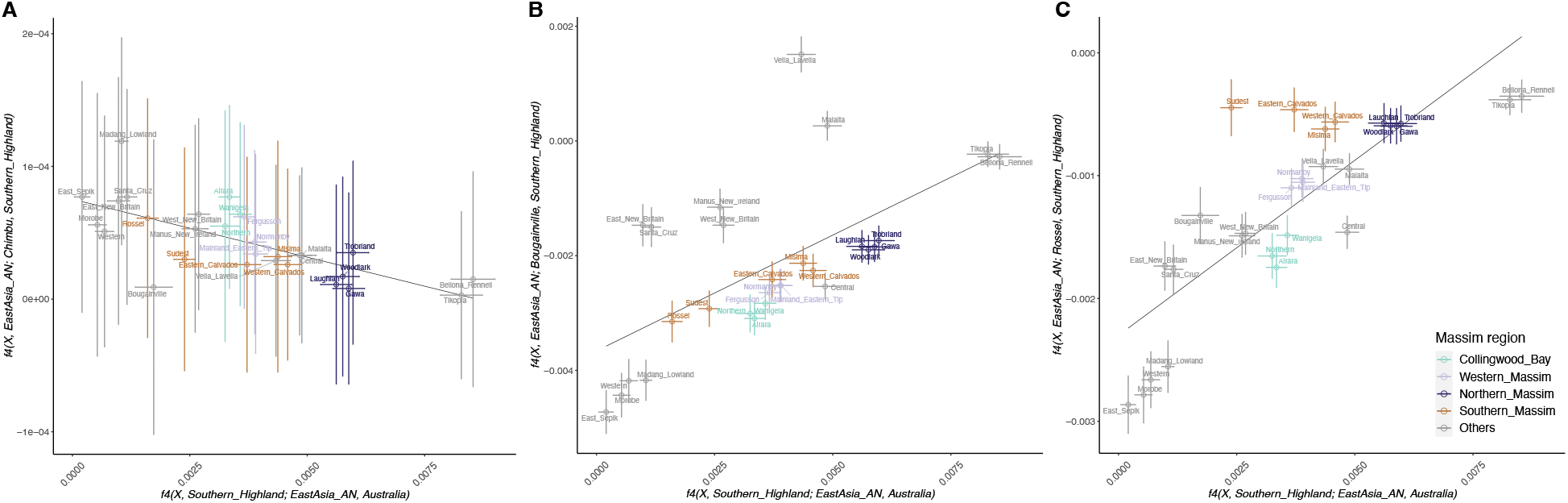
F4 statistics measuring differential PNG ancestry affinities of Oceanian groups in respective to East Asian ancestry affinity. The value of f4(Oceanian groups, Southern province highlanders; East Asian Austronesians; Australians) is on the x-axis, and the value of f4(Oceania groups, East Asian Austronesians; (A) Chimbu / (B) Bougainville/ (C) Rossel islanders; Southern Highlanders) is on the y-axis. X denotes the Oceanian groups, colored according to Massim region. Error bars indicate +/− three times standard error. Linear regression lines were computed using all point values shown on the plot.

We next used f4 statistics to investigate variation in Austronesian-related ancestry in the Massim groups. We first confirmed that the non-Papuan ancestry in the Massim groups is associated with Austronesian ancestry by redoing the plots in Fig. 6 and changing the f4 statistic on the X-axis of each plot to f4(Test groups, Southern Highlanders; East Asian Austronesians, non-Austronesian East Asians). The results (fig. S7) show the same pattern as Fig. 6, hence confirming that the Asian-related ancestry in all of the tested groups is indeed associated with Austronesian ancestry rather than some other East Asian-related ancestry. We followed this with an f4 statistic of the form f4(Test groups, Southern Highlands; East Asian Austronesians, Bellona_Rennell), plotted as before against f4(Test groups, Southern Highlands; East Asian Austronesians, Australians/non-Austronesian East Asians) to account for overall Austronesian-related ancestry, to test for differential affinities of test groups with East Asian Austronesians vs. Polynesian outliers (using Bellona_Rennell as a proxy). The results (fig. S8) indicate that the Bismarck and Solomon groups share excess ancestry with Polynesian outliers, while the Massim groups do not show any excess affinities with either East Asian Austronesians or Polynesian outliers.

We then used haplotype-based methods to verify the above inferences based on allele-sharing statistics. A ChromoPainter analysis in which all Massim groups except Rossel are used as recipients, and Rossel is included with other Oceanian groups as donors, confirms the distinct Papuan ancestry profiles in different Massim groups (fig. S9). In particular: Collingwood Bay groups from the PNG mainland share most ancestry with mainland PNG lowlander and highlander donors; western and northern Massim groups share more ancestry with Rossel and then with Central Province lowlander donors; and southern Massim groups share by far the most ancestry with Rossel islanders. The ChromoPainter results also show higher sharing between northern Massim and East Asian Austronesians, Central Province lowlanders, and Solomon groups, indicating not only higher Austronesian ancestry in northern Massim but also that their Austronesian ancestry is shared with these other groups.

To study the ancestry-specific variation we detected in the Massim in more detail, we applied local ancestry inference via RFMix and extracted Austronesian and Papuan ancestry-specific segments. The global ancestry proportions computed from the local ancestry inferred segments (table S2) showed that the northern Massim groups have the highest levels of Austronesian-related ancestry (~41-43%), while Rossel and its geographic neighbor Sudest have the highest Papuan-related ancestry (~73-80%). The amount of Austronesian-related ancestry inferred by RFMix is slightly lower than that inferred by the ADMIXTURE and GLOBETROTTER analyses, while similar proportions of Papuan ancestries are inferred by all three methods. This suggests that the Papuan ancestry-specific segments are identified with high confidence, but there is more uncertainty in inferring Austronesian-related ancestry segments, probably due to a poorer proxy for this source. A PCA based on Papuan ancestry-specific segments shows three poles, consisting of: the Bismarck and Solomon groups, Rossel, and Papuan highlanders (Fig. 7A; fig. S10A). The other Massim groups are placed towards Rossel, with Massim groups on/nearby the mainland (Collingwood Bay and western Massim) closer to the Papuan lowlanders, consistent with the ChromoPainter results. The identification of a novel Papuan ancestry associated with Rossel probably reflects strong genetic drift, as further evidenced by the high degree of sharing of identity-by-descent (IBD) fragments within this group (fig. S11). In contrast to the Papuan ancestry-specific PCA, in the Austronesian ancestry-specific PCA, all of the Oceanian groups fall together in a cluster quite distinct from East Asian groups (Fig. 7B; fig. S10B). Overall, the results from the haplotype-based methods (ChromoPainter and RFMix) are quite consistent with the results from the allele-sharing statistics (f3 and f4).

**Fig. 7.**
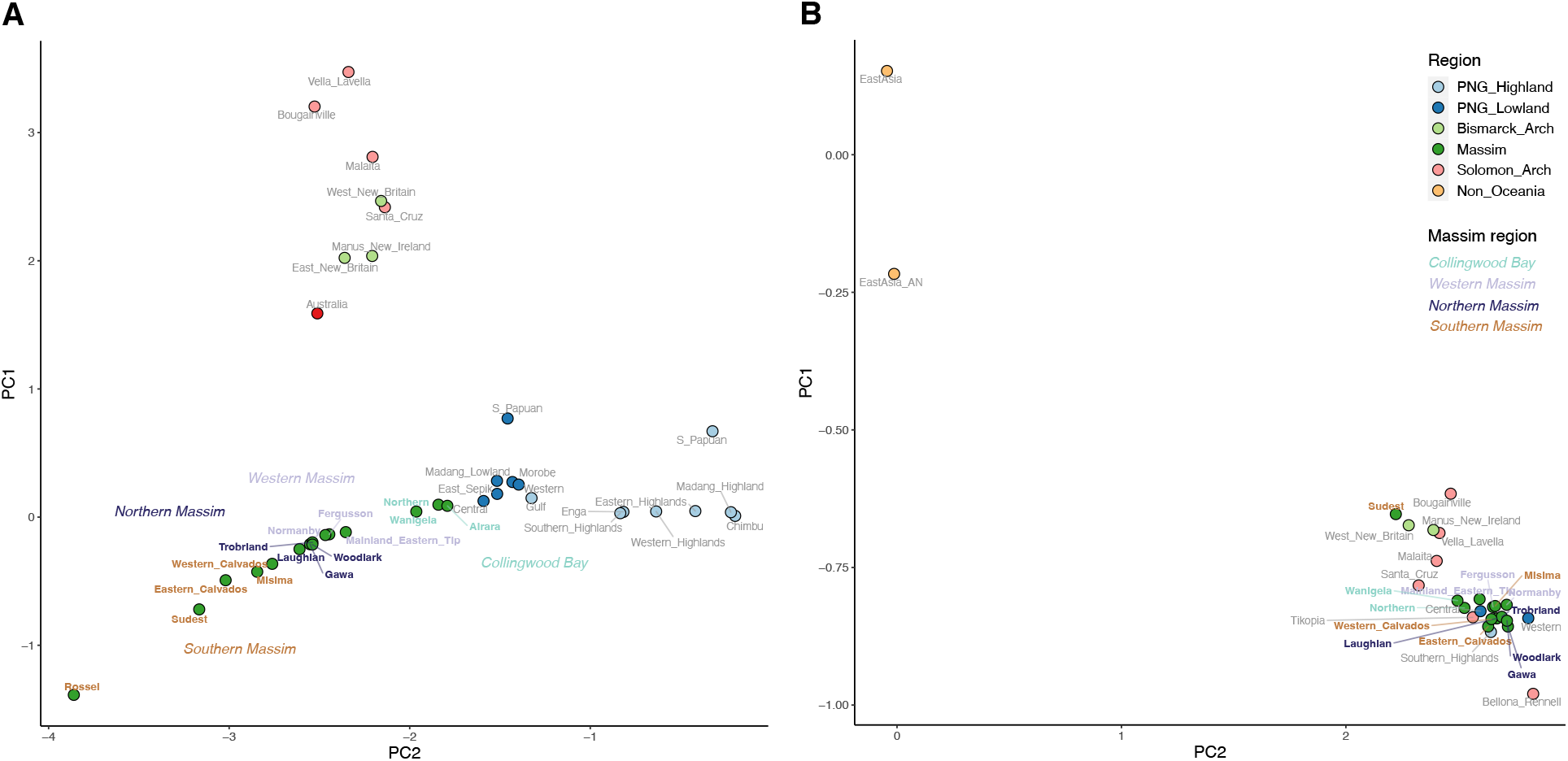
Papuan/Austronesian ancestry-specific PCA of Oceanian (and East Asian) groups. Plot of median position of PC1 vs. PC2 for individuals from Oceanian groups (see fig. S10 for the PCA plot of all Oceanian individuals) and using only (A) Papuan or (B) Austronesian segments identified by RFMix, colored according to region; Massim regions are further highlighted.

### Recent contact involving Massim groups and implications for the Kula tradition

To investigate in more detail genetic contacts involving the Massim groups in the recent past, we analyzed sharing of identify-by-descent (IBD) segments. Different size ranges of IBD segments are informative about contact at different times, extending back to ~4 kya (*33*). In the range of 1 to 5 cM, which corresponds to an average of ~2.7 kya, i.e. around the time of the arrival of Austronesians to the region (*9, 17*), there is strong IBD sharing among Oceanian groups, in particular among the Massim, the Bismarcks and Solomons, and among the Papuan highlanders (Fig. 8A; fig. S12). In the range of 5 to 10 cM, corresponding to ~675 ya, the strong connection between mainland Papua New Guinea and the offshore islands in Massim is no longer there, but the connection between northern and southern Massim remains. In the range of over 10 cM or ~225 ya, strong sharing is only seen within some parts of the Papuan highlands, northern Massim, and southern Massim, respectively. To further test the idea that IBD sharing in the range of 1 to 5 cM reflects the Austronesian settlement, we extracted and analyzed Austronesian ancestry-specific and Papuan ancestry-specific IBD segments. The results show that in the range of 1 to 5 cM, Austronesian ancestry-specific IBD segments account for most of the sharing between regions, while Papuan ancestry-specific IBD sharing shows a clear distinction between the PNG mainland and Massim groups, vs. the Bismarcks and Solomons (Fig. 8B and 8C; figs. S13 and S14). In particular, the strong sharing between many Massim groups and the Central Province group in the ChromoPainter analysis (fig. S9) can be attributed to Austronesian-related ancestry, as this accounts for the IBD-sharing between them (figs. S12 and S13). And, the closer relationship in Papuan-related ancestry between the PNG mainland and Massim groups, vs. either of these and the Bismarck/Solomon groups, is consistent with the PCA based on Papuan-related ancestry (Fig. 7).

**Fig. 8.**
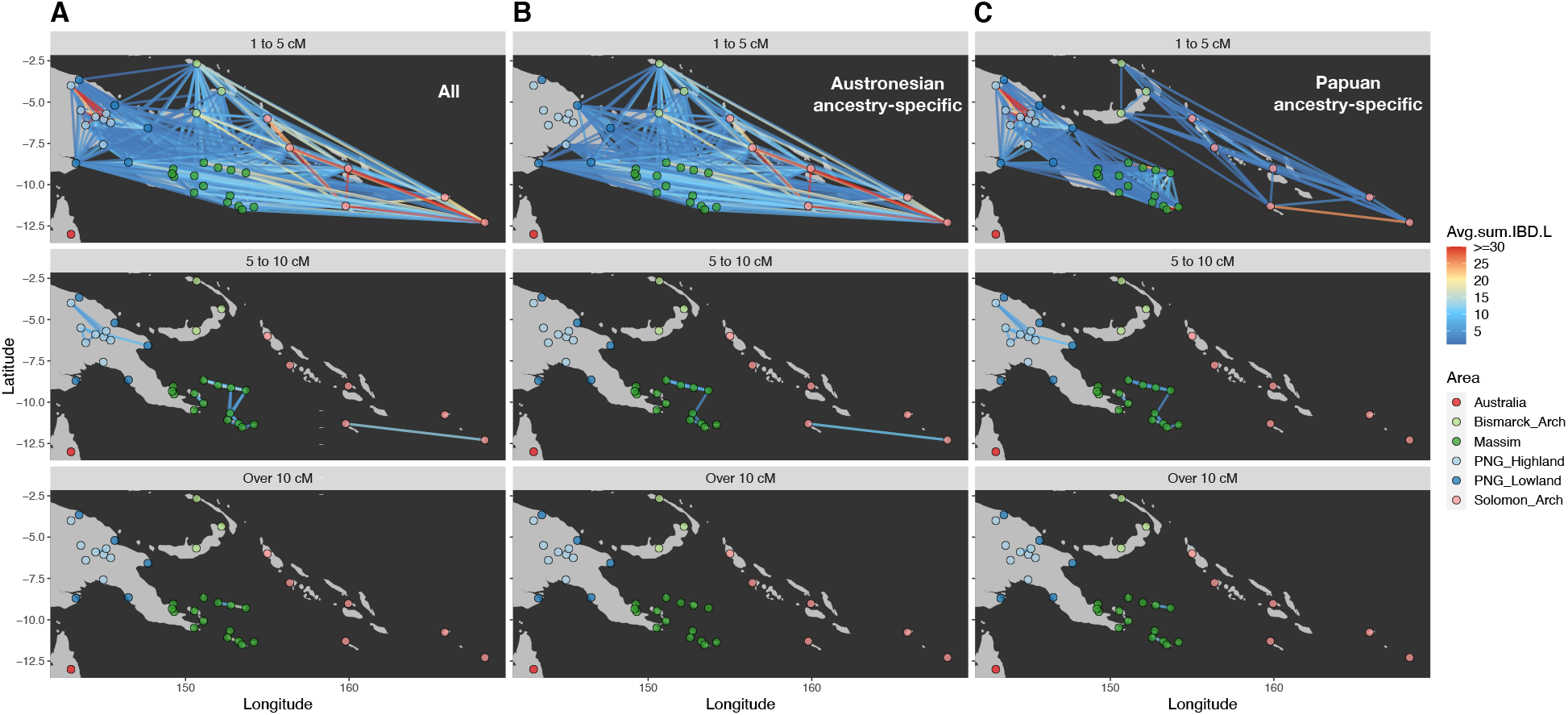
IBD-sharing network for Oceanians. Network visualizations of the mean summed length of: (A) all shared IBD blocks, (B) shared Papuan ancestry-specific IBD blocks; and (C) shared Austronesian ancestry-specific IBD blocks. Identified IBD blocks are in the range of 1 to 5 cM, 5 to 10 cM, and over 10 cM, with the mean number of shared IBD blocks equal to 0.5 or more. The group points are colored according to region. The heat plot segments are proportional to the average of the summed IBD length (cM).

Finally, to investigate the potential relationship between the Kula tradition and genomic variation, we compared the IBD sharing between Kula-practicing vs. non-Kula-practicing groups (in short, Kula vs. non-Kula). We measured IBD similarities among Kula/non-Kula taking the differences in IBD sharing within groups into account by dividing the mean IBD sharing between groups by the mean IBD sharing within groups. A plot of IBD similarity vs. geographic distance, for different lengths of IBD segments (corresponding to different time periods) shows that Kula groups exhibit higher IBD similarities than non-Kula groups separated by the same geographic distance (fig. S15). To test for statistical significance and further controlling for background sharing through time by additionally including groups outside Massim, we then computed a relative IBD similarity statistic that takes into account the overall IBD sharing among all Oceanians for different IBD segment size bins. We find that relative IBD similarity statistics are significantly positive for all Massim groups (i.e. both Kula and non-Kula groups), indicating that the Massim region overall shows evidence of more contact among groups than does the rest of Oceania. Moreover, Kula groups show constantly and significantly higher relative IBD similarities than non-Kula groups (Fig. 9; fig S16), even before the formation of the Kula Ring tradition ~500 ya (*14, 17*).

**Fig. 9.**
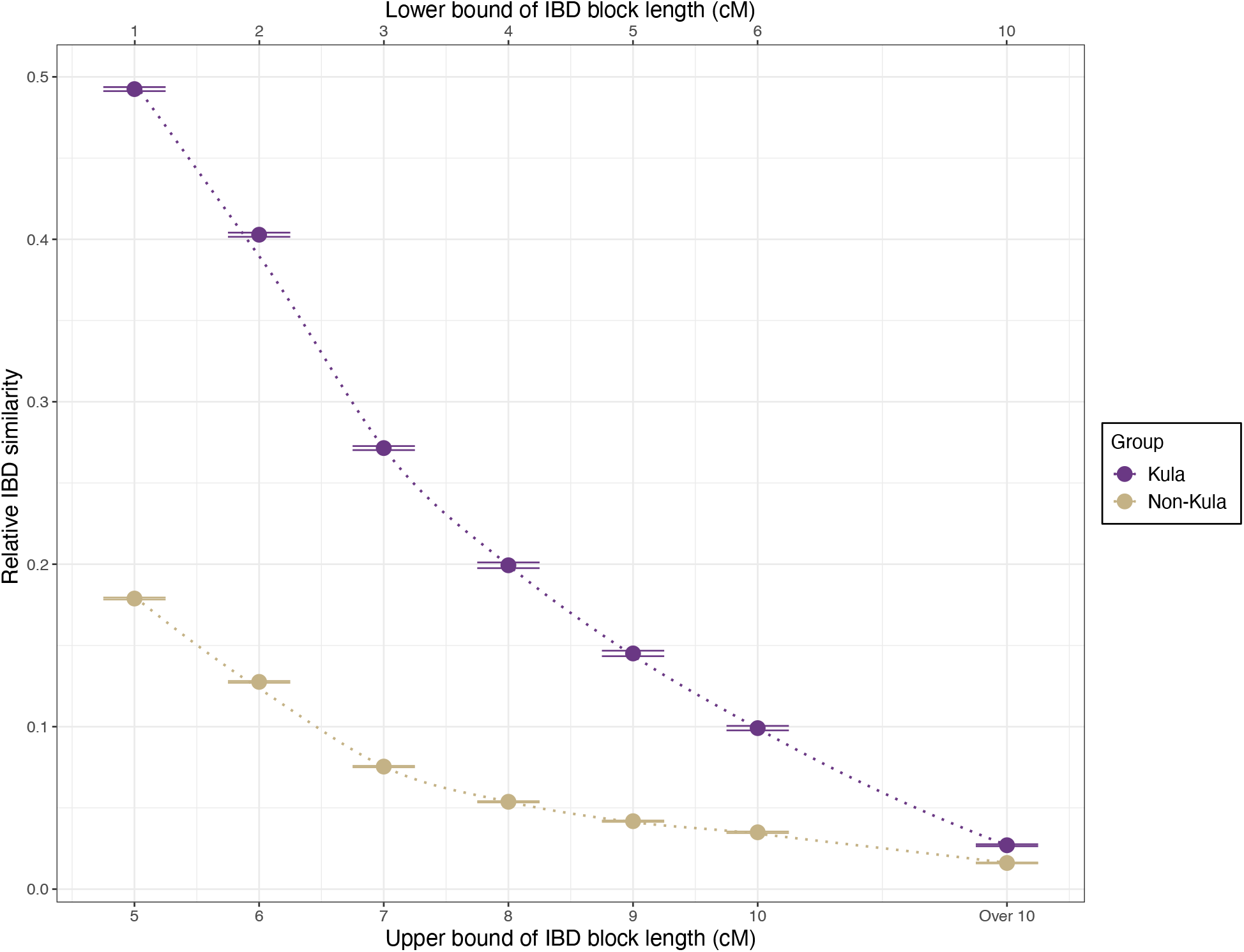
Relative IBD similarities of Kula vs. non-Kula groups. The lower and upper bound lengths of the IBD blocks used in the calculations are indicated at the top and the bottom of the plot, respectively. These intervals correspond to ~2.7, ~1.5, ~1.1, ~0.8, ~0.7, ~0.6, and ~0.2 thousand year ago (kya). The distance between the two horizontal lines on each point indicate +/− three times standard error.

## Discussion

Our analyses of genome-wide data from an extensive sampling of individuals from across the Massim, together with data from other regions, indicate that all Massim groups share both Austronesian-related and Papuan-related ancestry. However, we also find important regional distinctions within the Massim with respect to various aspects of these ancestries.

First, there is more Austronesian-related ancestry in the northern Massim (average based on ADMIXTURE of 52%; Fig. 3, table S2) than in the other regions (average of 34%, with Rossel having the lowest amount, 20%). While this may reflect additional pulses of admixture or more prolonged contact with Austronesians in the northern Massim, the estimated admixture dates do not correlate with the amount of Austronesian ancestry (fig. S5). Moreover, the GLOBETROTTER results suggest single pulses of admixture (fig. S3), as does the linear relationship in the f3 values between the Massim and Papuans vs. those between the Massim and Austronesians (Fig. 4) (*29*). We also do not see any differences in the Austronesian-related ancestry of different Massim groups (fig. S8). These results therefore suggest differences in the contact relationships between the indigenous Papuan groups and the incoming Austronesians in the northern Massim vs. elsewhere in the Massim, with Rossel being the least-impacted by the Austronesians, possibly due to its more remote location. It is also notable that while all of the Massim groups (with the exception of Rossel and 5 individuals from groups on mainland PNG) speak Austronesian languages, Austronesian ancestry is in the minority in most of the groups, and reaches a maximum of 52% in the northern Massim. Thus, the linguistic impact of the Austronesians in the Massim was greater than the genetic impact.

Second, the Papuan-related ancestry of Papuan-speaking Rossel Islanders is distinct from Papuan-related ancestries identified previously and elsewhere in Near Oceania. Previous studies of genome-wide data have shown a major distinction between Papuan ancestries in the PNG Highlands vs. Bougainville (*32*), and in the western vs. eastern regions of the PNG Highlands (*27, 28*). We find support for both of these distinctions, as well as for a novel Papuan-related ancestry on Rossel (Figs. 5 and 7A; fig. S10A) that is also present in lower amounts in other Massim groups, especially from the southern Massim. In addition to this novel Rossel-related Papuan ancestry, Massim groups (except Rossel) also have, at frequencies of 1-63%, another Papuan-related ancestry that is at highest frequency in the western PNG Highlands and the southern PNG lowlands (Fig. 5). Whether this ancestry predates, is associated with, or postdates the arrival of the Austronesians cannot be determined from our data. The existence of a novel Papuan-related ancestry on Rossel could reflect initial colonization by people with this ancestry, subsequent isolation and genetic drift, or both. Further attesting to its isolation is the fact that the unique Papuan-related (non-Austronesian) language, Yélî Dnye, is spoken on Rossel; Yélî Dnye has been variously classified as either a language isolate (*23, 24*) or as possibly related to Anêm and Ata, two languages of West New Britain in the Bismarck Archipelago (*22, 25*). Moreover, pottery was introduced relatively late on Rossel, around 500-550 years ago, compared to around 2.8 kya elsewhere in the southern Massim (*20*). The very high amount of sharing of IBD segments within Rossel (fig. S11) further supports isolation and genetic drift as responsible for the development of the novel Papuan-related ancestry on Rossel. We further note that the genetic isolation of Rossel in comparison to other Massim groups was not as apparent in a previous study of mtDNA and Y-chromosome variation in these same samples (*26*), attesting to the value of genome-wide data for studies of human population history.

Third, there is a striking contrast in patterns of sharing of IBD segments of Austronesian vs. Papuan ancestry between groups. There is extensive sharing of short IBD segments (1 to 5 cM) across the studied Massim region (Fig. 8A), and Austronesian segments are widely shared among the PNG lowland and island groups but not with the highlands (Fig. 8B), in keeping with previous observations of a lack of Austronesian-associated ancestry in the PNG highlands (*27, 28, 32, 34*). However, sharing of Papuan segments is strictly either among mainland PNG and Massim groups, or among groups from the Bismarcks and Solomons; there is no sharing of Papuan-related IBD segments between any mainland PNG or Massim group and any group from the Bismarcks or Solomons (Fig. 8C), which is remarkable. The extensive sharing of Austronesian-related IBD segments of 1-5 cM, which corresponds to a time of ~2.7 kya (*35*), is in keeping with the large impact of the Austronesian expansion, which spread rapidly across the lowland and island regions of Near Oceania shortly after its arrival around 3 kya (*6, 9, 17*). The distinct geographic pattern in the sharing of Papuan-related ancestry – which, to the best of our knowledge, has not been noted previously - must have a different explanation; it cannot reflect the spread of people with both Austronesian and Papuan ancestry as then there should not be a distinction between patterns of IBD sharing for Austronesian-related vs. Papuan-related segments. We speculate that the arrival of the Austronesians may have impacted the indigenous Papuan societies, resulting in enhanced movement of Papuans within these two geographic regions (mainland/Massim, and Bismarck/Solomon Archipelagos). However, IBD sharing of segments of 1 to 5 cM reflects a time span of a few thousand years, and so the results in Fig. 8C could also reflect movement of people prior to the arrival of the Austronesians, or movement unrelated to the arrival of the Austronesians (*28*). For example, the flooding of the shallow continental shelf east of New Guinea, following the Last Glacial Maximum, appears to have led to depopulation of the current islands of the Massim region, with subsequent recolonization after ~5 kya (*19*); perhaps the IBD-sharing results reflect these movements. Finally, we found increased sharing of IBD segments between Massim groups practicing Kula vs. Massim groups that do not participate in this specific trading exchange ritual in the Massim (Fig. 9; figs S15 and S16). Kula, made famous by Malinowski in his classic work *Argonauts of the Western Pacific* (1922), is an example of “... gift exchange with delayed reciprocity. Two kinds of objects are passed between a chain of partners in a large maritime region (the ‘Massim’), providing strong networks of support, a competitive element between the participants and the thrill of adventure” (*15*). The objects in question consist of decorated shell armbands (*mwali*) and decorated shell necklaces (*bagi* or *souvlava*) which travel in opposite directions through the participating islands (*12, 15*); long-distance travel between islands is largely restricted to Kula-related voyages, leading to the expectation that Kula would have an impact on patterns of gene flow between islands. Indeed, a previous study of mtDNA and Y chromosome variation in the same samples studied here found less genetic structure between islands for the Y chromosome than for mtDNA, suggesting a potential impact of male-mediated Kula voyages (*26*). The increased sharing of IBD segments for Kula vs. non-Kula groups in the Massim is further evidence for the impact of this cultural practice on patterns of gene flow. However, we note that the increased sharing of IBD segments for Kula vs. non-Kula groups occurs across all size classes of IBD segments (Fig. 9), and thus has an associated time depth of a few thousand years (*33, 35*), whereas archaeological evidence suggests a time depth for Kula of ~500 years (*14, 17*), as well as the existence of other inter-island exchange networks that preceded the formation of Kula (*17, 36*) and from which Kula may have developed. Thus, both archaeological and genetic evidence suggest that Kula should be viewed as arising out of a previous history of enhanced contact among the islands involved, rather than necessarily introducing novel avenues of contact that did not exist before. As expected from the nature of IBD sharing (*33, 35*), we observed a decline in the relative amount of IBD sharing with longer IBD segments (Fig. 9; figs S15 and S16); still, we identified some strong sharing within northern and southern Massim, suggesting extensive regional interactions in the last few hundred years.

In conclusion, our results concerning the Massim region of Papua New Guinea fill an important lacuna in genetic studies of Near Oceania. Austronesian-related ancestry varies across the region but overall is in the minority, even though Austronesian languages are in the majority. We demonstrate the existence of a novel Papuan-related ancestry that is associated with Rossel Island and probably arose as a consequence of its isolation. In addition to the expected signal of a rapid and widespread dispersion of Austronesian-related ancestry across coastal and island Near Oceania, we also found an unexpected signal of substantial movement of people with Papuan-related ancestry that was geographically restricted, occurring exclusively between mainland PNG and the Massim region, and between the Bismarck and Solomon Archipelagoes. We speculate that these movements of people with Papuan-related ancestry may reflect a disruptive impact of the arrival of Austronesian people. Finally, we document the effect of a cultural trait, the Kula, on relative amounts of IBD sharing among participating vs. non-participating groups; the Kula thus joins other examples of cultural traits, such as residence pattern (*37, 38*) and social stratification (*39*), that can influence human genetic diversity.

## Materials and Methods

### Sample and data information

Saliva samples were collected in 2001 with the approval of the Medical Board of PNG and with support from the Diocese of Alotau, PNG (Missionaries of the Sacred Heart, M.S.C.), particularly Fr Joe Ensing (M.S.C.) and then Bishop Desmond Charles Moore (M.S.C.). Written informed consent was obtained from each donor, after the project was explained and all questions answered to the satisfaction of the donor. Genetic work within this study was additional approved by the Ethics Commission of the University of Leipzig Medical Faculty. MtDNA and Y chromosome data were published in a previous study (*26*). Here, we generated genome-wide data (~1.6 million SNPs) for 255 (192 from Massim, 33 from Gulf Province, and 30 from Central Province) individuals on the Illumina Infinium Multi-Ethnic Global Array (MEGA); genotyping was carried out by the Laboratorio de Servicios Genómicos at the Laboratorio Nacional de Genómica para la Biodiversidad, Irapuato, Mexico on our request. We first merged our newly generated data with published data from Papuan populations on the same SNP array (*27*), and then with published whole-genome sequencing (WGS) data from Oceanians, East Asians, Europeans, and Africans (*30, 40, 41*). For the WGS data, we obtained the jointly re-called genotypes from Choin et al. (*41*). To avoid batch effects between the array and WGS data, we extracted the overlapping sites (including both monomorphic and polymorphic sites) and removed sites whose reference vs. alternative alleles were inconsistent between the array and WGS data. For sites with more than two alleles, we first flipped the WGS data and then removed those flipped sites that still had more than two alleles. Merging was done using PLINK v1.9 (*42*). For quality control, we first excluded sites with more than 5% missing data in the entire dataset and sites with more than 50% missing data and/or Hardy-Weinberg equilibrium p values below 0.00005 within a population group (except for groups with only one individual). Then, we removed individuals with more than 5% missing data or with parents speaking different languages or coming from different locations. We also filtered out individuals to exclude up to 1^st^ degree kinship pairs. Data missingness and Hardy-Weinberg equilibrium were calculated using PLINK v1.9 while the individuals to be removed to avoid 1^st^ degree relatedness were inferred using KING (*43*), as implemented in PLINK v2 (*44*). There are 776 individuals and 1,408,767 SNPs remaining after quality control (table S1 indicates the individuals that were removed and why).

The sampled Massim individuals were assigned to one of 14 groups according to parental geographic origins as described in the previous study (*26*): (1) Trobriand; (2) Gawa; (3) Woodlark (also known as Muyuw); (4) Laughlan (also known as Budibudi); (5) Fergusson (also known as Moratau), including a few individuals from nearby Dobu (also known as Watoa) and Goodenough (also known as Nidula); (6) Normanby (also known as Duau); (7) Milne Bay mainland eastern tip (Mainland eastern tip); (8) Misima, including some individuals from nearby Paneati, Panapompom and Kimuta; (9) Western Calvados (including: Motorina, Bagaman, Utian or Brooker Island, and Panaumala); (10) Eastern Calvados (including: Dadahai, Kuanak or Abaga Gaheia Island, Nimoa, Panatinane or Joannet Island, Panawina, Sabarl, and Wanim or Grass Island); (11) Sudest (also known as Vanatinai or Tagula); (12) Rossel (also known as Yela); (13) Wanigela (and nearby settlements); and (14) Airara (and nearby settlements). Together with a previously-studied group from the region (Northern, from North Collingwood Bay) from Bergström et al. (*27*), there are data from 15 groups over the entire Massim region including mainland and island parts. Sub-region groups were defined according to geography as in the previous study (*26*): Collingwood Bay (Northern, Wanigela, Airara), western Massim (Mainland eastern tip, Normanby, Fergusson), northern Massim (Trobriand, Gawa, Woodlark, Laughlan), and southern Massim (Misima, Western Calvados, Eastern Calvados, Sudest, Rossel). The locations of the newly-studied and the reference groups are shown in Fig. 1 (and fig. S1).

### Population structure analyses

For population structure analyses, variants were pruned beforehand for linkage disequilibrium using PLINK v1.9, excluding one variant from pairs with r2 > 0.4 within windows of 200 variants and a step size of 25 variants. Principal component analysis (PCA) was done with smartpca v16000 (*45*). Two individuals (papuan6278328 and papuan6278360) from Bergström et al. (*27*) were identified as PCA outliers and excluded from this study. We then ran ADMIXTURE v1.3.0 (*46*) for K = 2 to K = 15 with 100 replicates for each K with random seeds. We used pong v1.4.7 (*47*) to visualize the 20 ADMIXTURE replicates with the highest likelihoods for the major mode at each K. We also plotted K = 2 (to visualize the variation of East Asian- and Papuan-related ancestries) and K = 8 (the K with the lowest cross-validation error; fig. S6) on a map in R v4.0.3.

### F3 and F4 statistics

We used admixr v0.9.1 (*48*) from ADMIXTOOLS v7.0.2 (*49*) to compute f3- and f4-statistics, with significance assessed through block jackknife resampling across the genome. For f3- and f4-statistics, the African groups Mbuti and Yoruba were used together as the outgroup.

### Data phasing

We used consHap (*50*) to obtain a consensus of phasing from the results of SHAPIT v2 (*51*), BEAGLE v5.1 (*52*), and EAGLE v2 (*53*) using the genetic map from the 1000 Genomes Project Phase3 (*54*). In general, phasing accuracy can be increased by increasing the number of iterations and conditioning states on which haplotype estimation is based (*55*). We therefore ran SHAPIT v2 with options --burn 10, --prune 10 and --main 30 for iteration number with 500 conditioning states, leaving other parameters as default; BEAGLE v5.1 with options burnin=12, iterations=24, phase-states=24, and ne=800000 (this is smaller than the default value, but is recommended by the authors for populations with a smaller effective population size, as expected for island populations in Oceania); EAGLE v2 with --Kpbwt 40000 (which determines the number of conditioning haplotypes).

### Identity by descent (IBD) analyses

We identified shared IBD blocks between each pair of individuals and homozygous-by-descent (HBD; same as runs of homozygosity, ROH) blocks within each individual using refinedIBD (*56*). Both identified IBD and HBD blocks are considered as IBD blocks in our analyses, which is analogous to pairwise shared coalescence (PSC) segments in a previous study (*35*). The IBD blocks within a 0.6 cM gap were merged using the program merge-ibd-segments from BEAGLE utilities (https://faculty.washington.edu/browning/refined-ibd), allowing only 1 inconsistent genotype between the gap and block regions. We used IBD blocks at least 2 cM in length shared by individuals within a population to investigate the demography of each population group. Then, we used IBD blocks in 1-5 cM, 5-10 cM, and over 10 cM to investigate the sharing between individuals from different populations for different time periods (*33, 35*). For network visualization of the sharing between populations, the pairs with average sharing ≥ 0.5 (i.e., on average at least half of the pairs share IBD blocks) were kept to reduce noise and false positives. We summarize the patterns of shared IBD length within a population by averaging over all comparisons between individuals, i.e. we define the average of summed IBD length L as

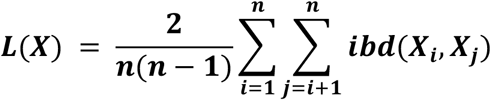

here, n is the number of individuals in population X and ibd(X_i_, X_j_) is the length of IBD shared between individuals X_i_, and X_j_. For the number of blocks, an analogous equation applies. Similarly, for sharing between two populations X, Y we define the average of summed IBD length L as

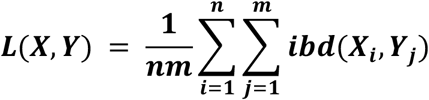

where m is the number of individuals in population Y. In order to compare IBD sharing between Massim groups (Kula vs non-Kula), we use a statistic motivated by FST (*57*). In particular, we define the similarity statistic S as

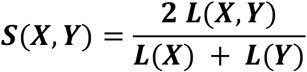

This statistic will be zero if there is no IBD sharing between X and Y (since L(X,Y) is zero), and it will be one if IBD sharing is independent of population structure, i.e L(X) = L(Y) = L(X,Y). For our analysis, we compute a matrix of pairwise S-statistics for all pairs of populations. In order to test whether there is more recent gene flow within the Kula/non-Kula group relative to the background, we use the relative similarity statistic R as

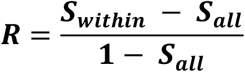

where S_within_ and S_all_ are the average pairwise S-statistics within Kula/non-Kula populations and all Oceanian populations, respectively. If the IBD-sharing among the Kula/non-Kula groups and among all Oceanians were equal then R will be zero, and the more shared migration the Kula/non-Kula groups have, the larger R will be. We calculate the standard error of the R-statistic by jackknife resampling the chromosomes. The significance of the R-statistic between Kula-vs. non-Kula groups was estimated by comparing the observed value to the distribution of simulated values generated by randomly assigning Massim groups to either Kula- or non-Kula groups and calculating the difference in relative IBD similarity. Scripts made for analyzing IBD results are available from https://github.com/dangliu/Massim_project.

### ChromoPainter and GLOBETROTTER analyses

To study haplotyple sharing, ChromoPainter v2 (*58*) was run on the phased dataset with sample sizes for each group randomly down-sampled to 5 (all individuals were used for groups with size < 5). We began with 10 iterations of the EM (expectation maximization) process to estimate the switch rate and global mutation probability, using chromosomes 1, 5, 10, 15, and 20. With the estimated switch and global mutation rates, we ran the chromosomal painting process for all chromosomes, which then gave the output for downstream analyses. We first attempted to paint the Massim chromosomes using them only as recipients and all of the other individuals as both donors and recipients. The EM estimation of switch rate and global mutation probability were ~142.27 and ~0.0002, respectively, which were then used as the starting values for these parameters for all donors in the painting process. To investigate differential affinities of Massim groups to PNG highlanders vs. Rossel, we also performed another run using the Massim (except for Rossel) only as recipients and all of the other individuals (including Rossel) as both donors and recipients. The EM estimation of switch rate and global mutation probability for this analysis were ~142.21 and ~0.0002, respectively.

To investigate the admixture of Austronesian- and Papuan-related ancestries, GLOBETROTTER (*59*) was run on the ChromoPainter output with all Massim as recipients and East Asians (including both Austronesian and non-Austronesians to increase power) and Papuan Highlanders (who showed no more than 0.0001 East Asian-related ancestry in the ADMIXTURE results for K=2) as surrogates. We first tested the certainty and potential waves of admixture events, and then estimated the major and minor sources as well as the dates of admixture. The Rossel group showed an unclear signal for admixture inference, while a single pulse of admixture was inferred for the other groups. The distributions of admixture dates were estimated via 100 bootstraps.

### ALDER admixture dating

We used ALDER v1.03 (*60*) with default settings, using East Asian Austronesian and Southern Highland groups as reference sources, to date the time of admixture of Austronesian and Papuan ancestries in Massim groups.

### Local ancestry inference and ancestry-specific analyses

We ran the PopPhased mode of RFMix v1.5.4 (*61*) to infer local ancestry across the genome for each individual in our dataset with options -e 3 (three EM iterations), -G 85 (85 generations since the admixture event; as suggested in a previous study (*27*)), -n 5 (minimum five reference haplotypes per tree node; as suggested by the authors of RFMix), --use-reference-panels-in-EM (the reference samples were used in EM iterations), and the East Asians (including both Austronesian and non-Austronesians to increase power) and Papuan highlanders (who showed no more than 0.0001 East Asian-related ancestry in the ADMIXTURE results for K=2) as reference panels. The Papuan highlanders were randomly down-sampled to 50 individuals, to be the same as the sample size of East Asians. We used 0.95 as the cutoff of forward-backward probability (i.e. the posterior probability of an inferred ancestry at a SNP in a haplotype). We interpreted the inferred East Asian-related segments as Austronesian-related segments, as indicated in f4 results (fig. S8). Global ancestry for each Massim group was calculated from the local ancestry profiles of the corresponding individuals. Papuan-/ Austronesian-related ancestry-specific PCA was performed with PCAmask (*62*) on East Asian and Oceanian individuals with at least 33% Papuan-/ Austronesian-related ancestry based on the ADMIXTURE results for K=2. The same local ancestry-inferred results were applied to IBD results to study Papuan-/ Austronesian-related ancestry-specific IBD. Analyses of local ancestry inference, global ancestry calculation, and ancestry-specific PCA were done using a pipeline from a previous study (*63*); scripts for preparing input files for these analyses were modified from the pipeline (available from https://github.com/dangliu/Massim_project). Ancestry-specific IBD analyses were carried out by following the pipeline described in a previous study (*64*).

## Acknowledgments

We thank: all sample donors for making this work possible; Joe Ensing (M.S.C.) for valuable assistance in sample collection; Roland Schröder for technical assistance; Benjamin Vernot and Lluis Quintana-Murci for data access; Janet Kelso, Sandra Oliveira and Irina Pugach for helpful advice concerning computational analyses; and Ben Shaw for helpful discussion of the archeological information concerning the Massim and the Kula tradition. The field work received valuable support by the Missionaries of the Sacred Heart (M.S.C.) and the Diocese of Alotau-Sideia via several Fathers serving on the different islands, the St Paul’s Pastoral Centre Hagita, and the then Bishop Desmond Charles Moore (M.S.C.).

## Author contributions

Conceptualization: MK, MS

Sample collection: MK, WS

Data analysis: DL

Development of IBD similarity statistics: BMP

Supervision: MS

Writing—original draft: DL, MS

Writing—review & editing: DL, BMP, WS, MK, MS

## Data and materials availability

The genome-wide SNP array data generated in this study is available from European Genome-Phenome Archive (EGA; https://ega-archive.org), under accession code EGASXXXXXXXXXXX. Scripts written/modified for ancestry specific analyses and IBD analyses are available from https://github.com/dangliu/Massim_project.

## Supplementary Materials

Please see the Supplementary Materials file (Kula.ms.bioRxiv.SM.v1.pdf).

