## Supplementary Materials for "Assessing human genome-wide variation in the Massim region of Papua New Guinea and implications for the Kula trading tradition"

**This PDF file includes:**

Figs. S1 to S16
Tables S1 to S2

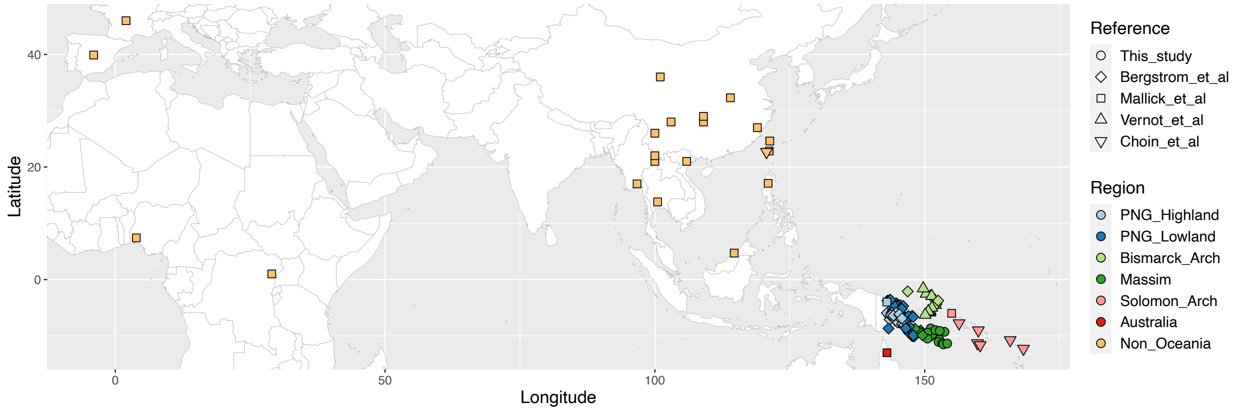


**Fig. S1.**
**Map showing the location of the studied groups.**

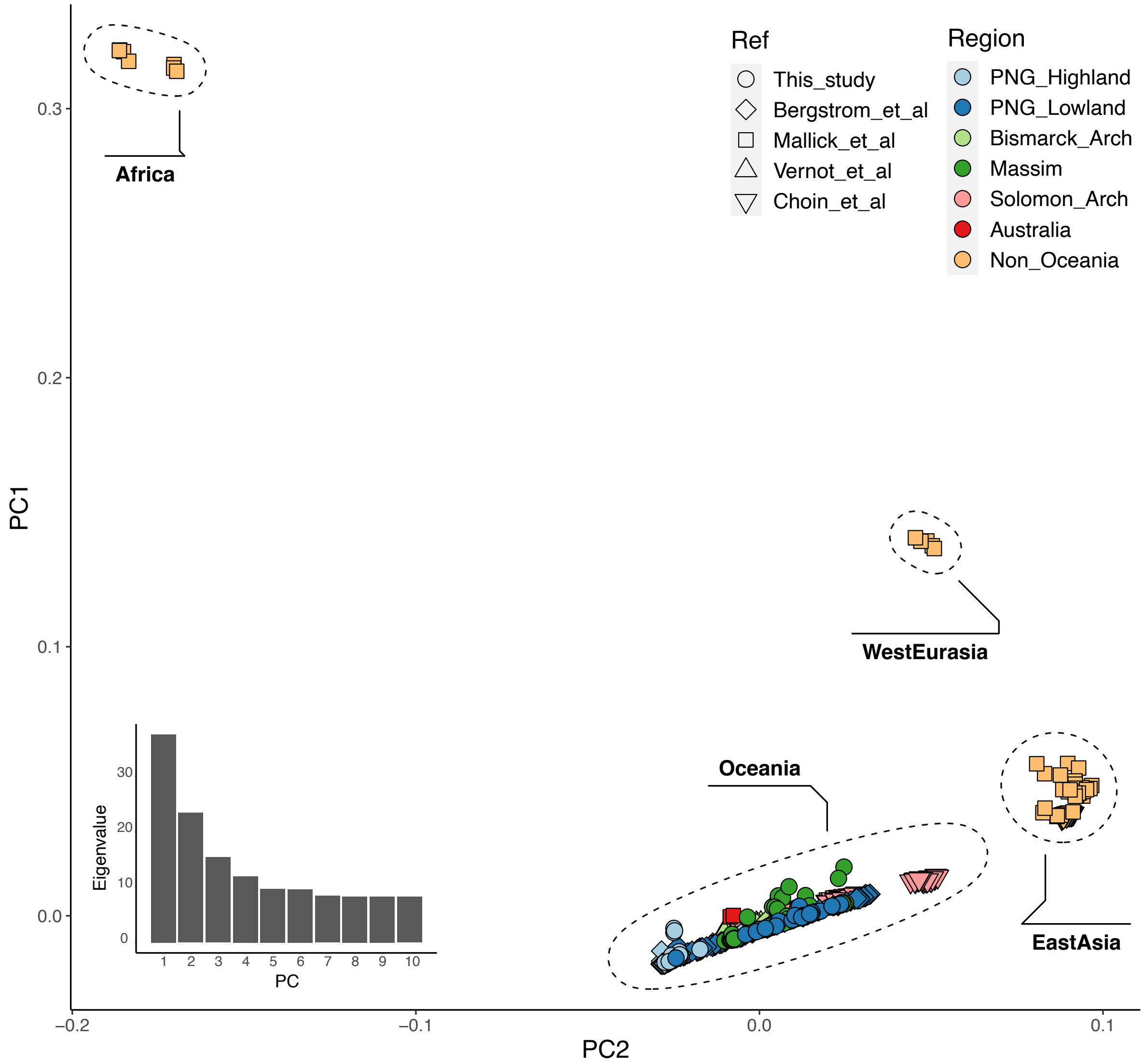


**Fig. S2.**
**Principal component analyses (PCA) plot of PC1 vs. PC2 for all individuals in the dataset**. Symbol colors indicate region and shapes indicate the corresponding publication. The eigenvalues from PC1 to PC10 are shown on the bottom left.

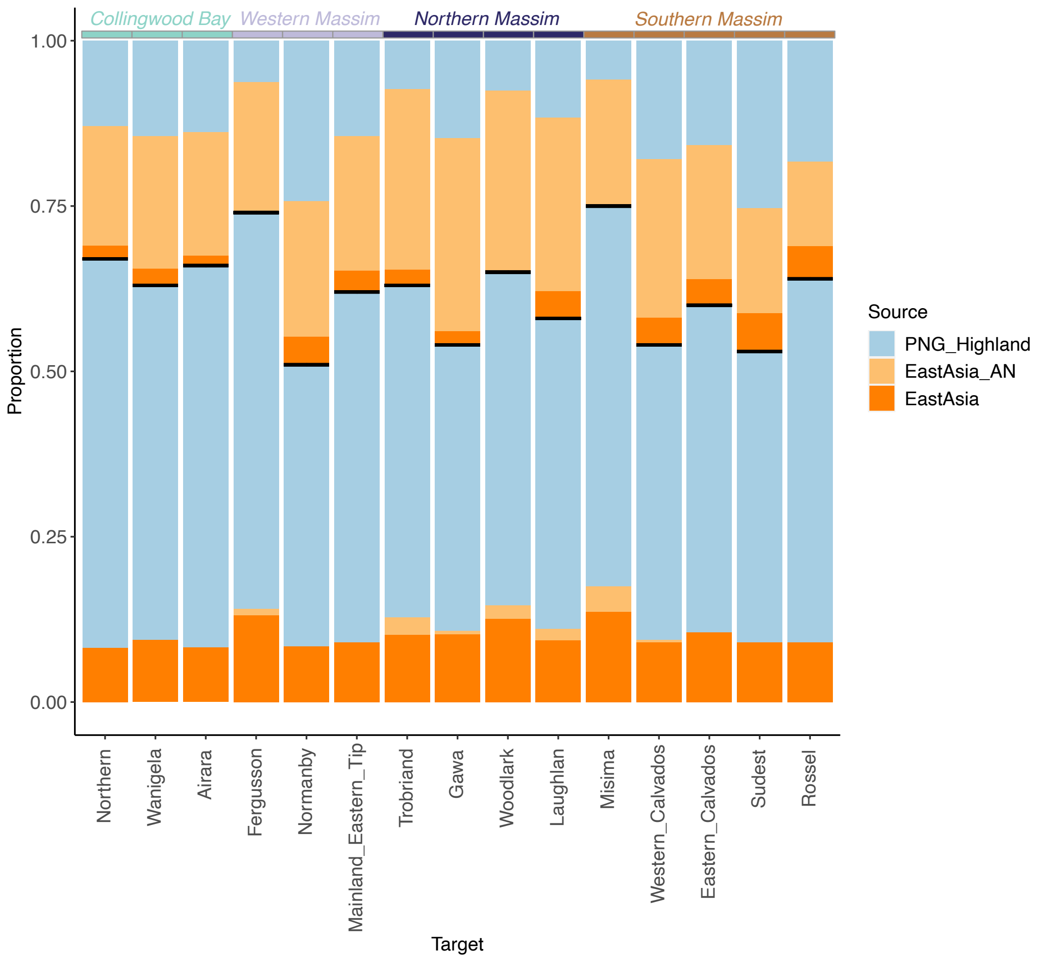


**Fig. S3.**

**Admixture sources inferred by GLOBETROTTER.** The middle horizontal black line in each bar separates the minor source (top) from the major source (bottom).

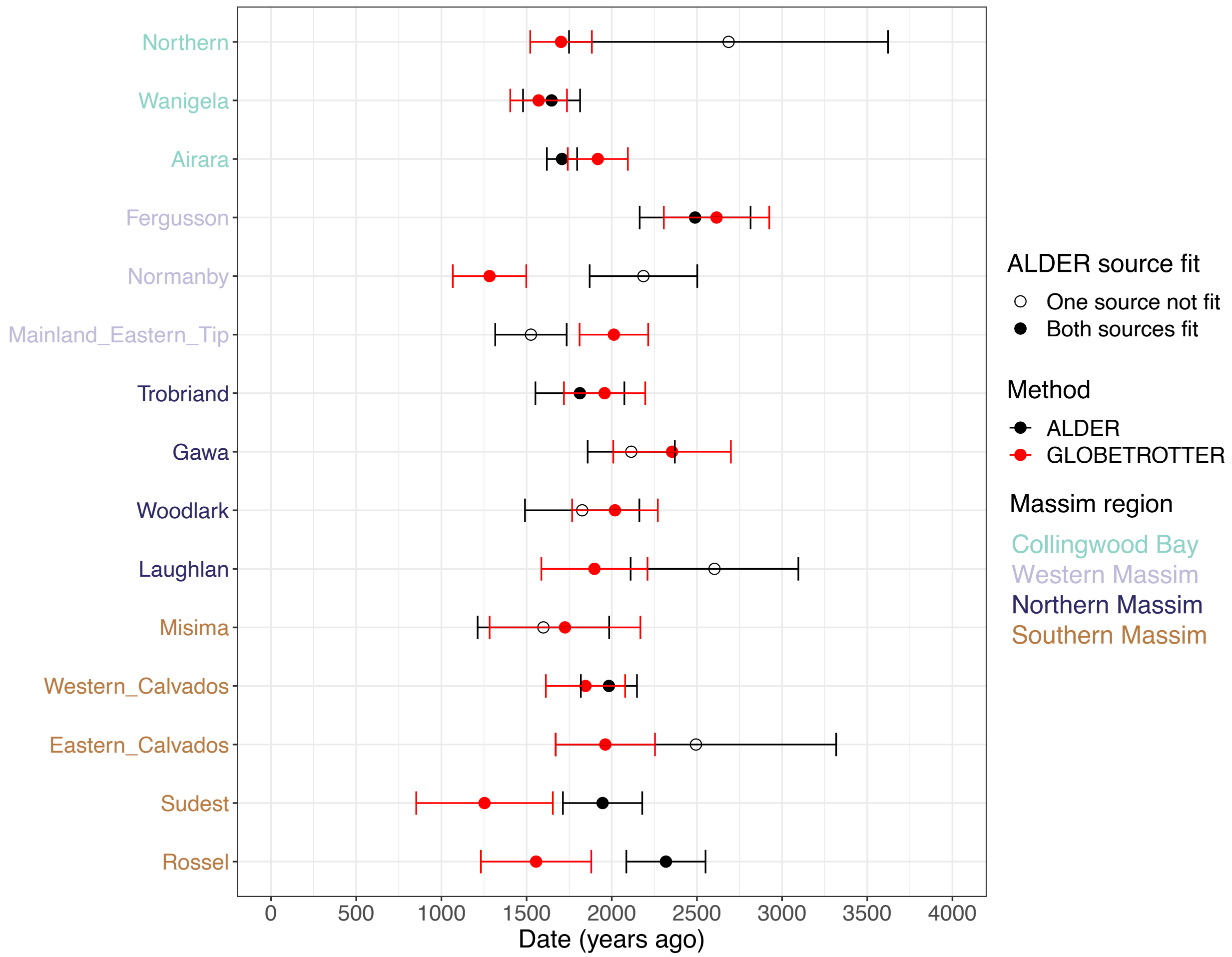


**Fig. S4.**

**Admixture dates inferred by GLOBETROTTER and ALDER.** Massim group labels are colored according to Massim region. GLOBETROTTER results are in red while ALDER results are in black. ALDER was performed using East Asian Austronesians and Southern Highlanders as sources; an empty dot indicates that one/both source(s) might not be a good proxy for the admixed target group while a solid dot denotes both sources fit well. Error bars indicate +/- one standard error. Although the linkage disequilibrium (LD) decay curve of ALDER for all of the Massim groups fit a curve weighted by 2 reference sources, some groups did not fit a curve with one reference source. This suggests that either the East Asian Austronesian or the Southern Highland group might not be a good proxy for the admixture in some of the Massim groups.

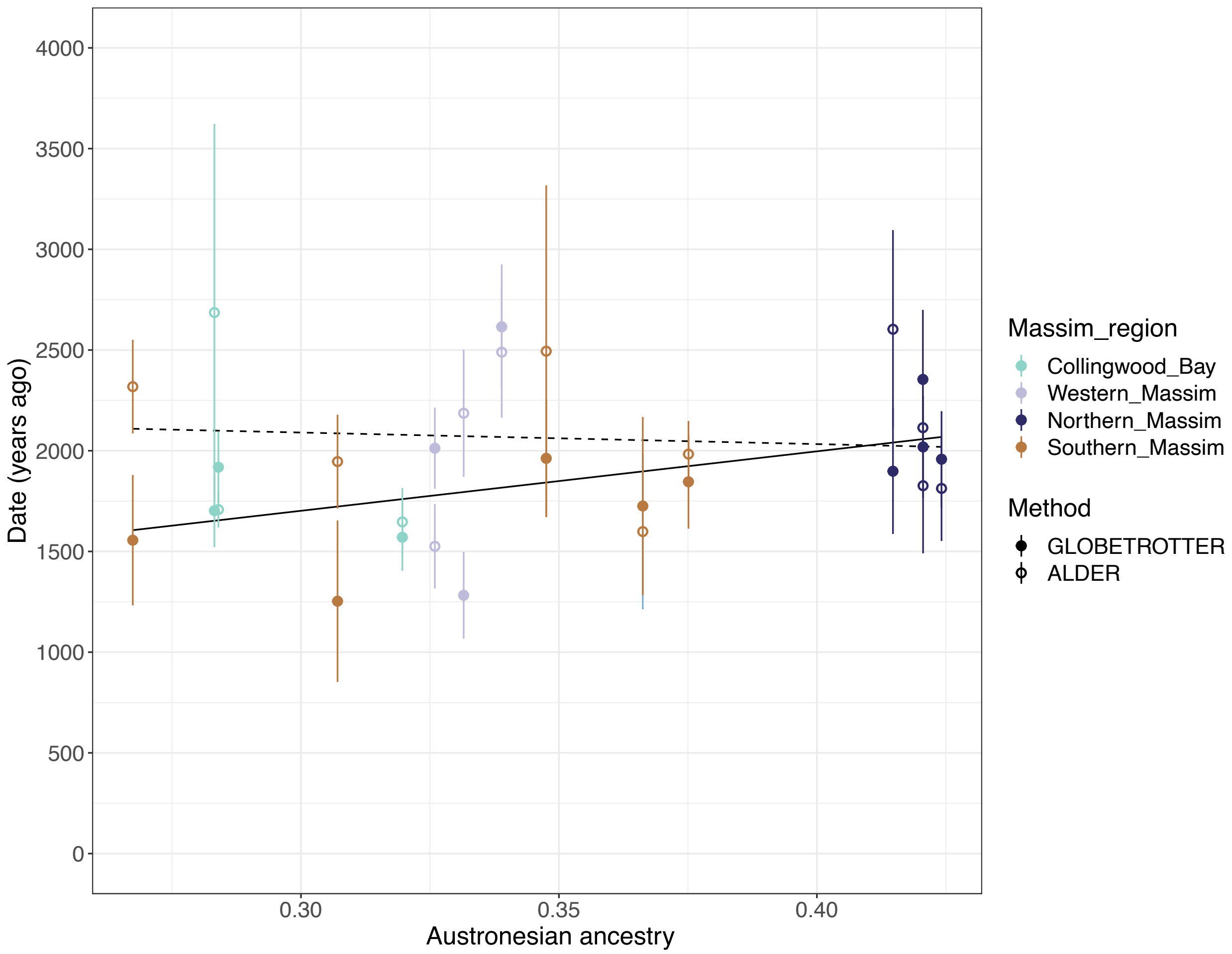


**Fig. S5.**

**Admixture dates vs. Austronesian ancestry for Massim groups.** A solid or empty point indicates that the admixture date was inferred by GLOBETROTTER or ALDER, respectively; points are colored according to Massim region. Error bar indicates +/- one standard error of the inferred admixture dates. A solid or dashed regression line was calculated for the GLOBETROTTER or ALDER points, respectively. The linear regression results for GLOBETROTTER points: $r^2$=0.193 and p=0.102; and for ADLER points: $r^2$=0.006 and p=0.778.

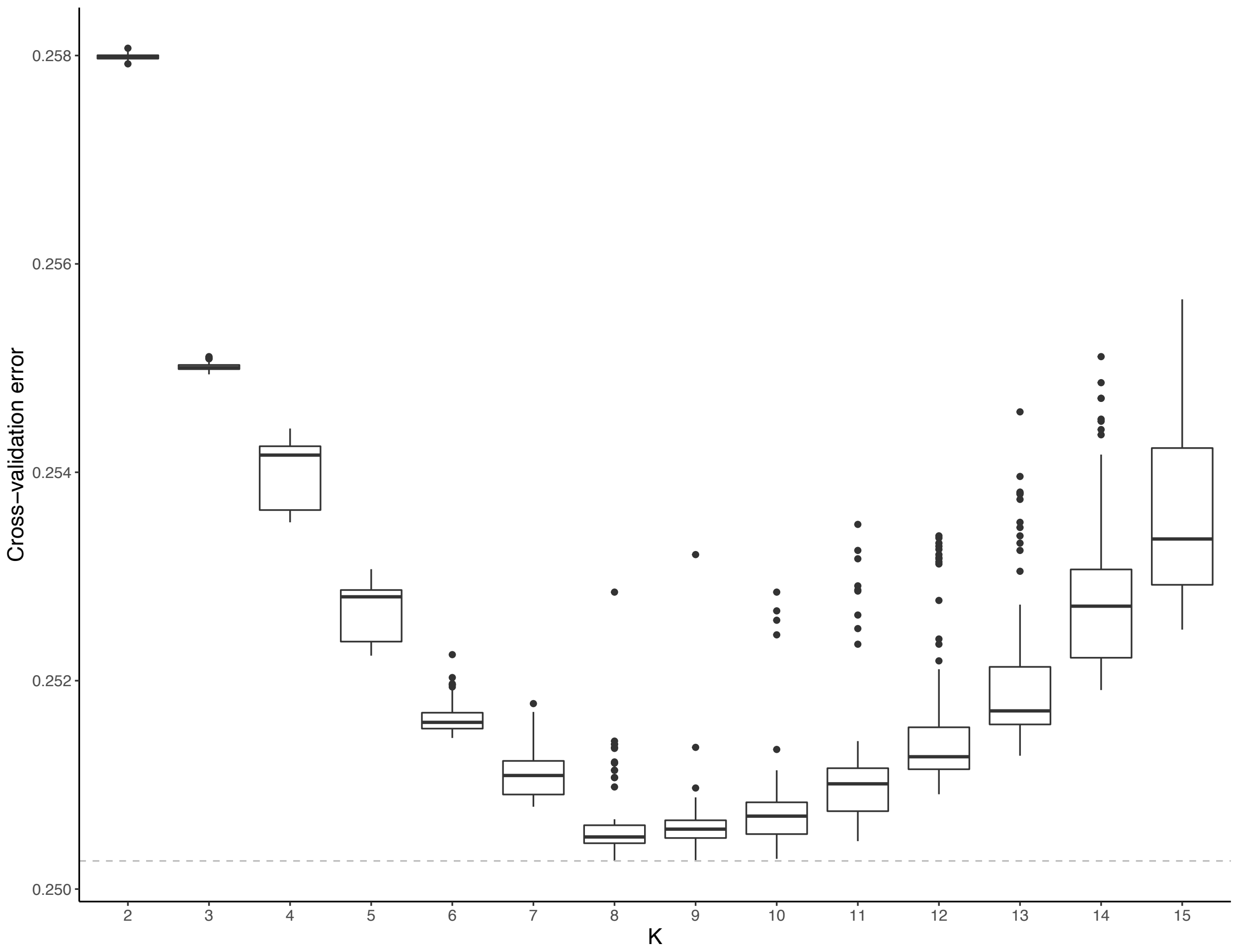


**Fig. S6.**
**Cross validation errors of ADMIXTURE runs for K= 2 to K = 15, based on 100 runs for each K value.**

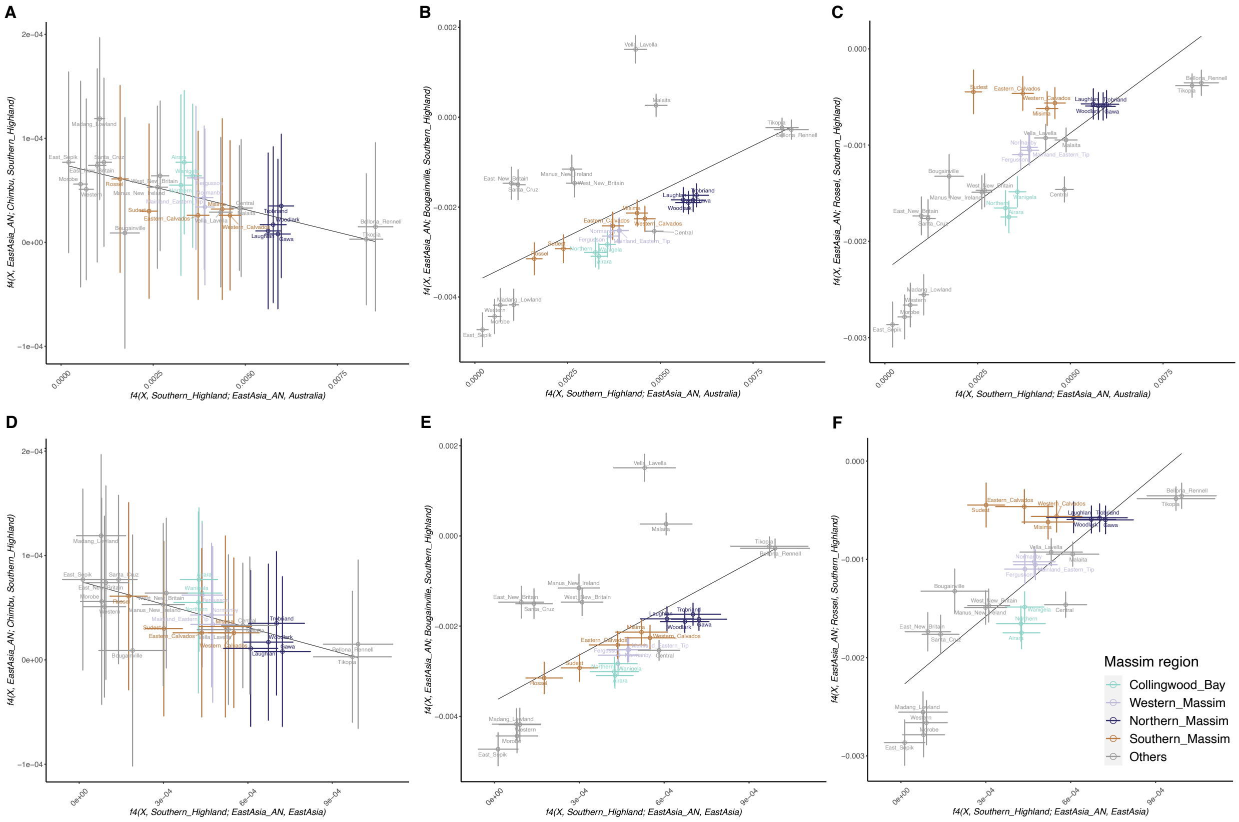


**Fig. S7.**

**F4 statistics measuring differential PNG ancestry affinities of Oceanian groups with respect to Austronesian ancestry affinity.** The value of f4(Oceanian groups, Southern province highlanders; East Asian Austronesians; (A)-(C) Australians/ (D)-(F) non-Austronesian East Asians) is on the x-axis, and the value of f4(Oceania groups, East Asian Austronesians; (A) and (D) Chimbu province highlanders/ (B) and (E) Bougainville/ (C) and (F) Rossel islanders; Southern province highlanders) is on the y-axis. X denotes the Oceanian groups, colored according to Massim region. Error bars indicate +/- three times standard error. Linear regression lines were computed using all point values shown on the plot.

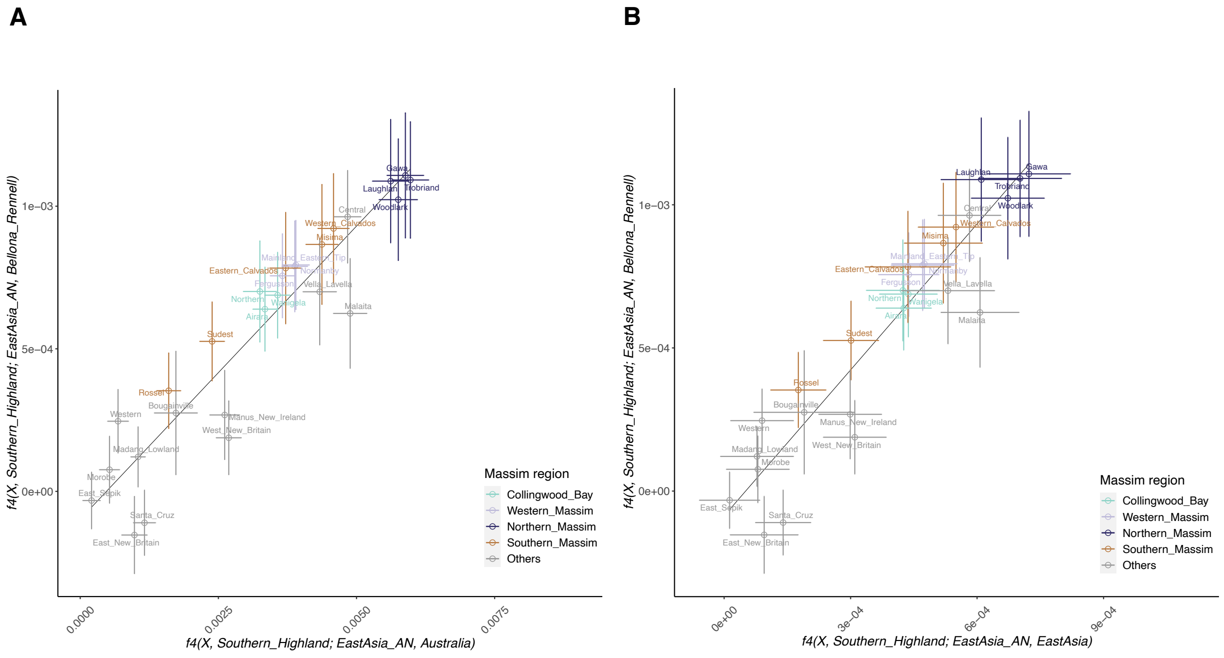


**Fig. S8.**

**F4 statistics measuring differential Austronesian ancestry affinities of Oceanian groups with respect to overall Austronesian ancestry affinity.** The value of f4(Oceanian groups, Southern province highlanders; East Asian Austronesians; (A) Australians / (B) non-Austronesian East Asians) is on the x-axis, and the value of f4(Oceanian groups, Southern province highlanders; East Asian Austronesians; Bellona/Rennell) is on the y-axis. X denotes the Oceanian groups, colored according to Massim region. Error bars indicate +/- three times standard error. Linear regression lines were computed using all point values shown on the plot.

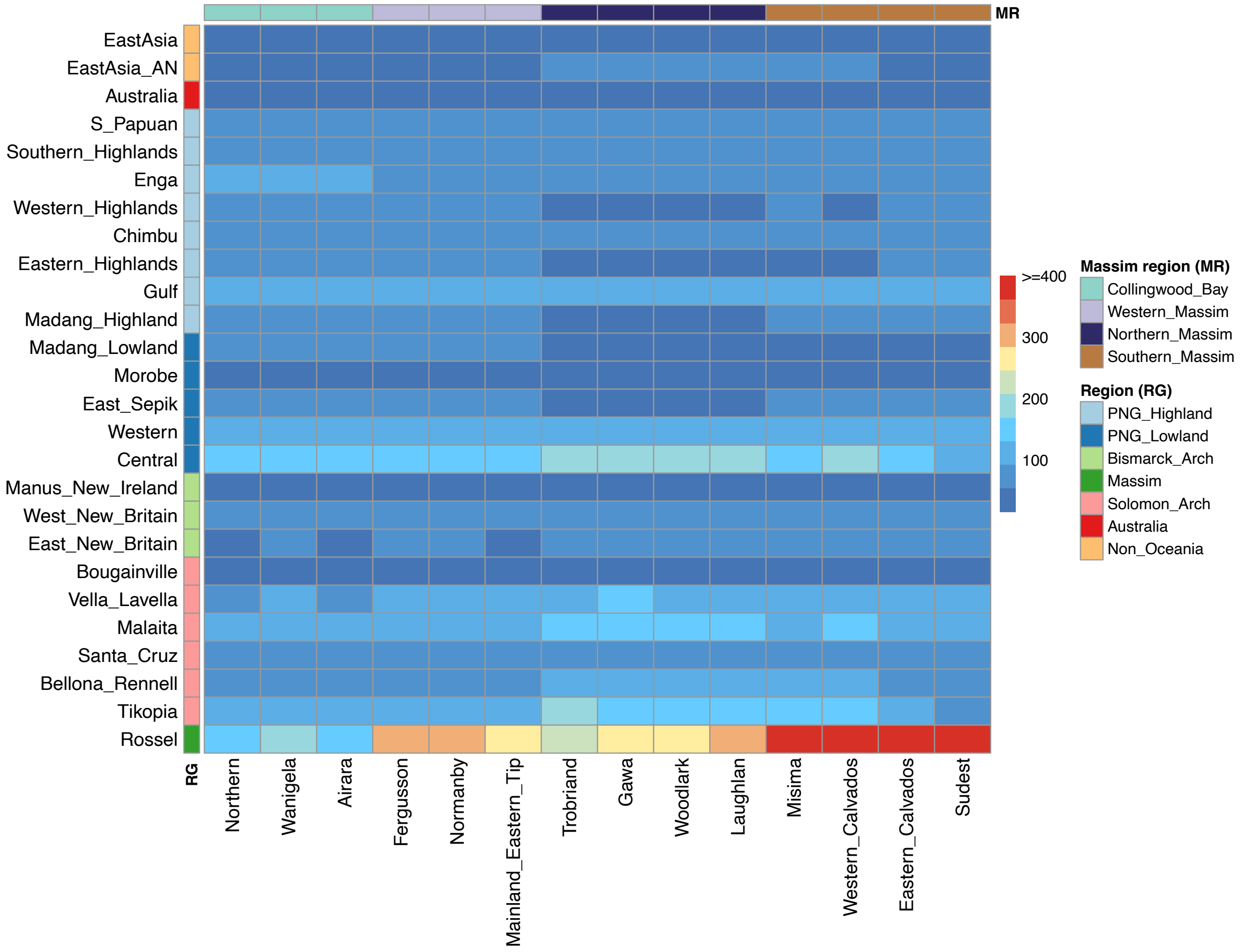


**Fig. S9.**

**ChromoPainter profile with Massim groups used only as recipients (except for Rossel, which was used as both a recipient and a donor).** The heatmap is proportional to the total copied length (cM) of a recipient from a donor. Color bars at the left and top of the heatmap indicate the region (RG) and Massim subregion (MR), respectively.

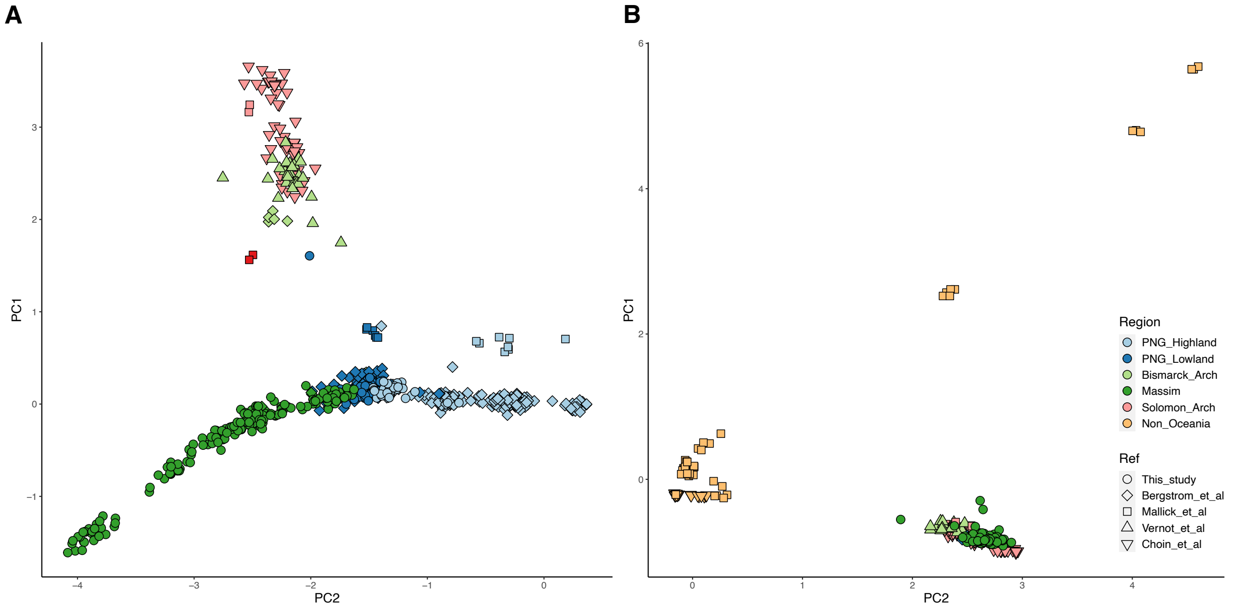


**Fig. S10.**

**Papuan/Austronesian ancestry-specific PCA of Oceanian (and East Asian) individuals.** Plot of PC1 vs. PC2 for all individuals from Oceanian (and East Asian) groups and using only (A) Papuan or (B) Austronesian segments identified by RFMix, colored according to region; Massim regions are further highlighted.

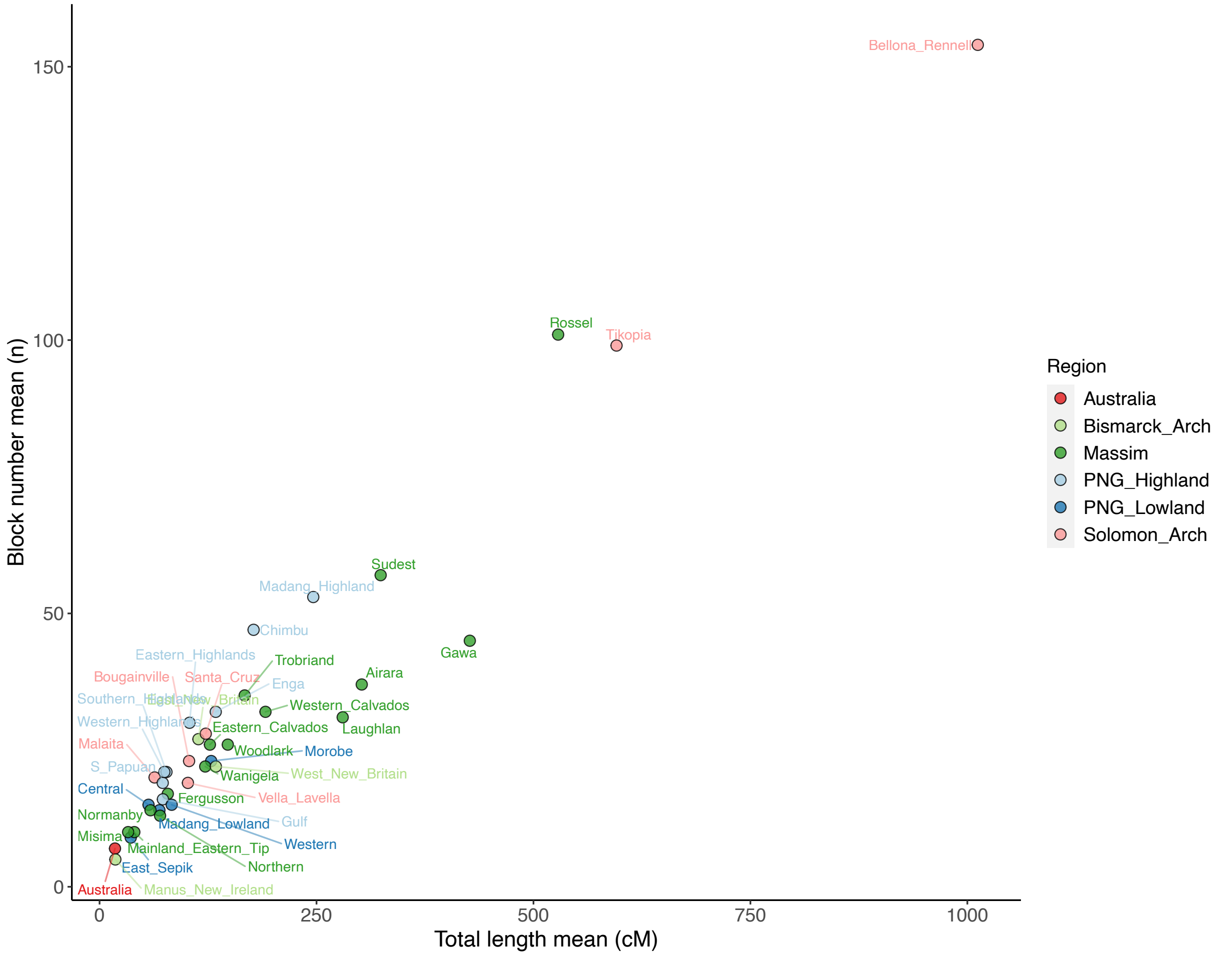


**Fig. S11.**

**IBD sharing within groups.** The groups are colored according to region.

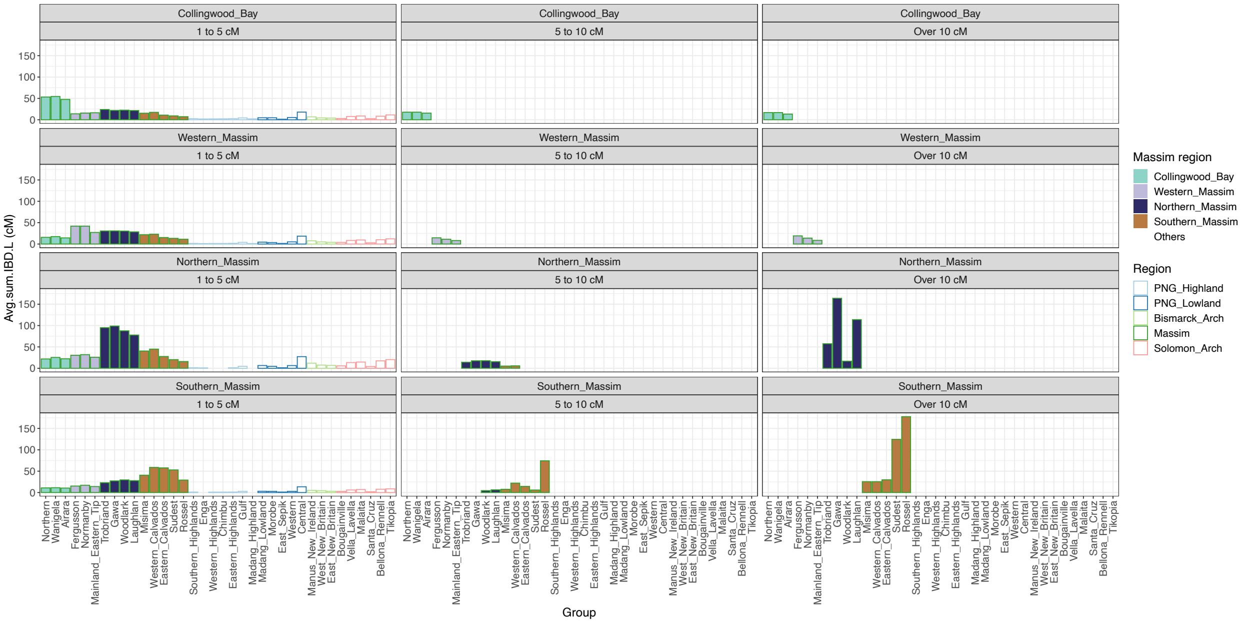


**Fig. S12.**

**Quantification of all IBD sharing between Massim region groups and Oceanian groups.** The average summed IBD length between Massim regional groups (in rows) and all Oceanian groups, for different size ranges (in columns), is depicted in the bar plots; filled bars are colored according to Massim region while the outline color of the empty bars indicates the Oceania region.

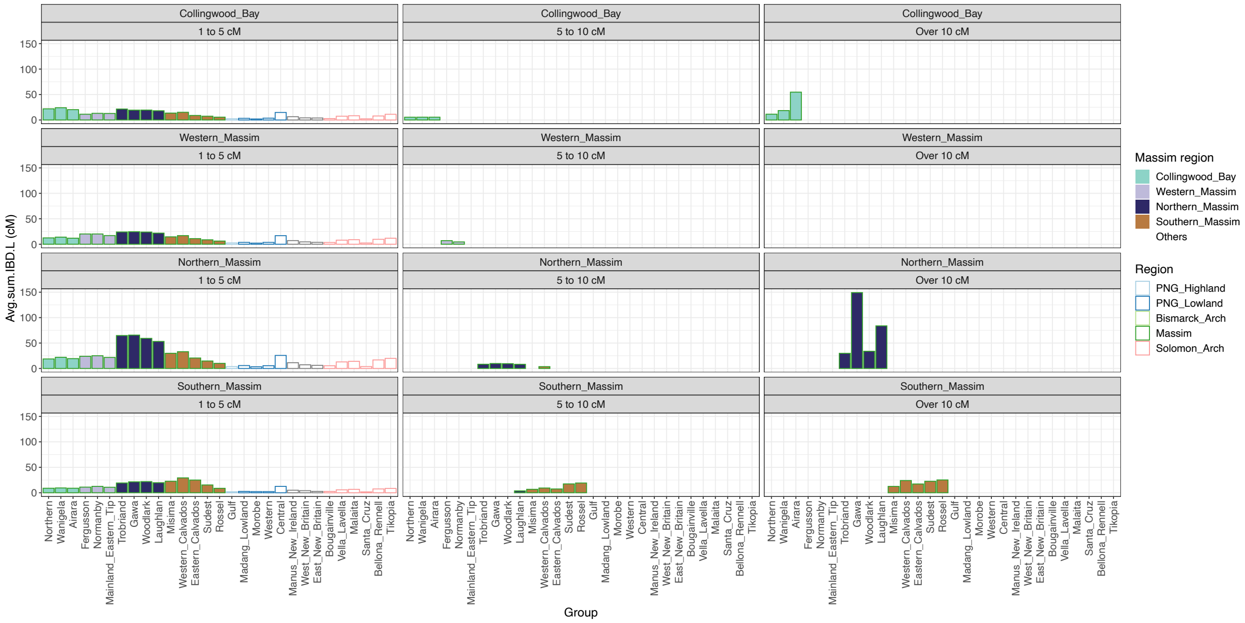


**Fig. S13.**

**Quantification of Austronesian ancestry-specific IBD sharing between Massim region groups and Oceanian groups.** The average summed Austronesian ancestry-specific IBD length between Massim regional groups (in rows) and all Oceanian groups, for different size ranges (in columns), is depicted in the bar plots; filled bars are colored according to Massim region while the outline color of the empty bars indicates the Oceania region.

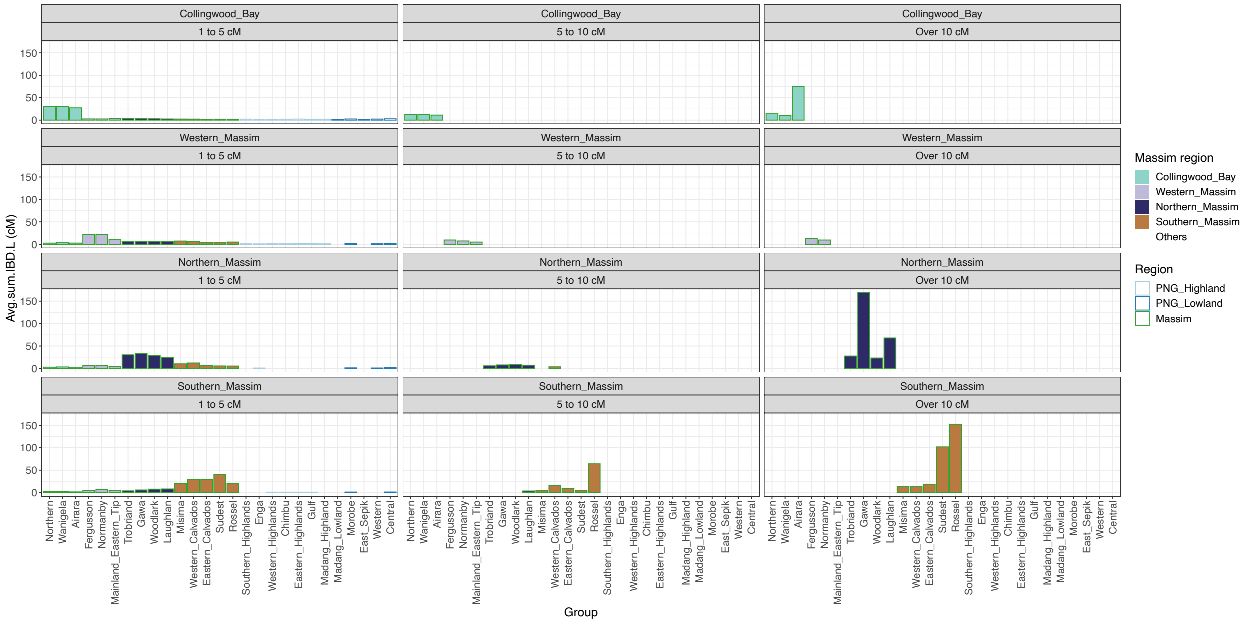


**Fig. S14.**

**Quantification of Papuan ancestry-specific IBD sharing between Massim region groups and Oceanian groups.** The average summed Papuan ancestry-specific IBD length between Massim regional groups (in rows) and all Oceanian groups, for different size ranges (in columns), is depicted in the bar plots; filled bars are colored according to Massim region while the outline color of the empty bars indicates the Oceania region.

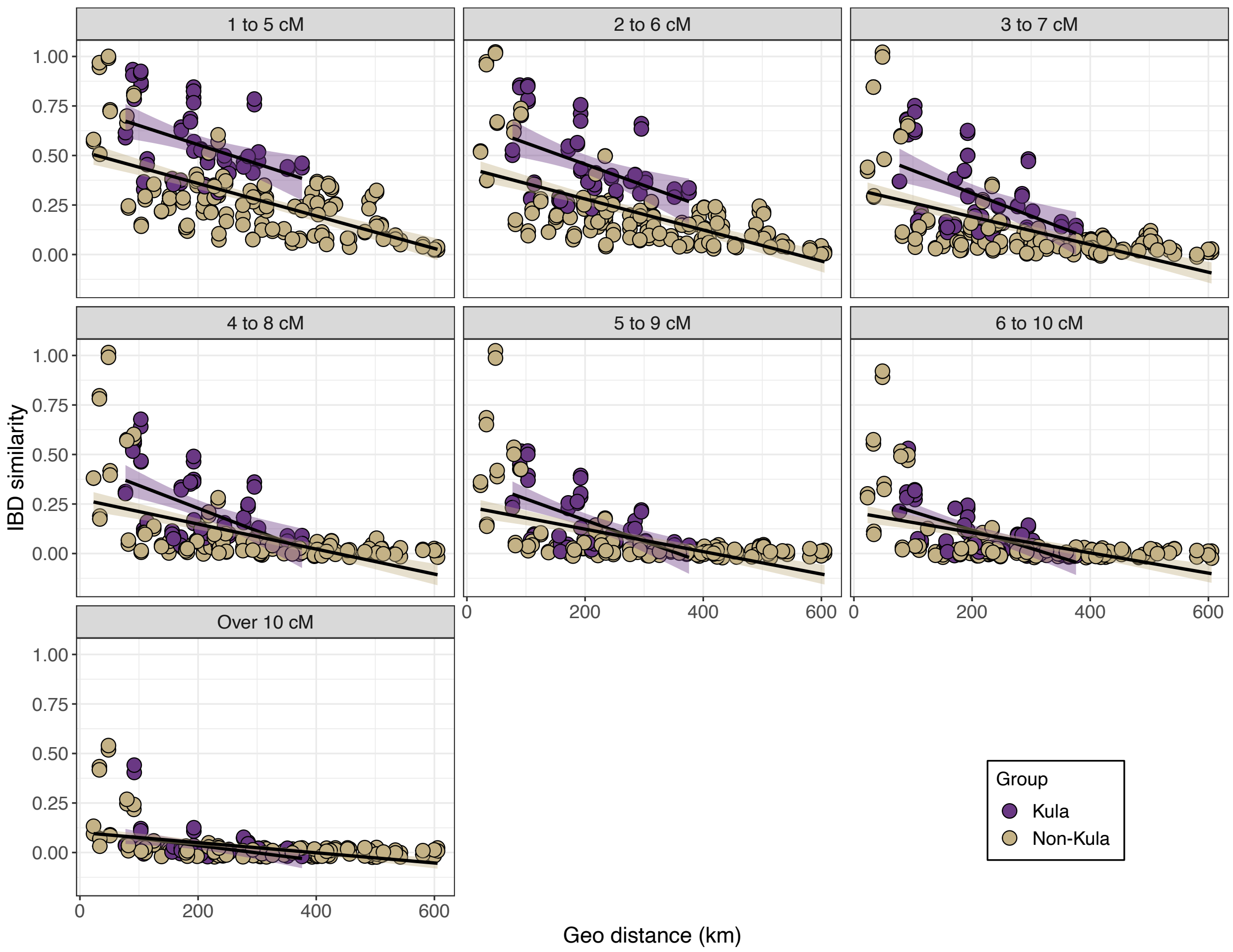


**Fig. S15.**

**IBD similarities of Kula- vs. non-Kula groups through space and time. Each point represents the IBD similarity calculated between a pair of Kula (purple) or non-Kula- (brown) groups.** Points with the same value were slightly jittered for visibility. The geographic distance (km) between pairs of groups is shown on the x-axis. Two regression lines with 95% confidence interval were separately calculated for the Kula or non-Kula points. Results calculated from different IBD size intervals are shown in different panels. These intervals (from 1-5 to over 10 cM) correspond to ~2.7, ~1.5, ~1.1, ~0.8, ~0.7, ~0.6, and ~0.2 thousand year ago (kya).

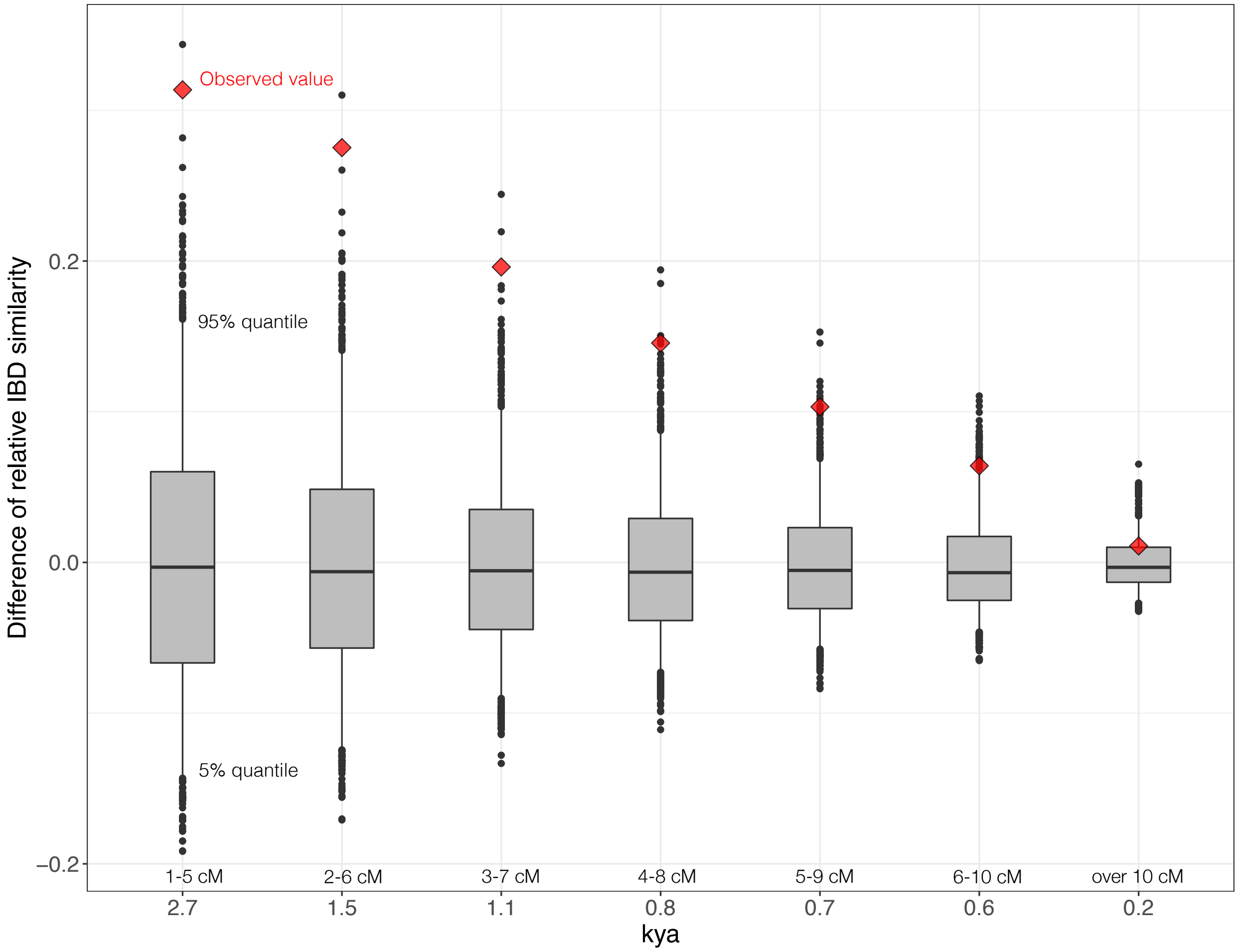


**Fig. S16.**

**Significance of the difference in relative IBD similarities between the Kula and non-Kula groups.** The red diamond is the observed value, while the boxplot is the distribution of simulated values obtained by randomly assigning groups as Kula or non-Kula. The upper whisker denotes the 95% quantile while the lower whisker denotes the 5% quantile.

**Table S1.**

**Metadata for the samples used in this study.** In the QC (quality control) column, samples were filtered by the indicated criteria in abbreviations, Kin: kinship up to 1st degree, Imiss: Individual with >5% missing data, PC: PCA outliers, Mixed: individual with parents speaking different languages or coming from different locations.

| Individual_ID | Population | Label_in_analysis | PNG_Province | District_Village_Island | Area | Region | Language_Family | Language | Longitude | Latitude | Reference | QC |
|---|---|---|---|---|---|---|---|---|---|---|---|---|
| MB_Ai01 | Arifama-Miniafia | Airara | NORTHERN | Airara | Massim | Oceania | Austronesian | Arifama-Miniafia | 149.3 | -9.51 | This_study | PASS |
| MB_Ai03 | Arifama-Miniafia | Airara | NORTHERN | Airara | Massim | Oceania | Austronesian | Arifama-Miniafia | 149.3 | -9.51 | This_study | PASS |
| MB_Ai05 | Arifama-Miniafia | Airara | NORTHERN | Airara | Massim | Oceania | Austronesian | Arifama-Miniafia | 149.3 | -9.51 | This_study | PASS |
| MB_Ai07 | Arifama-Miniafia | Airara | NORTHERN | Airara | Massim | Oceania | Austronesian | Arifama-Miniafia | 149.3 | -9.51 | This_study | PASS |
| MB_Ai09 | Arifama-Miniafia | Airara | NORTHERN | Airara | Massim | Oceania | Austronesian | Arifama-Miniafia | 149.3 | -9.51 | This_study | PASS |
| MB_Ai12 | Arifama-Miniafia | Airara | NORTHERN | Airara | Massim | Oceania | Austronesian | Arifama-Miniafia | 149.3 | -9.51 | This_study | PASS |
| MB_Ai14 | Arifama-Miniafia | Airara | NORTHERN | Airara | Massim | Oceania | Austronesian | Arifama-Miniafia | 149.3 | -9.51 | This_study | PASS |
| MB_Ai15 | Arifama-Miniafia | Airara | NORTHERN | Airara | Massim | Oceania | Austronesian | Arifama-Miniafia | 149.3 | -9.51 | This_study | PASS |
| MB_Ai17 | Arifama-Miniafia | Airara | NORTHERN | Airara | Massim | Oceania | Austronesian | Arifama-Miniafia | 149.3 | -9.51 | This_study | PASS |
| MB_Ai18 | Arifama-Miniafia | Airara | NORTHERN | Airara | Massim | Oceania | Austronesian | Arifama-Miniafia | 149.3 | -9.51 | This_study | PASS |
| MB_Ai24 | Arifama-Miniafia | Airara | NORTHERN | Airara | Massim | Oceania | Austronesian | Arifama-Miniafia | 149.3 | -9.51 | This_study | PASS |
| MB_Ai25 | Arifama-Miniafia | Airara | NORTHERN | Airara | Massim | Oceania | Austronesian | Arifama-Miniafia | 149.3 | -9.51 | This_study | PASS |
| MB_Ai29 | Arifama-Miniafia | Airara | NORTHERN | Airara | Massim | Oceania | Austronesian | Arifama-Miniafia | 149.3 | -9.51 | This_study | PASS |
| MB_Ai31 | Arifama-Miniafia | Airara | NORTHERN | Airara | Massim | Oceania | Austronesian | Arifama-Miniafia | 149.3 | -9.51 | This_study | PASS |
| MB_Ai32 | Arifama-Miniafia | Airara | NORTHERN | Airara | Massim | Oceania | Austronesian | Arifama-Miniafia | 149.3 | -9.51 | This_study | PASS |
| MB_Ai33 | Arifama-Miniafia | Airara | NORTHERN | Airara | Massim | Oceania | Austronesian | Arifama-Miniafia | 149.3 | -9.51 | This_study | PASS |
| MB_Ai36 | Arifama-Miniafia | Airara | NORTHERN | Airara | Massim | Oceania | Austronesian | Arifama-Miniafia | 149.3 | -9.51 | This_study | PASS |
| MB_Ai38 | Arifama-Miniafia | Airara | NORTHERN | Airara | Massim | Oceania | Austronesian | Arifama-Miniafia | 149.3 | -9.51 | This_study | PASS |
| MB_Ai39 | Arifama-Miniafia | Airara | NORTHERN | Airara | Massim | Oceania | Austronesian | Arifama-Miniafia | 149.3 | -9.51 | This_study | PASS |
| MB_Ha49 | Nimoa | Eastern_Calvados | MILNE_BAY | Eastern_Calvados_Dadahai-Kuanak-Nimoa-Joannet-Panawina-Sabarl-Wanim | Massim | Oceania | Austronesian | Nimoa | 153.06 | -11.16 | This_study | PASS |
| MB_Ni04 | Nimoa | Eastern_Calvados | MILNE_BAY | Eastern_Calvados_Dadahai-Kuanak-Nimoa-Joannet-Panawina-Sabarl-Wanim | Massim | Oceania | Austronesian | Nimoa | 153.25 | -11.3 | This_study | PASS |
| MB_Ni08 | Nimoa | Eastern_Calvados | MILNE_BAY | Eastern_Calvados_Dadahai-Kuanak-Nimoa-Joannet-Panawina-Sabarl-Wanim | Massim | Oceania | Austronesian | Nimoa | 153.25 | -11.3 | This_study | PASS |
| MB_Ni23 | Nimoa | Eastern_Calvados | MILNE_BAY | Eastern_Calvados_Dadahai-Kuanak-Nimoa-Joannet-Panawina-Sabarl-Wanim | Massim | Oceania | Austronesian | Nimoa | 153.25 | -11.3 | This_study | PASS |
| MB_Ni31 | Nimoa | Eastern_Calvados | MILNE_BAY | Eastern_Calvados_Dadahai-Kuanak-Nimoa-Joannet-Panawina-Sabarl-Wanim | Massim | Oceania | Austronesian | Nimoa | 153.25 | -11.3 | This_study | PASS |
| MB_Mo05 | Misima-Paneati_WC | Western_Calvados | MILNE_BAY | Western_Calvados_Motorina-Bagaman-Utian-Panaumala | Massim | Oceania | Austronesian | Misima-Paneati | 152.57 | -11.08 | This_study | PASS |

| | | | | | | | | | | | | |
|---|---|---|---|---|---|---|---|---|---|---|---|---|
| MB_Mo08 | Misima-Paneati_WC | Western_Calvados | MILNE_BAY | Western_Calvados_Motorina-Bagaman-Utian-Panaumala | Massim | Oceania | Austronesian | Misima-Paneati | 152.57 | -11.08 | This_study | PASS |
| MB_Mo10 | Misima-Paneati_WC | Western_Calvados | MILNE_BAY | Western_Calvados_Motorina-Bagaman-Utian-Panaumala | Massim | Oceania | Austronesian | Misima-Paneati | 152.57 | -11.08 | This_study | PASS |
| MB_Mo13 | Misima-Paneati_WC | Western_Calvados | MILNE_BAY | Western_Calvados_Motorina-Bagaman-Utian-Panaumala | Massim | Oceania | Austronesian | Misima-Paneati | 152.57 | -11.08 | This_study | PASS |
| MB_Mo14 | Misima-Paneati_WC | Western_Calvados | MILNE_BAY | Western_Calvados_Motorina-Bagaman-Utian-Panaumala | Massim | Oceania | Austronesian | Misima-Paneati | 152.57 | -11.08 | This_study | PASS |
| MB_Mo15 | Misima-Paneati_WC | Western_Calvados | MILNE_BAY | Western_Calvados_Motorina-Bagaman-Utian-Panaumala | Massim | Oceania | Austronesian | Misima-Paneati | 152.57 | -11.08 | This_study | PASS |
| MB_Mo19 | Misima-Paneati_WC | Western_Calvados | MILNE_BAY | Western_Calvados_Motorina-Bagaman-Utian-Panaumala | Massim | Oceania | Austronesian | Misima-Paneati | 152.57 | -11.08 | This_study | PASS |
| MB_Mo20 | Misima-Paneati_WC | Western_Calvados | MILNE_BAY | Western_Calvados_Motorina-Bagaman-Utian-Panaumala | Massim | Oceania | Austronesian | Misima-Paneati | 152.57 | -11.08 | This_study | PASS |
| MB_Mo23 | Misima-Paneati_WC | Western_Calvados | MILNE_BAY | Western_Calvados_Motorina-Bagaman-Utian-Panaumala | Massim | Oceania | Austronesian | Misima-Paneati | 152.57 | -11.08 | This_study | PASS |
| MB_Mo25 | Misima-Paneati_WC | Western_Calvados | MILNE_BAY | Western_Calvados_Motorina-Bagaman-Utian-Panaumala | Massim | Oceania | Austronesian | Misima-Paneati | 152.57 | -11.08 | This_study | PASS |
| MB_Mo26 | Misima-Paneati_WC | Western_Calvados | MILNE_BAY | Western_Calvados_Motorina-Bagaman-Utian-Panaumala | Massim | Oceania | Austronesian | Misima-Paneati | 152.57 | -11.08 | This_study | PASS |
| MB_Fe10 | Molima | Fergusson | MILNE_BAY | Fergusson-Dobu-Goodenough | Massim | Oceania | Austronesian | Molima | 150.54 | -9.46 | This_study | PASS |
| MB_Fe11 | Molima | Fergusson | MILNE_BAY | Fergusson-Dobu-Goodenough | Massim | Oceania | Austronesian | Molima | 150.54 | -9.46 | This_study | PASS |
| MB_Fe12 | Molima | Fergusson | MILNE_BAY | Fergusson-Dobu-Goodenough | Massim | Oceania | Austronesian | Molima | 150.54 | -9.46 | This_study | PASS |
| MB_Fe15 | Molima | Fergusson | MILNE_BAY | Fergusson-Dobu-Goodenough | Massim | Oceania | Austronesian | Molima | 150.54 | -9.46 | This_study | PASS |
| MB_Fe16 | Molima | Fergusson | MILNE_BAY | Fergusson-Dobu-Goodenough | Massim | Oceania | Austronesian | Molima | 150.54 | -9.46 | This_study | PASS |
| MB_Fe18 | Molima | Fergusson | MILNE_BAY | Fergusson-Dobu-Goodenough | Massim | Oceania | Austronesian | Molima | 150.54 | -9.46 | This_study | PASS |
| MB_Fe19 | Molima | Fergusson | MILNE_BAY | Fergusson-Dobu-Goodenough | Massim | Oceania | Austronesian | Molima | 150.54 | -9.46 | This_study | PASS |
| MB_Fe20 | Molima | Fergusson | MILNE_BAY | Fergusson-Dobu-Goodenough | Massim | Oceania | Austronesian | Molima | 150.54 | -9.46 | This_study | PASS |
| MB_Fe21 | Molima | Fergusson | MILNE_BAY | Fergusson-Dobu-Goodenough | Massim | Oceania | Austronesian | Molima | 150.54 | -9.46 | This_study | PASS |
| MB_Fe24 | Molima | Fergusson | MILNE_BAY | Fergusson-Dobu-Goodenough | Massim | Oceania | Austronesian | Molima | 150.54 | -9.46 | This_study | PASS |
| MB_Fe26 | Molima | Fergusson | MILNE_BAY | Fergusson-Dobu-Goodenough | Massim | Oceania | Austronesian | Molima | 150.54 | -9.46 | This_study | PASS |
| MB_Ha12 | Molima | Fergusson | MILNE_BAY | Fergusson-Dobu-Goodenough | Massim | Oceania | Austronesian | Molima | 150.54 | -9.46 | This_study | PASS |
| MB_Ha14 | Molima | Fergusson | MILNE_BAY | Fergusson-Dobu-Goodenough | Massim | Oceania | Austronesian | Molima | 150.54 | -9.46 | This_study | PASS |
| MB_Ha18 | Molima | Fergusson | MILNE_BAY | Fergusson-Dobu-Goodenough | Massim | Oceania | Austronesian | Molima | 150.54 | -9.46 | This_study | PASS |
| MB_Ha21 | Molima | Fergusson | MILNE_BAY | Fergusson-Dobu-Goodenough | Massim | Oceania | Austronesian | Molima | 150.54 | -9.46 | This_study | PASS |
| MB_Ha30 | Molima | Fergusson | MILNE_BAY | Fergusson-Dobu-Goodenough | Massim | Oceania | Austronesian | Molima | 150.54 | -9.46 | This_study | PASS |

| | | | | | | | | | | | | |
|---|---|---|---|---|---|---|---|---|---|---|---|---|
| MB_Ha45 | Molima | Fergusson | MILNE_BAY | Fergusson-Dobu-Goodenough | Massim | Oceania | Austronesian | Molima | 150.54 | -9.46 | This_study | PASS |
| MB_Ha53 | Molima | Fergusson | MILNE_BAY | Fergusson-Dobu-Goodenough | Massim | Oceania | Austronesian | Molima | 150.54 | -9.46 | This_study | PASS |
| MB_Ha54 | Molima | Fergusson | MILNE_BAY | Fergusson-Dobu-Goodenough | Massim | Oceania | Austronesian | Molima | 150.54 | -9.46 | This_study | PASS |
| MB_Ha62 | Molima | Fergusson | MILNE_BAY | Fergusson-Dobu-Goodenough | Massim | Oceania | Austronesian | Molima | 150.54 | -9.46 | This_study | PASS |
| MB_Ha67 | Molima | Fergusson | MILNE_BAY | Fergusson-Dobu-Goodenough | Massim | Oceania | Austronesian | Molima | 150.54 | -9.46 | This_study | Kin |
| MB_Ha68 | Molima | Fergusson | MILNE_BAY | Fergusson-Dobu-Goodenough | Massim | Oceania | Austronesian | Molima | 150.54 | -9.46 | This_study | PASS |
| MB_Ha80 | Molima | Fergusson | MILNE_BAY | Fergusson-Dobu-Goodenough | Massim | Oceania | Austronesian | Molima | 150.54 | -9.46 | This_study | PASS |
| MB_Ga03 | Muyuw_Gawa | Gawa | MILNE_BAY | Gawa | Massim | Oceania | Austronesian | Muyuw | 151.99 | -8.97 | This_study | PASS |
| MB_Ga04 | Muyuw_Gawa | Gawa | MILNE_BAY | Gawa | Massim | Oceania | Austronesian | Muyuw | 151.99 | -8.97 | This_study | PASS |
| MB_Ga06 | Muyuw_Gawa | Gawa | MILNE_BAY | Gawa | Massim | Oceania | Austronesian | Muyuw | 151.99 | -8.97 | This_study | PASS |
| MB_Ga09 | Muyuw_Gawa | Gawa | MILNE_BAY | Gawa | Massim | Oceania | Austronesian | Muyuw | 151.99 | -8.97 | This_study | PASS |
| MB_Ga10 | Muyuw_Gawa | Gawa | MILNE_BAY | Gawa | Massim | Oceania | Austronesian | Muyuw | 151.99 | -8.97 | This_study | PASS |
| MB_Ga12 | Muyuw_Gawa | Gawa | MILNE_BAY | Gawa | Massim | Oceania | Austronesian | Muyuw | 151.99 | -8.97 | This_study | PASS |
| MB_Ga14 | Muyuw_Gawa | Gawa | MILNE_BAY | Gawa | Massim | Oceania | Austronesian | Muyuw | 151.99 | -8.97 | This_study | PASS |
| MB_Ga15 | Muyuw_Gawa | Gawa | MILNE_BAY | Gawa | Massim | Oceania | Austronesian | Muyuw | 151.99 | -8.97 | This_study | PASS |
| MB_Ga16 | Muyuw_Gawa | Gawa | MILNE_BAY | Gawa | Massim | Oceania | Austronesian | Muyuw | 151.99 | -8.97 | This_study | PASS |
| MB_Ga17 | Muyuw_Gawa | Gawa | MILNE_BAY | Gawa | Massim | Oceania | Austronesian | Muyuw | 151.99 | -8.97 | This_study | PASS |
| MB_Ga18 | Muyuw_Gawa | Gawa | MILNE_BAY | Gawa | Massim | Oceania | Austronesian | Muyuw | 151.99 | -8.97 | This_study | PASS |
| MB_Ha07 | Tawala | Mainland_Eastern_Tip | MILNE_BAY | Mline_Bay_mainland_eastern_tip | Massim | Oceania | Austronesian | Tawala | 150.53 | -10.49 | This_study | PASS |
| MB_Ha10 | Tawala | Mainland_Eastern_Tip | MILNE_BAY | Mline_Bay_mainland_eastern_tip | Massim | Oceania | Austronesian | Tawala | 150.53 | -10.49 | This_study | PASS |
| MB_Ha13 | Tawala | Mainland_Eastern_Tip | MILNE_BAY | Mline_Bay_mainland_eastern_tip | Massim | Oceania | Austronesian | Tawala | 150.53 | -10.49 | This_study | PASS |
| MB_Ha17 | Tawala | Mainland_Eastern_Tip | MILNE_BAY | Mline_Bay_mainland_eastern_tip | Massim | Oceania | Austronesian | Tawala | 150.53 | -10.49 | This_study | PASS |
| MB_Ha32 | Tawala | Mainland_Eastern_Tip | MILNE_BAY | Mline_Bay_mainland_eastern_tip | Massim | Oceania | Austronesian | Tawala | 150.53 | -10.49 | This_study | PASS |
| MB_Ha33 | Tawala | Mainland_Eastern_Tip | MILNE_BAY | Mline_Bay_mainland_eastern_tip | Massim | Oceania | Austronesian | Tawala | 150.53 | -10.49 | This_study | PASS |
| MB_Ha42 | Tawala | Mainland_Eastern_Tip | MILNE_BAY | Mline_Bay_mainland_eastern_tip | Massim | Oceania | Austronesian | Tawala | 150.53 | -10.49 | This_study | PASS |
| MB_Ha50 | Tawala | Mainland_Eastern_Tip | MILNE_BAY | Mline_Bay_mainland_eastern_tip | Massim | Oceania | Austronesian | Tawala | 150.53 | -10.49 | This_study | Imiss |
| MB_Ha51 | Tawala | Mainland_Eastern_Tip | MILNE_BAY | Mline_Bay_mainland_eastern_tip | Massim | Oceania | Austronesian | Tawala | 150.53 | -10.49 | This_study | PASS |
| MB_Ha64 | Tawala | Mainland_Eastern_Tip | MILNE_BAY | Mline_Bay_mainland_eastern_tip | Massim | Oceania | Austronesian | Tawala | 150.53 | -10.49 | This_study | PASS |
| MB_Ha70 | Tawala | Mainland_Eastern_Tip | MILNE_BAY | Mline_Bay_mainland_eastern_tip | Massim | Oceania | Austronesian | Tawala | 150.53 | -10.49 | This_study | PASS |
| MB_Ha74 | Tawala | Mainland_Eastern_Tip | MILNE_BAY | Mline_Bay_mainland_eastern_tip | Massim | Oceania | Austronesian | Tawala | 150.53 | -10.49 | This_study | PASS |
| MB_Ji014 | Tawala | Mainland_Eastern_Tip | MILNE_BAY | Mline_Bay_mainland_eastern_tip | Massim | Oceania | Austronesian | Tawala | 150.53 | -10.49 | This_study | PASS |
| MB_Mo02 | Misima-Paneati | Misima | MILNE_BAY | Misima-Paneati-Panapompom-Kimuta | Massim | Oceania | Austronesian | Misima-Paneati | 152.72 | -10.67 | This_study | PASS |
| MB_Mo16 | Misima-Paneati | Misima | MILNE_BAY | Misima-Paneati-Panapompom-Kimuta | Massim | Oceania | Austronesian | Misima-Paneati | 152.72 | -10.67 | This_study | PASS |
| MB_Mo18 | Misima-Paneati | Misima | MILNE_BAY | Misima-Paneati-Panapompom-Kimuta | Massim | Oceania | Austronesian | Misima-Paneati | 152.72 | -10.67 | This_study | PASS |
| MB_Ni38 | Misima-Paneati | Misima | MILNE_BAY | Misima-Paneati-Panapompom-Kimuta | Massim | Oceania | Austronesian | Misima-Paneati | 152.72 | -10.67 | This_study | PASS |

| | | | | | | | | | | | | |
|---|---|---|---|---|---|---|---|---|---|---|---|---|
| MB_Fe02 | Bunana | Normanby | MILNE_BAY | Normanby | Massim | Oceania | Austronesian | Bunana | 151.09 | -10.08 | This_study | PASS |
| MB_Fe03 | Bunana | Normanby | MILNE_BAY | Normanby | Massim | Oceania | Austronesian | Bunana | 151.09 | -10.08 | This_study | PASS |
| MB_Fe04 | Bunana | Normanby | MILNE_BAY | Normanby | Massim | Oceania | Austronesian | Bunana | 151.09 | -10.08 | This_study | PASS |
| MB_Fe23 | Bunana | Normanby | MILNE_BAY | Normanby | Massim | Oceania | Austronesian | Bunana | 151.09 | -10.08 | This_study | PASS |
| MB_Ha05 | Bunana | Normanby | MILNE_BAY | Normanby | Massim | Oceania | Austronesian | Bunana | 151.09 | -10.08 | This_study | PASS |
| MB_Ha08 | Bunana | Normanby | MILNE_BAY | Normanby | Massim | Oceania | Austronesian | Bunana | 151.09 | -10.08 | This_study | PASS |
| MB_Ha09 | Bunana | Normanby | MILNE_BAY | Normanby | Massim | Oceania | Austronesian | Bunana | 151.09 | -10.08 | This_study | PASS |
| MB_Ha11 | Bunana | Normanby | MILNE_BAY | Normanby | Massim | Oceania | Austronesian | Bunana | 151.09 | -10.08 | This_study | PASS |
| MB_Ha20 | Bunana | Normanby | MILNE_BAY | Normanby | Massim | Oceania | Austronesian | Bunana | 151.09 | -10.08 | This_study | PASS |
| MB_Ha26 | Bunana | Normanby | MILNE_BAY | Normanby | Massim | Oceania | Austronesian | Bunana | 151.09 | -10.08 | This_study | PASS |
| MB_Ha28 | Bunana | Normanby | MILNE_BAY | Normanby | Massim | Oceania | Austronesian | Bunana | 151.09 | -10.08 | This_study | PASS |
| MB_Ha35 | Bunana | Normanby | MILNE_BAY | Normanby | Massim | Oceania | Austronesian | Bunana | 151.09 | -10.08 | This_study | PASS |
| MB_Ha56 | Bunana | Normanby | MILNE_BAY | Normanby | Massim | Oceania | Austronesian | Bunana | 151.09 | -10.08 | This_study | PASS |
| MB_Ha58 | Bunana | Normanby | MILNE_BAY | Normanby | Massim | Oceania | Austronesian | Bunana | 151.09 | -10.08 | This_study | PASS |
| MB_Ha76 | Bunana | Normanby | MILNE_BAY | Normanby | Massim | Oceania | Austronesian | Bunana | 151.09 | -10.08 | This_study | PASS |
| MB_Ha84 | Bunana | Normanby | MILNE_BAY | Normanby | Massim | Oceania | Austronesian | Bunana | 151.09 | -10.08 | This_study | PASS |
| MB_Ha85 | Bunana | Normanby | MILNE_BAY | Normanby | Massim | Oceania | Austronesian | Bunana | 151.09 | -10.08 | This_study | PASS |
| MB_Su02 | Sudest | Sudest | MILNE_BAY | Sudest | Massim | Oceania | Austronesian | Sudest | 153.46 | -11.52 | This_study | PASS |
| MB_Su03 | Sudest | Sudest | MILNE_BAY | Sudest | Massim | Oceania | Austronesian | Sudest | 153.46 | -11.52 | This_study | PASS |
| MB_Su04 | Sudest | Sudest | MILNE_BAY | Sudest | Massim | Oceania | Austronesian | Sudest | 153.46 | -11.52 | This_study | PASS |
| MB_Su06 | Sudest | Sudest | MILNE_BAY | Sudest | Massim | Oceania | Austronesian | Sudest | 153.46 | -11.52 | This_study | PASS |
| MB_Su07 | Sudest | Sudest | MILNE_BAY | Sudest | Massim | Oceania | Austronesian | Sudest | 153.46 | -11.52 | This_study | PASS |
| MB_Su08 | Sudest | Sudest | MILNE_BAY | Sudest | Massim | Oceania | Austronesian | Sudest | 153.46 | -11.52 | This_study | PASS |
| MB_Su10 | Sudest | Sudest | MILNE_BAY | Sudest | Massim | Oceania | Austronesian | Sudest | 153.46 | -11.52 | This_study | PASS |
| MB_Su12 | Sudest | Sudest | MILNE_BAY | Sudest | Massim | Oceania | Austronesian | Sudest | 153.46 | -11.52 | This_study | PASS |
| MB_Su14 | Sudest | Sudest | MILNE_BAY | Sudest | Massim | Oceania | Austronesian | Sudest | 153.46 | -11.52 | This_study | PASS |
| MB_Su20 | Sudest | Sudest | MILNE_BAY | Sudest | Massim | Oceania | Austronesian | Sudest | 153.46 | -11.52 | This_study | PASS |
| MB_Su22 | Sudest | Sudest | MILNE_BAY | Sudest | Massim | Oceania | Austronesian | Sudest | 153.46 | -11.52 | This_study | PASS |
| MB_Su25 | Sudest | Sudest | MILNE_BAY | Sudest | Massim | Oceania | Austronesian | Sudest | 153.46 | -11.52 | This_study | PASS |
| MB_Su26 | Sudest | Sudest | MILNE_BAY | Sudest | Massim | Oceania | Austronesian | Sudest | 153.46 | -11.52 | This_study | PASS |
| MB_Su28 | Sudest | Sudest | MILNE_BAY | Sudest | Massim | Oceania | Austronesian | Sudest | 153.46 | -11.52 | This_study | PASS |
| MB_Su29 | Sudest | Sudest | MILNE_BAY | Sudest | Massim | Oceania | Austronesian | Sudest | 153.46 | -11.52 | This_study | PASS |
| MB_Su30 | Sudest | Sudest | MILNE_BAY | Sudest | Massim | Oceania | Austronesian | Sudest | 153.46 | -11.52 | This_study | PASS |
| MB_Wa06 | Maisin | Wanigela | NORTHERN | Wanigela | Massim | Oceania | Austronesian | Maisin | 149.18 | -9.34 | This_study | PASS |
| MB_Wa07 | Maisin | Wanigela | NORTHERN | Wanigela | Massim | Oceania | Austronesian | Maisin | 149.18 | -9.34 | This_study | PASS |
| MB_Wa09 | Maisin | Wanigela | NORTHERN | Wanigela | Massim | Oceania | Austronesian | Maisin | 149.18 | -9.34 | This_study | PASS |
| MB_Wa11 | Maisin | Wanigela | NORTHERN | Wanigela | Massim | Oceania | Austronesian | Maisin | 149.18 | -9.34 | This_study | PASS |
| MB_Wa13 | Maisin | Wanigela | NORTHERN | Wanigela | Massim | Oceania | Austronesian | Maisin | 149.18 | -9.34 | This_study | PASS |
| MB_Wa14 | Maisin | Wanigela | NORTHERN | Wanigela | Massim | Oceania | Austronesian | Maisin | 149.18 | -9.34 | This_study | PASS |
| MB_Wa15 | Maisin | Wanigela | NORTHERN | Wanigela | Massim | Oceania | Austronesian | Maisin | 149.18 | -9.34 | This_study | PASS |
| MB_Wa16 | Maisin | Wanigela | NORTHERN | Wanigela | Massim | Oceania | Austronesian | Maisin | 149.18 | -9.34 | This_study | PASS |
| MB_Wa18 | Maisin | Wanigela | NORTHERN | Wanigela | Massim | Oceania | Austronesian | Maisin | 149.18 | -9.34 | This_study | PASS |
| MB_Wa19 | Maisin | Wanigela | NORTHERN | Wanigela | Massim | Oceania | Austronesian | Maisin | 149.18 | -9.34 | This_study | PASS |
| MB_Wa20 | Maisin | Wanigela | NORTHERN | Wanigela | Massim | Oceania | Austronesian | Maisin | 149.18 | -9.34 | This_study | PASS |
| MB_Wa21 | Maisin | Wanigela | NORTHERN | Wanigela | Massim | Oceania | Austronesian | Maisin | 149.18 | -9.34 | This_study | PASS |
| MB_Wa22 | Maisin | Wanigela | NORTHERN | Wanigela | Massim | Oceania | Austronesian | Maisin | 149.18 | -9.34 | This_study | PASS |
| MB_Wa23 | Maisin | Wanigela | NORTHERN | Wanigela | Massim | Oceania | Austronesian | Maisin | 149.18 | -9.34 | This_study | PASS |
| MB_Wa26 | Maisin | Wanigela | NORTHERN | Wanigela | Massim | Oceania | Austronesian | Maisin | 149.18 | -9.34 | This_study | PASS |
| MB_Bu01 | Budibud | Laughlan | MILNE_BAY | Laughlan | Massim | Oceania | Austronesian | Budibud | 153.69 | -9.29 | This_study | PASS |

| | | | | | | | | | | | | |
|---|---|---|---|---|---|---|---|---|---|---|---|---|
| MB_Bu02 | Budibud | Laughlan | MILNE_BAY | Laughlan | Massim | Oceania | Austronesian | Budibud | 153.69 | -9.29 | This_study | PASS |
| MB_Bu05 | Budibud | Laughlan | MILNE_BAY | Laughlan | Massim | Oceania | Austronesian | Budibud | 153.69 | -9.29 | This_study | PASS |
| MB_Bu06 | Budibud | Laughlan | MILNE_BAY | Laughlan | Massim | Oceania | Austronesian | Budibud | 153.69 | -9.29 | This_study | PASS |
| MB_Bu07 | Budibud | Laughlan | MILNE_BAY | Laughlan | Massim | Oceania | Austronesian | Budibud | 153.69 | -9.29 | This_study | PASS |
| MB_Bu09 | Budibud | Laughlan | MILNE_BAY | Laughlan | Massim | Oceania | Austronesian | Budibud | 153.69 | -9.29 | This_study | PASS |
| MB_Ha59 | Budibud | Laughlan | MILNE_BAY | Laughlan | Massim | Oceania | Austronesian | Budibud | 153.69 | -9.29 | This_study | PASS |
| MB_Wo04 | Muyuw_Woodlark | Woodlark | MILNE_BAY | Woodlark | Massim | Oceania | Austronesian | Muyuw | 152.78 | -9.12 | This_study | PASS |
| MB_Wo06 | Muyuw_Woodlark | Woodlark | MILNE_BAY | Woodlark | Massim | Oceania | Austronesian | Muyuw | 152.78 | -9.12 | This_study | PASS |
| MB_Wo09 | Muyuw_Woodlark | Woodlark | MILNE_BAY | Woodlark | Massim | Oceania | Austronesian | Muyuw | 152.78 | -9.12 | This_study | PASS |
| MB_Wo10 | Muyuw_Woodlark | Woodlark | MILNE_BAY | Woodlark | Massim | Oceania | Austronesian | Muyuw | 152.78 | -9.12 | This_study | PASS |
| MB_Wo14 | Muyuw_Woodlark | Woodlark | MILNE_BAY | Woodlark | Massim | Oceania | Austronesian | Muyuw | 152.78 | -9.12 | This_study | PASS |
| MB_Wo18 | Muyuw_Woodlark | Woodlark | MILNE_BAY | Woodlark | Massim | Oceania | Austronesian | Muyuw | 152.78 | -9.12 | This_study | PASS |
| MB_Wo20 | Muyuw_Woodlark | Woodlark | MILNE_BAY | Woodlark | Massim | Oceania | Austronesian | Muyuw | 152.78 | -9.12 | This_study | PASS |
| MB_Ga01 | Kilivila | Trobriand | MILNE_BAY | Trobriand | Massim | Oceania | Austronesian | Kilivila | 151.11 | -8.67 | This_study | PASS |
| MB_Wo19 | Kilivila | Trobriand | MILNE_BAY | Trobriand | Massim | Oceania | Austronesian | Kilivila | 151.11 | -8.67 | This_study | PASS |
| MB_Wo21 | Kilivila | Trobriand | MILNE_BAY | Trobriand | Massim | Oceania | Austronesian | Kilivila | 151.11 | -8.67 | This_study | PASS |
| T02 | Kilivila | Trobriand | MILNE_BAY | Trobriand | Massim | Oceania | Austronesian | Kilivila | 151.11 | -8.67 | This_study | PASS |
| T09 | Kilivila | Trobriand | MILNE_BAY | Trobriand | Massim | Oceania | Austronesian | Kilivila | 151.11 | -8.67 | This_study | PASS |
| T16 | Kilivila | Trobriand | MILNE_BAY | Trobriand | Massim | Oceania | Austronesian | Kilivila | 151.11 | -8.67 | This_study | PASS |
| T36 | Kilivila | Trobriand | MILNE_BAY | Trobriand | Massim | Oceania | Austronesian | Kilivila | 151.11 | -8.67 | This_study | PASS |
| T40 | Kilivila | Trobriand | MILNE_BAY | Trobriand | Massim | Oceania | Austronesian | Kilivila | 151.11 | -8.67 | This_study | PASS |
| T47 | Kilivila | Trobriand | MILNE_BAY | Trobriand | Massim | Oceania | Austronesian | Kilivila | 151.11 | -8.67 | This_study | PASS |
| T63 | Kilivila | Trobriand | MILNE_BAY | Trobriand | Massim | Oceania | Austronesian | Kilivila | 151.11 | -8.67 | This_study | PASS |
| T66 | Kilivila | Trobriand | MILNE_BAY | Trobriand | Massim | Oceania | Austronesian | Kilivila | 151.11 | -8.67 | This_study | PASS |
| T68 | Kilivila | Trobriand | MILNE_BAY | Trobriand | Massim | Oceania | Austronesian | Kilivila | 151.11 | -8.67 | This_study | PASS |
| Ji_058 | Yele | Rossel | MILNE_BAY | Rossel | Massim | Oceania | Yele-West_New_Britain | Yele | 154.18 | -11.36 | This_study | PASS |
| Ji_059 | Yele | Rossel | MILNE_BAY | Rossel | Massim | Oceania | Yele-West_New_Britain | Yele | 154.18 | -11.36 | This_study | PASS |
| Ji_060 | Yele | Rossel | MILNE_BAY | Rossel | Massim | Oceania | Yele-West_New_Britain | Yele | 154.18 | -11.36 | This_study | PASS |
| Ji_061 | Yele | Rossel | MILNE_BAY | Rossel | Massim | Oceania | Yele-West_New_Britain | Yele | 154.18 | -11.36 | This_study | PASS |
| Ji_062 | Yele | Rossel | MILNE_BAY | Rossel | Massim | Oceania | Yele-West_New_Britain | Yele | 154.18 | -11.36 | This_study | PASS |
| Ji_063 | Yele | Rossel | MILNE_BAY | Rossel | Massim | Oceania | Yele-West_New_Britain | Yele | 154.18 | -11.36 | This_study | PASS |
| Ji_064 | Yele | Rossel | MILNE_BAY | Rossel | Massim | Oceania | Yele-West_New_Britain | Yele | 154.18 | -11.36 | This_study | PASS |
| Ji_065 | Yele | Rossel | MILNE_BAY | Rossel | Massim | Oceania | Yele-West_New_Britain | Yele | 154.18 | -11.36 | This_study | PASS |
| Ji_066 | Yele | Rossel | MILNE_BAY | Rossel | Massim | Oceania | Yele-West_New_Britain | Yele | 154.18 | -11.36 | This_study | PASS |
| Ji_067 | Yele | Rossel | MILNE_BAY | Rossel | Massim | Oceania | Yele-West_New_Britain | Yele | 154.18 | -11.36 | This_study | PASS |
| Ji_069 | Yele | Rossel | MILNE_BAY | Rossel | Massim | Oceania | Yele-West_New_Britain | Yele | 154.18 | -11.36 | This_study | PASS |
| Ji_070 | Yele | Rossel | MILNE_BAY | Rossel | Massim | Oceania | Yele-West_New_Britain | Yele | 154.18 | -11.36 | This_study | PASS |
| Ji_072 | Yele | Rossel | MILNE_BAY | Rossel | Massim | Oceania | Yele-West_New_Britain | Yele | 154.18 | -11.36 | This_study | PASS |
| Ji_073 | Yele | Rossel | MILNE_BAY | Rossel | Massim | Oceania | Yele-West_New_Britain | Yele | 154.18 | -11.36 | This_study | PASS |
| Ji_074 | Yele | Rossel | MILNE_BAY | Rossel | Massim | Oceania | Yele-West_New_Britain | Yele | 154.18 | -11.36 | This_study | PASS |
| Ji_076 | Yele | Rossel | MILNE_BAY | Rossel | Massim | Oceania | Yele-West_New_Britain | Yele | 154.18 | -11.36 | This_study | PASS |

| | | | | | | | | | | | | |
|---|---|---|---|---|---|---|---|---|---|---|---|---|
| Ji_077 | Yele | Rossel | MILNE_BAY | Rossel | Massim | Oceania | Yele-West_New_Britain | Yele | 154.18 | -11.36 | This_study | PASS |
| Ji_078 | Yele | Rossel | MILNE_BAY | Rossel | Massim | Oceania | Yele-West_New_Britain | Yele | 154.18 | -11.36 | This_study | PASS |
| Ji_079 | Yele | Rossel | MILNE_BAY | Rossel | Massim | Oceania | Yele-West_New_Britain | Yele | 154.18 | -11.36 | This_study | PASS |
| Ji_081 | Yele | Rossel | MILNE_BAY | Rossel | Massim | Oceania | Yele-West_New_Britain | Yele | 154.18 | -11.36 | This_study | PASS |
| Ji_083 | Yele | Rossel | MILNE_BAY | Rossel | Massim | Oceania | Yele-West_New_Britain | Yele | 154.18 | -11.36 | This_study | PASS |
| Ji_085 | Yele | Rossel | MILNE_BAY | Rossel | Massim | Oceania | Yele-West_New_Britain | Yele | 154.18 | -11.36 | This_study | PASS |
| Ji_086 | Yele | Rossel | MILNE_BAY | Rossel | Massim | Oceania | Yele-West_New_Britain | Yele | 154.18 | -11.36 | This_study | Kin |
| Ji_087 | Yele | Rossel | MILNE_BAY | Rossel | Massim | Oceania | Yele-West_New_Britain | Yele | 154.18 | -11.36 | This_study | PASS |
| Ji_089 | Yele | Rossel | MILNE_BAY | Rossel | Massim | Oceania | Yele-West_New_Britain | Yele | 154.18 | -11.36 | This_study | PASS |
| Ji_090 | Yele | Rossel | MILNE_BAY | Rossel | Massim | Oceania | Yele-West_New_Britain | Yele | 154.18 | -11.36 | This_study | PASS |
| Ji_092 | Yele | Rossel | MILNE_BAY | Rossel | Massim | Oceania | Yele-West_New_Britain | Yele | 154.18 | -11.36 | This_study | PASS |
| Ji_093 | Yele | Rossel | MILNE_BAY | Rossel | Massim | Oceania | Yele-West_New_Britain | Yele | 154.18 | -11.36 | This_study | PASS |
| Ji_094 | Yele | Rossel | MILNE_BAY | Rossel | Massim | Oceania | Yele-West_New_Britain | Yele | 154.18 | -11.36 | This_study | PASS |
| Ji_095 | Yele | Rossel | MILNE_BAY | Rossel | Massim | Oceania | Yele-West_New_Britain | Yele | 154.18 | -11.36 | This_study | PASS |
| Ji_096 | Yele | Rossel | MILNE_BAY | Rossel | Massim | Oceania | Yele-West_New_Britain | Yele | 154.18 | -11.36 | This_study | PASS |
| Ji_097 | Yele | Rossel | MILNE_BAY | Rossel | Massim | Oceania | Yele-West_New_Britain | Yele | 154.18 | -11.36 | This_study | PASS |
| Ka01 | Purari_Kapuna | Gulf | GULF | Kapuna | PNG_Highland | Oceania | Trans-New_Guinea | Purari | 144.97 | -7.58 | This_study | PASS |
| Ka02 | Purari_Kapuna | Gulf | GULF | Kapuna | PNG_Highland | Oceania | Trans-New_Guinea | Purari | 144.97 | -7.58 | This_study | PASS |
| Ka05 | Purari_Kapuna | Gulf | GULF | Kapuna | PNG_Highland | Oceania | Trans-New_Guinea | Purari | 144.97 | -7.58 | This_study | PASS |
| Ka10 | Purari_Kapuna | Gulf | GULF | Kapuna | PNG_Highland | Oceania | Trans-New_Guinea | Purari | 144.97 | -7.58 | This_study | PASS |
| Ka11 | Purari_Kapuna | Gulf | GULF | Kapuna | PNG_Highland | Oceania | Trans-New_Guinea | Purari | 144.97 | -7.58 | This_study | PASS |
| Ka12 | Purari_Kapuna | Gulf | GULF | Kapuna | PNG_Highland | Oceania | Trans-New_Guinea | Purari | 144.97 | -7.58 | This_study | PASS |
| Ka13 | Purari_Kapuna | Gulf | GULF | Kapuna | PNG_Highland | Oceania | Trans-New_Guinea | Purari | 144.97 | -7.58 | This_study | PASS |
| Ka14 | Purari_Kapuna | Gulf | GULF | Kapuna | PNG_Highland | Oceania | Trans-New_Guinea | Purari | 144.97 | -7.58 | This_study | PASS |
| Ka15 | Purari_Kapuna | Gulf | GULF | Kapuna | PNG_Highland | Oceania | Trans-New_Guinea | Purari | 144.97 | -7.58 | This_study | PASS |
| Ka16 | Purari_Kapuna | Gulf | GULF | Kapuna | PNG_Highland | Oceania | Trans-New_Guinea | Purari | 144.97 | -7.58 | This_study | PASS |
| Ka17 | Purari_Kapuna | Gulf | GULF | Kapuna | PNG_Highland | Oceania | Trans-New_Guinea | Purari | 144.97 | -7.58 | This_study | PASS |
| Ka19 | Purari_Kapuna | Gulf | GULF | Kapuna | PNG_Highland | Oceania | Trans-New_Guinea | Purari | 144.97 | -7.58 | This_study | PASS |
| Ka20 | Purari_Kapuna | Gulf | GULF | Kapuna | PNG_Highland | Oceania | Trans-New_Guinea | Purari | 144.97 | -7.58 | This_study | PASS |
| Ka21 | Purari_Kapuna | Gulf | GULF | Kapuna | PNG_Highland | Oceania | Trans-New_Guinea | Purari | 144.97 | -7.58 | This_study | PASS |
| Ka22 | Purari_Kapuna | Gulf | GULF | Kapuna | PNG_Highland | Oceania | Trans-New_Guinea | Purari | 144.97 | -7.58 | This_study | PASS |
| Ka23 | Purari_Kapuna | Gulf | GULF | Kapuna | PNG_Highland | Oceania | Trans-New_Guinea | Purari | 144.97 | -7.58 | This_study | PASS |
| Ka24 | Purari_Kapuna | Gulf | GULF | Kapuna | PNG_Highland | Oceania | Trans-New_Guinea | Purari | 144.97 | -7.58 | This_study | PASS |

| | | | | | | | | | | | | |
|---|---|---|---|---|---|---|---|---|---|---|---|---|
| Ka25 | Purari_Kapuna | Gulf | GULF | Kapuna | PNG_Highland | Oceania | Trans-New_Guinea | Purari | 144.97 | -7.58 | This_study | PASS |
| Ka27 | Purari_Kapuna | Gulf | GULF | Kapuna | PNG_Highland | Oceania | Trans-New_Guinea | Purari | 144.97 | -7.58 | This_study | PASS |
| Ka28 | Purari_Kapuna | Gulf | GULF | Kapuna | PNG_Highland | Oceania | Trans-New_Guinea | Purari | 144.97 | -7.58 | This_study | PASS |
| Ka31 | Purari_Kapuna | Gulf | GULF | Kapuna | PNG_Highland | Oceania | Trans-New_Guinea | Purari | 144.97 | -7.58 | This_study | PASS |
| Ka32 | Purari_Kapuna | Gulf | GULF | Kapuna | PNG_Highland | Oceania | Trans-New_Guinea | Purari | 144.97 | -7.58 | This_study | PASS |
| Ka33 | Purari_Kapuna | Gulf | GULF | Kapuna | PNG_Highland | Oceania | Trans-New_Guinea | Purari | 144.97 | -7.58 | This_study | Kin |
| Ka34 | Purari_Kapuna | Gulf | GULF | Kapuna | PNG_Highland | Oceania | Trans-New_Guinea | Purari | 144.97 | -7.58 | This_study | PASS |
| Ka35 | Purari_Kapuna | Gulf | GULF | Kapuna | PNG_Highland | Oceania | Trans-New_Guinea | Purari | 144.97 | -7.58 | This_study | PASS |
| Ka37 | Purari_Kapuna | Gulf | GULF | Kapuna | PNG_Highland | Oceania | Trans-New_Guinea | Purari | 144.97 | -7.58 | This_study | PASS |
| Ka38 | Purari_Kapuna | Gulf | GULF | Kapuna | PNG_Highland | Oceania | Trans-New_Guinea | Purari | 144.97 | -7.58 | This_study | PASS |
| Ka39 | Purari_Kapuna | Gulf | GULF | Kapuna | PNG_Highland | Oceania | Trans-New_Guinea | Purari | 144.97 | -7.58 | This_study | PASS |
| Ka40 | Purari_Kapuna | Gulf | GULF | Kapuna | PNG_Highland | Oceania | Trans-New_Guinea | Purari | 144.97 | -7.58 | This_study | PASS |
| Ka42 | Purari_Kapuna | Gulf | GULF | Kapuna | PNG_Highland | Oceania | Trans-New_Guinea | Purari | 144.97 | -7.58 | This_study | PASS |
| Ka43 | Purari_Kapuna | Gulf | GULF | Kapuna | PNG_Highland | Oceania | Trans-New_Guinea | Purari | 144.97 | -7.58 | This_study | PASS |
| Ka44 | Purari_Kapuna | Gulf | GULF | Kapuna | PNG_Highland | Oceania | Trans-New_Guinea | Purari | 144.97 | -7.58 | This_study | PASS |
| Ka45 | Purari_Kapuna | Gulf | GULF | Kapuna | PNG_Highland | Oceania | Trans-New_Guinea | Purari | 144.97 | -7.58 | This_study | PASS |
| Be01 | Waima_Bereina | Central | CENTRAL | Bereina | PNG_Lowland | Oceania | Austronesian | Waima | 146.51 | -8.63 | This_study | PASS |
| Be03 | Waima_Bereina | Central | CENTRAL | Bereina | PNG_Lowland | Oceania | Austronesian | Waima | 146.51 | -8.63 | This_study | PASS |
| Be06 | Waima_Bereina | Central | CENTRAL | Bereina | PNG_Lowland | Oceania | Austronesian | Waima | 146.51 | -8.63 | This_study | PASS |
| Be07 | Waima_Bereina | Central | CENTRAL | Bereina | PNG_Lowland | Oceania | Austronesian | Waima | 146.51 | -8.63 | This_study | PASS |
| Be09 | Waima_Bereina | Central | CENTRAL | Bereina | PNG_Lowland | Oceania | Austronesian | Waima | 146.51 | -8.63 | This_study | PASS |
| Be10 | Waima_Bereina | Central | CENTRAL | Bereina | PNG_Lowland | Oceania | Austronesian | Waima | 146.51 | -8.63 | This_study | PASS |
| Be11 | Waima_Bereina | Central | CENTRAL | Bereina | PNG_Lowland | Oceania | Austronesian | Waima | 146.51 | -8.63 | This_study | PASS |
| Be12 | Waima_Bereina | Central | CENTRAL | Bereina | PNG_Lowland | Oceania | Austronesian | Waima | 146.51 | -8.63 | This_study | PASS |
| Be13 | Waima_Bereina | Central | CENTRAL | Bereina | PNG_Lowland | Oceania | Austronesian | Waima | 146.51 | -8.63 | This_study | PASS |
| Be14 | Waima_Bereina | Central | CENTRAL | Bereina | PNG_Lowland | Oceania | Austronesian | Waima | 146.51 | -8.63 | This_study | PASS |
| Be15 | Waima_Bereina | Central | CENTRAL | Bereina | PNG_Lowland | Oceania | Austronesian | Waima | 146.51 | -8.63 | This_study | PASS |
| Be16 | Waima_Bereina | Central | CENTRAL | Bereina | PNG_Lowland | Oceania | Austronesian | Waima | 146.51 | -8.63 | This_study | PASS |
| Be17 | Waima_Bereina | Central | CENTRAL | Bereina | PNG_Lowland | Oceania | Austronesian | Waima | 146.51 | -8.63 | This_study | PASS |
| Be18 | Waima_Bereina | Central | CENTRAL | Bereina | PNG_Lowland | Oceania | Austronesian | Waima | 146.51 | -8.63 | This_study | PASS |
| Be19 | Waima_Bereina | Central | CENTRAL | Bereina | PNG_Lowland | Oceania | Austronesian | Waima | 146.51 | -8.63 | This_study | PASS |
| Be20 | Waima_Bereina | Central | CENTRAL | Bereina | PNG_Lowland | Oceania | Austronesian | Waima | 146.51 | -8.63 | This_study | PASS |
| Be21 | Waima_Bereina | Central | CENTRAL | Bereina | PNG_Lowland | Oceania | Austronesian | Waima | 146.51 | -8.63 | This_study | PASS |
| Be22 | Waima_Bereina | Central | CENTRAL | Bereina | PNG_Lowland | Oceania | Austronesian | Waima | 146.51 | -8.63 | This_study | PASS |
| Be23 | Waima_Bereina | Central | CENTRAL | Bereina | PNG_Lowland | Oceania | Austronesian | Waima | 146.51 | -8.63 | This_study | PASS |
| Be24 | Waima_Bereina | Central | CENTRAL | Bereina | PNG_Lowland | Oceania | Austronesian | Waima | 146.51 | -8.63 | This_study | PASS |
| Be25 | Waima_Bereina | Central | CENTRAL | Bereina | PNG_Lowland | Oceania | Austronesian | Waima | 146.51 | -8.63 | This_study | PASS |
| Be26 | Waima_Bereina | Central | CENTRAL | Bereina | PNG_Lowland | Oceania | Austronesian | Waima | 146.51 | -8.63 | This_study | PASS |
| Be27 | Waima_Bereina | Central | CENTRAL | Bereina | PNG_Lowland | Oceania | Austronesian | Waima | 146.51 | -8.63 | This_study | PASS |
| Be28 | Waima_Bereina | Central | CENTRAL | Bereina | PNG_Lowland | Oceania | Austronesian | Waima | 146.51 | -8.63 | This_study | PASS |
| Be29 | Waima_Bereina | Central | CENTRAL | Bereina | PNG_Lowland | Oceania | Austronesian | Waima | 146.51 | -8.63 | This_study | PASS |

| | | | | | | | | | | | | |
|---|---|---|---|---|---|---|---|---|---|---|---|---|
| Be30 | Waima_Bereina | Central | CENTRAL | Bereina | PNG_Lowland | Oceania | Austronesian | Waima | 146.51 | -8.63 | This_study | PASS |
| Be31 | Waima_Bereina | Central | CENTRAL | Bereina | PNG_Lowland | Oceania | Austronesian | Waima | 146.51 | -8.63 | This_study | PASS |
| Be32 | Waima_Bereina | Central | CENTRAL | Bereina | PNG_Lowland | Oceania | Austronesian | Waima | 146.51 | -8.63 | This_study | PASS |
| Be33 | Waima_Bereina | Central | CENTRAL | Bereina | PNG_Lowland | Oceania | Austronesian | Waima | 146.51 | -8.63 | This_study | PASS |
| Be34 | Waima_Bereina | Central | CENTRAL | Bereina | PNG_Lowland | Oceania | Austronesian | Waima | 146.51 | -8.63 | This_study | PASS |
| papuan6278203 | Hula | Central | CENTRAL | Hood_Peninsula | PNG_Lowland | Oceania | Austronesian | Hula | 147.72 | -10.05 | Bergstrom_et_al | PASS |
| papuan6278314 | Hula | Central | CENTRAL | Hood_Peninsula | PNG_Lowland | Oceania | Austronesian | Hula | 147.72 | -10.05 | Bergstrom_et_al | PASS |
| papuan6278328 | Hula | Central | CENTRAL | Hood_Peninsula | PNG_Lowland | Oceania | Austronesian | Hula | 147.72 | -10.05 | Bergstrom_et_al | PC |
| papuan6278245 | Hula | Central | CENTRAL | Hood_Peninsula | PNG_Lowland | Oceania | Austronesian | Hula | 147.72 | -10.05 | Bergstrom_et_al | PASS |
| papuan6278379 | Hula | Central | CENTRAL | Hood_Peninsula | PNG_Lowland | Oceania | Austronesian | Hula | 147.72 | -10.05 | Bergstrom_et_al | PASS |
| papuan6278322 | Hula | Central | CENTRAL | Hood_Peninsula | PNG_Lowland | Oceania | Austronesian | Hula | 147.72 | -10.05 | Bergstrom_et_al | PASS |
| papuan6278315 | Keapara | Central | CENTRAL | Magarida_Palathaona | PNG_Lowland | Oceania | Austronesian | Keapara | 148.05 | -10.1 | Bergstrom_et_al | PASS |
| papuan6278242 | Keapara | Central | CENTRAL | Magarida_Palathaona | PNG_Lowland | Oceania | Austronesian | Keapara | 148.05 | -10.1 | Bergstrom_et_al | PASS |
| papuan6278227 | Keapara | Central | CENTRAL | Magarida_Palathaona | PNG_Lowland | Oceania | Austronesian | Keapara | 148.05 | -10.1 | Bergstrom_et_al | PASS |
| papuan6278312 | Keapara | Central | CENTRAL | Magarida_Palathaona | PNG_Lowland | Oceania | Austronesian | Keapara | 148.05 | -10.1 | Bergstrom_et_al | PASS |
| papuan6278334 | Motu | Central | CENTRAL | Magarida_Lea-lea | PNG_Lowland | Oceania | Austronesian | Motu | 147.05 | -9.4 | Bergstrom_et_al | PASS |
| papuan6278259 | Motu | Central | CENTRAL | Magarida_Lea-lea | PNG_Lowland | Oceania | Austronesian | Motu | 147.05 | -9.4 | Bergstrom_et_al | Imiss |
| papuan6278212 | Motu | Central | CENTRAL | Magarida_Lea-lea | PNG_Lowland | Oceania | Austronesian | Motu | 147.05 | -9.4 | Bergstrom_et_al | PASS |
| papuan6278324 | Motu | Central | CENTRAL | Magarida_Lea-lea | PNG_Lowland | Oceania | Austronesian | Motu | 147.05 | -9.4 | Bergstrom_et_al | PASS |
| papuan6278213 | Motu | Central | CENTRAL | Magarida_Lea-lea | PNG_Lowland | Oceania | Austronesian | Motu | 147.05 | -9.4 | Bergstrom_et_al | PASS |
| papuan6278211 | Motu | Central | CENTRAL | Magarida_Lea-lea | PNG_Lowland | Oceania | Austronesian | Motu | 147.05 | -9.4 | Bergstrom_et_al | PASS |
| papuan6278360 | Motu | Central | CENTRAL | Magarida_Lea-lea | PNG_Lowland | Oceania | Austronesian | Motu | 147.05 | -9.4 | Bergstrom_et_al | PC |
| papuan6278376 | Motu | Central | CENTRAL | Magarida_Lea-lea | PNG_Lowland | Oceania | Austronesian | Motu | 147.05 | -9.4 | Bergstrom_et_al | PASS |
| papuan6278297 | Motu | Central | CENTRAL | Magarida_Lea-lea | PNG_Lowland | Oceania | Austronesian | Motu | 147.05 | -9.4 | Bergstrom_et_al | PASS |
| papuan6278235 | Motu | Central | CENTRAL | Magarida_Lea-lea | PNG_Lowland | Oceania | Austronesian | Motu | 147.05 | -9.4 | Bergstrom_et_al | PASS |
| papuan6278205 | Motu | Central | CENTRAL | Magarida_Lea-lea | PNG_Lowland | Oceania | Austronesian | Motu | 147.05 | -9.4 | Bergstrom_et_al | PASS |
| papuan6278221 | Motu | Central | CENTRAL | Magarida_Lea-lea | PNG_Lowland | Oceania | Austronesian | Motu | 147.05 | -9.4 | Bergstrom_et_al | PASS |
| papuan6278244 | Motu | Central | CENTRAL | Magarida_Lea-lea | PNG_Lowland | Oceania | Austronesian | Motu | 147.05 | -9.4 | Bergstrom_et_al | PASS |
| papuan6278354 | Motu | Central | CENTRAL | Magarida_Lea-lea | PNG_Lowland | Oceania | Austronesian | Motu | 147.05 | -9.4 | Bergstrom_et_al | PASS |
| papuan6278329 | Motu | Central | CENTRAL | Magarida_Lea-lea | PNG_Lowland | Oceania | Austronesian | Motu | 147.05 | -9.4 | Bergstrom_et_al | PASS |
| papuan6278341 | Sinaugoro | Central | CENTRAL | Rigo | PNG_Lowland | Oceania | Austronesian | Sinaugoro | 147.9 | -9.9 | Bergstrom_et_al | PASS |
| papuan6278302 | Sinaugoro | Central | CENTRAL | Rigo | PNG_Lowland | Oceania | Austronesian | Sinaugoro | 147.9 | -9.9 | Bergstrom_et_al | PASS |
| papuan6278311 | Sinaugoro | Central | CENTRAL | Rigo | PNG_Lowland | Oceania | Austronesian | Sinaugoro | 147.9 | -9.9 | Bergstrom_et_al | PASS |
| papuan6278228 | Waima_Magarida | Central | CENTRAL | Magarida | PNG_Lowland | Oceania | Austronesian | Waima | 146.5 | -8.65 | Bergstrom_et_al | PASS |
| papuan6278331 | Waima_Magarida | Central | CENTRAL | Magarida | PNG_Lowland | Oceania | Austronesian | Waima | 146.5 | -8.65 | Bergstrom_et_al | PASS |
| papuan6278258 | Waima_Magarida | Central | CENTRAL | Magarida | PNG_Lowland | Oceania | Austronesian | Waima | 146.5 | -8.65 | Bergstrom_et_al | PASS |
| papuan6278304 | Waima_Magarida | Central | CENTRAL | Magarida | PNG_Lowland | Oceania | Austronesian | Waima | 146.5 | -8.65 | Bergstrom_et_al | PASS |
| papuan6278069 | Waima_Magarida | Central | CENTRAL | Magarida | PNG_Lowland | Oceania | Austronesian | Waima | 146.5 | -8.65 | Bergstrom_et_al | PASS |
| papuan6278234 | Humene | Central | CENTRAL | Magarida_Tubusereia | PNG_Lowland | Oceania | Trans-New_Guinea | Humene | 147.5 | -9.7 | Bergstrom_et_al | PASS |
| papuan6278446 | Mountain_Koiali | Central | CENTRAL | Port_Moresby | PNG_Lowland | Oceania | Trans-New_Guinea | Mountain_Koiali | 147.45 | -9.05 | Bergstrom_et_al | PASS |
| papuan6278299 | Grass_Koiari | Central | CENTRAL | Magarida | PNG_Lowland | Oceania | Trans-New_Guinea | Grass_Koiari | 147.45 | -9.5 | Bergstrom_et_al | PASS |
| papuan6278349 | Grass_Koiari | Central | CENTRAL | Magarida | PNG_Lowland | Oceania | Trans-New_Guinea | Grass_Koiari | 147.45 | -9.5 | Bergstrom_et_al | PASS |
| papuan6278321 | Grass_Koiari | Central | CENTRAL | Magarida | PNG_Lowland | Oceania | Trans-New_Guinea | Grass_Koiari | 147.45 | -9.5 | Bergstrom_et_al | PASS |
| papuan6278367 | Fuyug | Central | CENTRAL | Goilala | PNG_Lowland | Oceania | Trans-New_Guinea | Fuyug | 147.25 | -8.6 | Bergstrom_et_al | PASS |
| papuan6278342 | Fuyug | Central | CENTRAL | Goilala | PNG_Lowland | Oceania | Trans-New_Guinea | Fuyug | 147.25 | -8.6 | Bergstrom_et_al | PASS |
| papuan6278204 | Tauade | Central | CENTRAL | Goilala | PNG_Lowland | Oceania | Trans-New_Guinea | Tauade | 147.1 | -8.35 | Bergstrom_et_al | PASS |
| papuan6277937 | Chuave | Chimbu | CHIMBU | Chuave | PNG_Highland | Oceania | Trans-New_Guinea | Chuave | 145.1 | -6.2 | Bergstrom_et_al | PASS |
| papuan6277945 | Chuave | Chimbu | CHIMBU | Chuave | PNG_Highland | Oceania | Trans-New_Guinea | Chuave | 145.1 | -6.2 | Bergstrom_et_al | PASS |

| | | | | | | | | | | | | |
|---|---|---|---|---|---|---|---|---|---|---|---|---|
| papuan6277969 | Chuave | Chimbu | CHIMBU | Chuave | PNG_Highland | Oceania | Trans-New_Guinea | Chuave | 145.1 | -6.2 | Bergstrom_et_al | PASS |
| papuan6277939 | Dadibi | Chimbu | CHIMBU | Karimui | PNG_Highland | Oceania | Trans-New_Guinea | Dadibi | 144.6 | -6.5 | Bergstrom_et_al | PASS |
| papuan6277953 | Golin | Chimbu | CHIMBU | Kundiawa_Minono | PNG_Highland | Oceania | Trans-New_Guinea | Golin | 144.85 | -6.15 | Bergstrom_et_al | PASS |
| papuan6277915 | Golin | Chimbu | CHIMBU | Kundiawa_Minono | PNG_Highland | Oceania | Trans-New_Guinea | Golin | 144.85 | -6.15 | Bergstrom_et_al | PASS |
| papuan6278046 | Golin | Chimbu | CHIMBU | Kundiawa_Minono | PNG_Highland | Oceania | Trans-New_Guinea | Golin | 144.85 | -6.15 | Bergstrom_et_al | PASS |
| papuan6277989 | Golin | Chimbu | CHIMBU | Kundiawa_Minono | PNG_Highland | Oceania | Trans-New_Guinea | Golin | 144.85 | -6.15 | Bergstrom_et_al | PASS |
| papuan6278006 | Golin | Chimbu | CHIMBU | Kundiawa_Minono | PNG_Highland | Oceania | Trans-New_Guinea | Golin | 144.85 | -6.15 | Bergstrom_et_al | PASS |
| papuan6277931 | Golin | Chimbu | CHIMBU | Kundiawa_Minono | PNG_Highland | Oceania | Trans-New_Guinea | Golin | 144.85 | -6.15 | Bergstrom_et_al | PASS |
| papuan6277888 | Kuman | Chimbu | CHIMBU | Kundiawa | PNG_Highland | Oceania | Trans-New_Guinea | Kuman | 145 | -5.9 | Bergstrom_et_al | PASS |
| papuan6277818 | Kuman | Chimbu | CHIMBU | Kundiawa | PNG_Highland | Oceania | Trans-New_Guinea | Kuman | 145 | -5.9 | Bergstrom_et_al | PASS |
| papuan6277959 | Kuman | Chimbu | CHIMBU | Kundiawa | PNG_Highland | Oceania | Trans-New_Guinea | Kuman | 145 | -5.9 | Bergstrom_et_al | PASS |
| papuan6277943 | Kuman | Chimbu | CHIMBU | Kundiawa | PNG_Highland | Oceania | Trans-New_Guinea | Kuman | 145 | -5.9 | Bergstrom_et_al | PASS |
| papuan6277935 | Kuman | Chimbu | CHIMBU | Kundiawa | PNG_Highland | Oceania | Trans-New_Guinea | Kuman | 145 | -5.9 | Bergstrom_et_al | PASS |
| papuan6277928 | Kuman | Chimbu | CHIMBU | Kundiawa | PNG_Highland | Oceania | Trans-New_Guinea | Kuman | 145 | -5.9 | Bergstrom_et_al | PASS |
| papuan6277956 | Kuman | Chimbu | CHIMBU | Kundiawa | PNG_Highland | Oceania | Trans-New_Guinea | Kuman | 145 | -5.9 | Bergstrom_et_al | PASS |
| papuan6277933 | Kuman | Chimbu | CHIMBU | Kundiawa | PNG_Highland | Oceania | Trans-New_Guinea | Kuman | 145 | -5.9 | Bergstrom_et_al | PASS |
| papuan6277992 | Kuman | Chimbu | CHIMBU | Kundiawa | PNG_Highland | Oceania | Trans-New_Guinea | Kuman | 145 | -5.9 | Bergstrom_et_al | PASS |
| papuan6277930 | Kuman | Chimbu | CHIMBU | Kundiawa | PNG_Highland | Oceania | Trans-New_Guinea | Kuman | 145 | -5.9 | Bergstrom_et_al | PASS |
| papuan6277970 | Kuman | Chimbu | CHIMBU | Kundiawa | PNG_Highland | Oceania | Trans-New_Guinea | Kuman | 145 | -5.9 | Bergstrom_et_al | PASS |
| papuan6277960 | Kuman | Chimbu | CHIMBU | Kundiawa | PNG_Highland | Oceania | Trans-New_Guinea | Kuman | 145 | -5.9 | Bergstrom_et_al | PASS |
| papuan6278019 | Kuman | Chimbu | CHIMBU | Kundiawa | PNG_Highland | Oceania | Trans-New_Guinea | Kuman | 145 | -5.9 | Bergstrom_et_al | PASS |
| papuan6277984 | Sinasina | Chimbu | CHIMBU | Kundiawa | PNG_Highland | Oceania | Trans-New_Guinea | Sinasina | 145.1 | -6.05 | Bergstrom_et_al | PASS |
| papuan6277968 | Sinasina | Chimbu | CHIMBU | Kundiawa | PNG_Highland | Oceania | Trans-New_Guinea | Sinasina | 145.1 | -6.05 | Bergstrom_et_al | PASS |
| papuan6277921 | Sinasina | Chimbu | CHIMBU | Kundiawa | PNG_Highland | Oceania | Trans-New_Guinea | Sinasina | 145.1 | -6.05 | Bergstrom_et_al | PASS |
| papuan6277977 | Sinasina | Chimbu | CHIMBU | Kundiawa | PNG_Highland | Oceania | Trans-New_Guinea | Sinasina | 145.1 | -6.05 | Bergstrom_et_al | PASS |
| papuan6277938 | Sinasina | Chimbu | CHIMBU | Kundiawa | PNG_Highland | Oceania | Trans-New_Guinea | Sinasina | 145.1 | -6.05 | Bergstrom_et_al | PASS |
| papuan6277996 | Sinasina | Chimbu | CHIMBU | Kundiawa | PNG_Highland | Oceania | Trans-New_Guinea | Sinasina | 145.1 | -6.05 | Bergstrom_et_al | PASS |
| papuan6278004 | Sinasina | Chimbu | CHIMBU | Kundiawa | PNG_Highland | Oceania | Trans-New_Guinea | Sinasina | 145.1 | -6.05 | Bergstrom_et_al | PASS |
| papuan6278062 | Sinasina | Chimbu | CHIMBU | Kundiawa | PNG_Highland | Oceania | Trans-New_Guinea | Sinasina | 145.1 | -6.05 | Bergstrom_et_al | PASS |
| papuan6278014 | Sinasina | Chimbu | CHIMBU | Kundiawa | PNG_Highland | Oceania | Trans-New_Guinea | Sinasina | 145.1 | -6.05 | Bergstrom_et_al | PASS |
| papuan6278524 | Sinasina | Chimbu | CHIMBU | Kundiawa | PNG_Highland | Oceania | Trans-New_Guinea | Sinasina | 145.1 | -6.05 | Bergstrom_et_al | PASS |
| papuan6278332 | Kuanua | East_New_Britain | EAST_NEW_BRITAIN | Rabaul | Bismarck_Arch | Oceania | Austronesian | Kuanua | 152.2 | -4.35 | Bergstrom_et_al | PASS |
| papuan6278210 | Kuanua | East_New_Britain | EAST_NEW_BRITAIN | Rabaul | Bismarck_Arch | Oceania | Austronesian | Kuanua | 152.2 | -4.35 | Bergstrom_et_al | PASS |

| | | | | | | | | | | | | |
|---|---|---|---|---|---|---|---|---|---|---|---|---|
| papuan6278269 | Kuanua | East_New_Britain | EAST_NEW_BRITAIN | Rabaul | Bismarck_Arch | Oceania | Austronesian | Kuanua | 152.2 | -4.35 | Bergstrom_et_al | PASS |
| papuan6278306 | MIXED | MIXED | EAST_NEW_BRITAIN | MIXED | Bismarck_Arch | Oceania | MIXED | MIXED | NA | NA | Bergstrom_et_al | Mixed |
| papuan6278343 | Kairiru | East_Sepik | EAST_SEPIK | Wewak | PNG_Lowland | Oceania | Austronesian | Kairiru | 143.65 | -3.6 | Bergstrom_et_al | PASS |
| papuan6278035 | Ambulas | East_Sepik | EAST_SEPIK | Wewak | PNG_Lowland | Oceania | Lower_Sepik-Ramu | Ambulas | 143.1 | -3.75 | Bergstrom_et_al | PASS |
| papuan6278219 | Ambulas | East_Sepik | EAST_SEPIK | Wewak | PNG_Lowland | Oceania | Lower_Sepik-Ramu | Ambulas | 143.1 | -3.75 | Bergstrom_et_al | PASS |
| papuan6278587 | Angoram | East_Sepik | EAST_SEPIK | Wewak_Yamen | PNG_Lowland | Oceania | Lower_Sepik-Ramu | Angoram | 144.05 | -4.1 | Bergstrom_et_al | PASS |
| papuan6278294 | Angoram | East_Sepik | EAST_SEPIK | Wewak_Yamen | PNG_Lowland | Oceania | Lower_Sepik-Ramu | Angoram | 144.05 | -4.1 | Bergstrom_et_al | PASS |
| papuan6278012 | Boikin | East_Sepik | EAST_SEPIK | Yangoru | PNG_Lowland | Oceania | Lower_Sepik-Ramu | Boikin | 143.3 | -3.23 | Bergstrom_et_al | PASS |
| papuan6278021 | Boikin | East_Sepik | EAST_SEPIK | Yangoru | PNG_Lowland | Oceania | Lower_Sepik-Ramu | Boikin | 143.5 | -3.65 | Bergstrom_et_al | PASS |
| papuan6278289 | Boikin | East_Sepik | EAST_SEPIK | Yangoru | PNG_Lowland | Oceania | Lower_Sepik-Ramu | Boikin | 143.5 | -3.65 | Bergstrom_et_al | PASS |
| papuan6278083 | Boikin | East_Sepik | EAST_SEPIK | Yangoru | PNG_Lowland | Oceania | Lower_Sepik-Ramu | Boikin | 143.5 | -3.65 | Bergstrom_et_al | PASS |
| papuan6278491 | MIXED | MIXED | EAST_SEPIK | MIXED | PNG_Lowland | Oceania | MIXED | MIXED | NA | NA | Bergstrom_et_al | Mixed |
| papuan6278470 | MIXED | MIXED | EAST_SEPIK | MIXED | PNG_Lowland | Oceania | MIXED | MIXED | NA | NA | Bergstrom_et_al | Mixed |
| papuan6278555 | Alekano | Eastern_Highlands | EASTERN_HIGHLANDS | Goroka_Kifamu_Mission | PNG_Highland | Oceania | Trans-New_Guinea | Alekano | 145.35 | -6.05 | Bergstrom_et_al | PASS |
| papuan6278523 | Alekano | Eastern_Highlands | EASTERN_HIGHLANDS | Goroka_Kifamu_Mission | PNG_Highland | Oceania | Trans-New_Guinea | Alekano | 145.35 | -6.05 | Bergstrom_et_al | PASS |
| papuan6278540 | Alekano | Eastern_Highlands | EASTERN_HIGHLANDS | Goroka_Kifamu_Mission | PNG_Highland | Oceania | Trans-New_Guinea | Alekano | 145.35 | -6.05 | Bergstrom_et_al | PASS |
| papuan6278288 | Alekano | Eastern_Highlands | EASTERN_HIGHLANDS | Goroka_Kifamu_Mission | PNG_Highland | Oceania | Trans-New_Guinea | Alekano | 145.35 | -6.05 | Bergstrom_et_al | PASS |
| papuan6278295 | Awiyaana | Eastern_Highlands | EASTERN_HIGHLANDS | Okapa | PNG_Highland | Oceania | Trans-New_Guinea | Awiyaana | 145.8 | -6.6 | Bergstrom_et_al | PASS |
| papuan6278253 | Awiyaana | Eastern_Highlands | EASTERN_HIGHLANDS | Okapa | PNG_Highland | Oceania | Trans-New_Guinea | Awiyaana | 145.8 | -6.6 | Bergstrom_et_al | PASS |
| papuan6278363 | Awiyaana | Eastern_Highlands | EASTERN_HIGHLANDS | Okapa | PNG_Highland | Oceania | Trans-New_Guinea | Awiyaana | 145.8 | -6.6 | Bergstrom_et_al | PASS |
| papuan6278386 | Awiyaana | Eastern_Highlands | EASTERN_HIGHLANDS | Okapa | PNG_Highland | Oceania | Trans-New_Guinea | Awiyaana | 145.8 | -6.6 | Bergstrom_et_al | PASS |
| papuan6278283 | Awiyaana | Eastern_Highlands | EASTERN_HIGHLANDS | Okapa | PNG_Highland | Oceania | Trans-New_Guinea | Awiyaana | 145.8 | -6.6 | Bergstrom_et_al | PASS |
| papuan6278532 | Benabena | Eastern_Highlands | EASTERN_HIGHLANDS | Goroka | PNG_Highland | Oceania | Trans-New_Guinea | Benabena | 145.5 | -6.1 | Bergstrom_et_al | PASS |
| papuan6278601 | Kamano | Eastern_Highlands | EASTERN_HIGHLANDS | Okapa | PNG_Highland | Oceania | Trans-New_Guinea | Kamano | 145.7 | -6.25 | Bergstrom_et_al | PASS |
| papuan6278586 | Kamano | Eastern_Highlands | EASTERN_HIGHLANDS | Okapa | PNG_Highland | Oceania | Trans-New_Guinea | Kamano | 145.7 | -6.25 | Bergstrom_et_al | PASS |
| papuan6278563 | Siane | Eastern_Highlands | EASTERN_HIGHLANDS | Goroka_Watabung | PNG_Highland | Oceania | Trans-New_Guinea | Siane | 145.2 | -6.1 | Bergstrom_et_al | PASS |
| papuan6278539 | Siane | Eastern_Highlands | EASTERN_HIGHLANDS | Goroka_Watabung | PNG_Highland | Oceania | Trans-New_Guinea | Siane | 145.2 | -6.1 | Bergstrom_et_al | PASS |
| papuan6278492 | Siane | Eastern_Highlands | EASTERN_HIGHLANDS | Goroka_Watabung | PNG_Highland | Oceania | Trans-New_Guinea | Siane | 145.2 | -6.1 | Bergstrom_et_al | PASS |
| papuan6278556 | Siane | Eastern_Highlands | EASTERN_HIGHLANDS | Goroka_Watabung | PNG_Highland | Oceania | Trans-New_Guinea | Siane | 145.2 | -6.1 | Bergstrom_et_al | PASS |
| papuan6278338 | Simbari | Eastern_Highlands | EASTERN_HIGHLANDS | Marawaka | PNG_Highland | Oceania | Trans-New_Guinea | Simbari | 145.6 | -7.05 | Bergstrom_et_al | PASS |
| papuan6277940 | Tokano | Eastern_Highlands | EASTERN_HIGHLANDS | Goroka_Watabung | PNG_Highland | Oceania | Trans-New_Guinea | Tokano | 145.25 | -6.05 | Bergstrom_et_al | PASS |
| papuan6278507 | Yagaria | Eastern_Highlands | EASTERN_HIGHLANDS | Goroka | PNG_Highland | Oceania | Trans-New_Guinea | Yagaria | 145.3 | -6.25 | Bergstrom_et_al | PASS |
| papuan6278516 | Yagaria | Eastern_Highlands | EASTERN_HIGHLANDS | Goroka | PNG_Highland | Oceania | Trans-New_Guinea | Yagaria | 145.3 | -6.25 | Bergstrom_et_al | PASS |
| papuan6278515 | Yaweyuha | Eastern_Highlands | EASTERN_HIGHLANDS | Goroka | PNG_Highland | Oceania | Trans-New_Guinea | Yaweyuha | 145.3 | -6.15 | Bergstrom_et_al | PASS |
| papuan6278224 | Yaweyuha | Eastern_Highlands | EASTERN_HIGHLANDS | Goroka | PNG_Highland | Oceania | Trans-New_Guinea | Yaweyuha | 145.3 | -6.15 | Bergstrom_et_al | PASS |
| papuan6278166 | Yipma | Eastern_Highlands | EASTERN_HIGHLANDS | Marawaka | PNG_Highland | Oceania | Trans-New_Guinea | Yipma | 145.8 | -6.9 | Bergstrom_et_al | PASS |

| | | | | | | | | | | | | |
|---|---|---|---|---|---|---|---|---|---|---|---|---|
| papuan6278174 | Yipma | Eastern_Highlands | EASTERN_HIGHLANDS | Marawaka | PNG_Highland | Oceania | Trans-New_Guinea | Yipma | 145.8 | -6.9 | Bergstrom_et_al | PASS |
| papuan6278134 | Yipma | Eastern_Highlands | EASTERN_HIGHLANDS | Marawaka | PNG_Highland | Oceania | Trans-New_Guinea | Yipma | 145.8 | -6.9 | Bergstrom_et_al | PASS |
| papuan6278126 | Yipma | Eastern_Highlands | EASTERN_HIGHLANDS | Marawaka | PNG_Highland | Oceania | Trans-New_Guinea | Yipma | 145.8 | -6.9 | Bergstrom_et_al | PASS |
| papuan6278020 | Enga | Enga | ENGA | Wabag | PNG_Highland | Oceania | Trans-New_Guinea | Enga | 143.6 | -5.5 | Bergstrom_et_al | PASS |
| papuan6278028 | Enga | Enga | ENGA | Wabag | PNG_Highland | Oceania | Trans-New_Guinea | Enga | 143.6 | -5.5 | Bergstrom_et_al | PASS |
| papuan6278042 | Enga | Enga | ENGA | Wabag | PNG_Highland | Oceania | Trans-New_Guinea | Enga | 143.6 | -5.5 | Bergstrom_et_al | PASS |
| papuan6278111 | Enga | Enga | ENGA | Wabag | PNG_Highland | Oceania | Trans-New_Guinea | Enga | 143.6 | -5.5 | Bergstrom_et_al | PASS |
| papuan6278128 | Enga | Enga | ENGA | Wabag | PNG_Highland | Oceania | Trans-New_Guinea | Enga | 143.6 | -5.5 | Bergstrom_et_al | PASS |
| papuan6277902 | Enga | Enga | ENGA | Wabag | PNG_Highland | Oceania | Trans-New_Guinea | Enga | 143.6 | -5.5 | Bergstrom_et_al | PASS |
| papuan6278027 | Enga | Enga | ENGA | Wabag | PNG_Highland | Oceania | Trans-New_Guinea | Enga | 143.6 | -5.5 | Bergstrom_et_al | PASS |
| papuan6278011 | Enga | Enga | ENGA | Wabag | PNG_Highland | Oceania | Trans-New_Guinea | Enga | 143.6 | -5.5 | Bergstrom_et_al | PASS |
| papuan6278382 | Enga | Enga | ENGA | Wabag | PNG_Highland | Oceania | Trans-New_Guinea | Enga | 143.6 | -5.5 | Bergstrom_et_al | PASS |
| papuan6278430 | Enga | Enga | ENGA | Wabag | PNG_Highland | Oceania | Trans-New_Guinea | Enga | 143.6 | -5.5 | Bergstrom_et_al | PASS |
| papuan6278158 | Akoye | Gulf | GULF | Kaberofe | PNG_Highland | Oceania | Trans-New_Guinea | Akoye | 145.7 | -7.65 | Bergstrom_et_al | PASS |
| papuan6278150 | Akoye | Gulf | GULF | Kaberofe | PNG_Highland | Oceania | Trans-New_Guinea | Akoye | 145.7 | -7.65 | Bergstrom_et_al | PASS |
| papuan6278142 | Akoye | Gulf | GULF | Kaberofe | PNG_Highland | Oceania | Trans-New_Guinea | Akoye | 145.7 | -7.65 | Bergstrom_et_al | PASS |
| papuan6278220 | Ikobi | Gulf | GULF | Kerema | PNG_Highland | Oceania | Trans-New_Guinea | Ikobi | 143.5 | -7.1 | Bergstrom_et_al | PASS |
| papuan6278414 | MIXED | MIXED | GULF | MIXED | PNG_Highland | Oceania | MIXED | MIXED | NA | NA | Bergstrom_et_al | Mixed |
| papuan6278390 | MIXED | MIXED | GULF | MIXED | PNG_Highland | Oceania | MIXED | MIXED | NA | NA | Bergstrom_et_al | Mixed |
| papuan6278317 | MIXED | MIXED | GULF | MIXED | PNG_Highland | Oceania | MIXED | MIXED | NA | NA | Bergstrom_et_al | Mixed |
| papuan6278292 | MIXED | MIXED | GULF | MIXED | PNG_Highland | Oceania | MIXED | MIXED | NA | NA | Bergstrom_et_al | Mixed |
| papuan6278366 | Orokolo | Gulf | GULF | Kerema | PNG_Highland | Oceania | Trans-New_Guinea | Orokolo | 145.3 | -7.6 | Bergstrom_et_al | PASS |
| papuan6278236 | Orokolo | Gulf | GULF | Kerema | PNG_Highland | Oceania | Trans-New_Guinea | Orokolo | 145.3 | -7.6 | Bergstrom_et_al | PASS |
| papuan6278387 | Orokolo | Gulf | GULF | Kerema | PNG_Highland | Oceania | Trans-New_Guinea | Orokolo | 145.3 | -7.6 | Bergstrom_et_al | PASS |
| papuan6278347 | Orokolo | Gulf | GULF | Kerema | PNG_Highland | Oceania | Trans-New_Guinea | Orokolo | 145.3 | -7.6 | Bergstrom_et_al | PASS |
| papuan6278381 | Orokolo | Gulf | GULF | Kerema | PNG_Highland | Oceania | Trans-New_Guinea | Orokolo | 145.3 | -7.6 | Bergstrom_et_al | PASS |
| papuan6278274 | Purari_Purari | Gulf | GULF | Puari | PNG_Highland | Oceania | Trans-New_Guinea | Purari | 145 | -7.6 | Bergstrom_et_al | PASS |
| papuan6278251 | Tairuma | Gulf | GULF | Kerema | PNG_Highland | Oceania | Trans-New_Guinea | Tairuma | 145.8 | -7.95 | Bergstrom_et_al | PASS |
| papuan6278327 | Tairuma | Gulf | GULF | Kerema | PNG_Highland | Oceania | Trans-New_Guinea | Tairuma | 145.8 | -7.95 | Bergstrom_et_al | PASS |
| papuan6278243 | Tairuma | Gulf | GULF | Kerema | PNG_Highland | Oceania | Trans-New_Guinea | Tairuma | 145.8 | -7.95 | Bergstrom_et_al | PASS |
| papuan6278319 | Tairuma | Gulf | GULF | Kerema | PNG_Highland | Oceania | Trans-New_Guinea | Tairuma | 145.8 | -7.95 | Bergstrom_et_al | PASS |
| papuan6278303 | Tairuma | Gulf | GULF | Kerema | PNG_Highland | Oceania | Trans-New_Guinea | Tairuma | 145.8 | -7.95 | Bergstrom_et_al | PASS |
| papuan6278316 | Tairuma | Gulf | GULF | Kerema | PNG_Highland | Oceania | Trans-New_Guinea | Tairuma | 145.8 | -7.95 | Bergstrom_et_al | PASS |

| | | | | | | | | | | | | |
|---|---|---|---|---|---|---|---|---|---|---|---|---|
| papuan6278300 | Toaripi | Gulf | GULF | Kerema | PNG_Highland | Oceania | Trans-New_Guinea | Toaripi | 146.2 | -8.1 | Bergstrom_et_al | PASS |
| papuan6278260 | Toaripi | Gulf | GULF | Kerema | PNG_Highland | Oceania | Trans-New_Guinea | Toaripi | 146.2 | -8.1 | Bergstrom_et_al | PASS |
| papuan6278398 | Toaripi | Gulf | GULF | Kerema | PNG_Highland | Oceania | Trans-New_Guinea | Toaripi | 146.2 | -8.1 | Bergstrom_et_al | PASS |
| papuan6278229 | Toaripi | Gulf | GULF | Kerema | PNG_Highland | Oceania | Trans-New_Guinea | Toaripi | 146.2 | -8.1 | Bergstrom_et_al | PASS |
| papuan6278368 | Toaripi | Gulf | GULF | Kerema | PNG_Highland | Oceania | Trans-New_Guinea | Toaripi | 146.2 | -8.1 | Bergstrom_et_al | PASS |
| papuan6278369 | Toaripi | Gulf | GULF | Kerema | PNG_Highland | Oceania | Trans-New_Guinea | Toaripi | 146.2 | -8.1 | Bergstrom_et_al | PASS |
| papuan6278374 | Toaripi | Gulf | GULF | Kerema | PNG_Highland | Oceania | Trans-New_Guinea | Toaripi | 146.2 | -8.1 | Bergstrom_et_al | PASS |
| papuan6278546 | Takia | Madang_Lowland | MADANG | Badilu | PNG_Lowland | Oceania | Austronesian | Takia | 146 | -4.7 | Bergstrom_et_al | PASS |
| papuan6277971 | Aruamu | Madang_Lowland | MADANG | Bogia | PNG_Lowland | Oceania | Lower_Sepik-Ramu | Aruamu | 144.85 | -4.3 | Bergstrom_et_al | PASS |
| papuan6278603 | Giri | Madang_Lowland | MADANG | Garati | PNG_Lowland | Oceania | Lower_Sepik-Ramu | Giri | 144.7 | -4.25 | Bergstrom_et_al | PASS |
| papuan6278596 | Kominimung | Madang_Lowland | MADANG | Madang | PNG_Lowland | Oceania | Lower_Sepik-Ramu | Kominimung | 144.75 | -4.7 | Bergstrom_et_al | PASS |
| papuan6278525 | Rao | Madang_Lowland | MADANG | Madang | PNG_Lowland | Oceania | Lower_Sepik-Ramu | Rao | 144.5 | -4.8 | Bergstrom_et_al | PASS |
| papuan6278569 | Rao | Madang_Lowland | MADANG | Madang | PNG_Lowland | Oceania | Lower_Sepik-Ramu | Rao | 144.5 | -4.8 | Bergstrom_et_al | PASS |
| papuan6278593 | MIXED | MIXED | MADANG | MIXED | PNG_Lowland | Oceania | MIXED | MIXED | NA | NA | Bergstrom_et_al | Mixed |
| papuan6278529 | MIXED | MIXED | MADANG | MIXED | PNG_Lowland | Oceania | MIXED | MIXED | NA | NA | Bergstrom_et_al | Mixed |
| papuan6278565 | MIXED | MIXED | MADANG | MIXED | PNG_Lowland | Oceania | MIXED | MIXED | NA | NA | Bergstrom_et_al | Mixed |
| papuan6278583 | MIXED | MIXED | MADANG | MIXED | PNG_Lowland | Oceania | MIXED | MIXED | NA | NA | Bergstrom_et_al | Mixed |
| papuan6278561 | MIXED | MIXED | MADANG | MIXED | PNG_Lowland | Oceania | MIXED | MIXED | NA | NA | Bergstrom_et_al | Mixed |
| papuan6278584 | MIXED | MIXED | MADANG | MIXED | PNG_Lowland | Oceania | MIXED | MIXED | NA | NA | Bergstrom_et_al | Mixed |
| papuan6278557 | Amaimon | Madang_Lowland | MADANG | Madang | PNG_Lowland | Oceania | Trans-New_Guinea | Amaimon | 145.37 | -5.2 | Bergstrom_et_al | PASS |
| papuan6277941 | Amele | Madang_Lowland | MADANG | Utu_Mission | PNG_Lowland | Oceania | Trans-New_Guinea | Amele | 145.7 | -5.25 | Bergstrom_et_al | PASS |
| papuan6278592 | Amele | Madang_Lowland | MADANG | Utu_Mission | PNG_Lowland | Oceania | Trans-New_Guinea | Amele | 145.7 | -5.25 | Bergstrom_et_al | PASS |
| papuan6278553 | Bemal | Madang_Lowland | MADANG | Trans-Gogol | PNG_Lowland | Oceania | Trans-New_Guinea | Bemal | 145.45 | -5.3 | Bergstrom_et_al | PASS |
| papuan6277872 | Biyom | Madang_Highland | MADANG | Brahmin | PNG_Highland | Oceania | Trans-New_Guinea | Biyom | 145.45 | -5.85 | Bergstrom_et_al | PASS |
| papuan6277918 | Gende | Madang_Highland | MADANG | Bundi | PNG_Highland | Oceania | Trans-New_Guinea | Gende | 145.15 | -5.7 | Bergstrom_et_al | PASS |
| papuan6277910 | Gende | Madang_Highland | MADANG | Bundi | PNG_Highland | Oceania | Trans-New_Guinea | Gende | 145.15 | -5.7 | Bergstrom_et_al | PASS |
| papuan6277885 | Gende | Madang_Highland | MADANG | Bundi | PNG_Highland | Oceania | Trans-New_Guinea | Gende | 145.15 | -5.7 | Bergstrom_et_al | PASS |
| papuan6277906 | Gende | Madang_Highland | MADANG | Bundi | PNG_Highland | Oceania | Trans-New_Guinea | Gende | 145.15 | -5.7 | Bergstrom_et_al | PASS |
| papuan6277897 | Gende | Madang_Highland | MADANG | Bundi | PNG_Highland | Oceania | Trans-New_Guinea | Gende | 145.15 | -5.7 | Bergstrom_et_al | PASS |
| papuan6277881 | Gende | Madang_Highland | MADANG | Bundi | PNG_Highland | Oceania | Trans-New_Guinea | Gende | 145.15 | -5.7 | Bergstrom_et_al | PASS |
| papuan6277873 | Gende | Madang_Highland | MADANG | Bundi | PNG_Highland | Oceania | Trans-New_Guinea | Gende | 145.15 | -5.7 | Bergstrom_et_al | PASS |
| papuan6277967 | Gende | Madang_Highland | MADANG | Bundi | PNG_Highland | Oceania | Trans-New_Guinea | Gende | 145.15 | -5.7 | Bergstrom_et_al | PASS |
| papuan6277868 | Gende | Madang_Highland | MADANG | Bundi | PNG_Highland | Oceania | Trans-New_Guinea | Gende | 145.15 | -5.7 | Bergstrom_et_al | PASS |
| papuan6277883 | Gende | Madang_Highland | MADANG | Bundi | PNG_Highland | Oceania | Trans-New_Guinea | Gende | 145.15 | -5.7 | Bergstrom_et_al | PASS |
| papuan6277934 | Gende | Madang_Highland | MADANG | Bundi | PNG_Highland | Oceania | Trans-New_Guinea | Gende | 145.15 | -5.7 | Bergstrom_et_al | PASS |
| papuan6277907 | Gende | Madang_Highland | MADANG | Bundi | PNG_Highland | Oceania | Trans-New_Guinea | Gende | 145.15 | -5.7 | Bergstrom_et_al | PASS |
| papuan6277849 | Gende | Madang_Highland | MADANG | Bundi | PNG_Highland | Oceania | Trans-New_Guinea | Gende | 145.15 | -5.7 | Bergstrom_et_al | PASS |
| papuan6277911 | Gende | Madang_Highland | MADANG | Bundi | PNG_Highland | Oceania | Trans-New_Guinea | Gende | 145.15 | -5.7 | Bergstrom_et_al | PASS |

| | | | | | | | | | | | | |
|---|---|---|---|---|---|---|---|---|---|---|---|---|
| papuan6277889 | Gende | Madang_Highland | MADANG | Bundi | PNG_Highland | Oceania | Trans-New_Guinea | Gende | 145.15 | -5.7 | Bergstrom_et_al | PASS |
| papuan6277899 | Gende | Madang_Highland | MADANG | Bundi | PNG_Highland | Oceania | Trans-New_Guinea | Gende | 145.15 | -5.7 | Bergstrom_et_al | PASS |
| papuan6277866 | Gende | Madang_Highland | MADANG | Bundi | PNG_Highland | Oceania | Trans-New_Guinea | Gende | 145.15 | -5.7 | Bergstrom_et_al | PASS |
| papuan6277890 | Gende | Madang_Highland | MADANG | Bundi | PNG_Highland | Oceania | Trans-New_Guinea | Gende | 145.15 | -5.7 | Bergstrom_et_al | PASS |
| papuan6277898 | Gende | Madang_Highland | MADANG | Bundi | PNG_Highland | Oceania | Trans-New_Guinea | Gende | 145.15 | -5.7 | Bergstrom_et_al | PASS |
| papuan6277859 | Gende | Madang_Highland | MADANG | Bundi | PNG_Highland | Oceania | Trans-New_Guinea | Gende | 145.15 | -5.7 | Bergstrom_et_al | PASS |
| papuan6277827 | Gende | Madang_Highland | MADANG | Bundi | PNG_Highland | Oceania | Trans-New_Guinea | Gende | 145.15 | -5.7 | Bergstrom_et_al | PASS |
| papuan6277820 | Gende | Madang_Highland | MADANG | Bundi | PNG_Highland | Oceania | Trans-New_Guinea | Gende | 145.15 | -5.7 | Bergstrom_et_al | Kin |
| papuan6277836 | Gende | Madang_Highland | MADANG | Bundi | PNG_Highland | Oceania | Trans-New_Guinea | Gende | 145.15 | -5.7 | Bergstrom_et_al | PASS |
| papuan6277844 | Gende | Madang_Highland | MADANG | Bundi | PNG_Highland | Oceania | Trans-New_Guinea | Gende | 145.15 | -5.7 | Bergstrom_et_al | PASS |
| papuan6277852 | Gende | Madang_Highland | MADANG | Bundi | PNG_Highland | Oceania | Trans-New_Guinea | Gende | 145.15 | -5.7 | Bergstrom_et_al | PASS |
| papuan6277884 | Gende | Madang_Highland | MADANG | Bundi | PNG_Highland | Oceania | Trans-New_Guinea | Gende | 145.15 | -5.7 | Bergstrom_et_al | PASS |
| papuan6277876 | Gende | Madang_Highland | MADANG | Bundi | PNG_Highland | Oceania | Trans-New_Guinea | Gende | 145.15 | -5.7 | Bergstrom_et_al | PASS |
| papuan6277864 | Gende | Madang_Highland | MADANG | Bundi | PNG_Highland | Oceania | Trans-New_Guinea | Gende | 145.15 | -5.7 | Bergstrom_et_al | PASS |
| papuan6277905 | Gende | Madang_Highland | MADANG | Bundi | PNG_Highland | Oceania | Trans-New_Guinea | Gende | 145.15 | -5.7 | Bergstrom_et_al | PASS |
| papuan6277817 | Gende | Madang_Highland | MADANG | Bundi | PNG_Highland | Oceania | Trans-New_Guinea | Gende | 145.15 | -5.7 | Bergstrom_et_al | PASS |
| papuan6277874 | Gende | Madang_Highland | MADANG | Bundi | PNG_Highland | Oceania | Trans-New_Guinea | Gende | 145.15 | -5.7 | Bergstrom_et_al | PASS |
| papuan6277882 | Gende | Madang_Highland | MADANG | Bundi | PNG_Highland | Oceania | Trans-New_Guinea | Gende | 145.15 | -5.7 | Bergstrom_et_al | PASS |
| papuan6277919 | Gende | Madang_Highland | MADANG | Bundi | PNG_Highland | Oceania | Trans-New_Guinea | Gende | 145.15 | -5.7 | Bergstrom_et_al | Kin |
| papuan6277974 | Gende | Madang_Highland | MADANG | Bundi | PNG_Highland | Oceania | Trans-New_Guinea | Gende | 145.15 | -5.7 | Bergstrom_et_al | PASS |
| papuan6277966 | Gende | Madang_Highland | MADANG | Bundi | PNG_Highland | Oceania | Trans-New_Guinea | Gende | 145.15 | -5.7 | Bergstrom_et_al | Kin |
| papuan6277950 | Gende | Madang_Highland | MADANG | Bundi | PNG_Highland | Oceania | Trans-New_Guinea | Gende | 145.15 | -5.7 | Bergstrom_et_al | PASS |
| papuan6277999 | Gende | Madang_Highland | MADANG | Bundi | PNG_Highland | Oceania | Trans-New_Guinea | Gende | 145.15 | -5.7 | Bergstrom_et_al | PASS |
| papuan6277991 | Gende | Madang_Highland | MADANG | Bundi | PNG_Highland | Oceania | Trans-New_Guinea | Gende | 145.15 | -5.7 | Bergstrom_et_al | PASS |
| papuan6277826 | Gende | Madang_Highland | MADANG | Bundi | PNG_Highland | Oceania | Trans-New_Guinea | Gende | 145.15 | -5.7 | Bergstrom_et_al | PASS |
| papuan6277828 | Gende | Madang_Highland | MADANG | Bundi | PNG_Highland | Oceania | Trans-New_Guinea | Gende | 145.15 | -5.7 | Bergstrom_et_al | PASS |
| papuan6277819 | Gende | Madang_Highland | MADANG | Bundi | PNG_Highland | Oceania | Trans-New_Guinea | Gende | 145.15 | -5.7 | Bergstrom_et_al | PASS |
| papuan6277891 | Gende | Madang_Highland | MADANG | Bundi | PNG_Highland | Oceania | Trans-New_Guinea | Gende | 145.15 | -5.7 | Bergstrom_et_al | PASS |
| papuan6277829 | Gende | Madang_Highland | MADANG | Bundi | PNG_Highland | Oceania | Trans-New_Guinea | Gende | 145.15 | -5.7 | Bergstrom_et_al | PASS |
| papuan6278607 | Gende | Madang_Highland | MADANG | Bundi | PNG_Highland | Oceania | Trans-New_Guinea | Gende | 145.15 | -5.7 | Bergstrom_et_al | PASS |
| papuan6278541 | MIXED | Madang_Lowland | MADANG | MIXED | PNG_Lowland | Oceania | Trans-New_Guinea | MIXED | 145.65 | -5.2 | Bergstrom_et_al | PASS |
| papuan6278577 | Korak | Madang_Lowland | MADANG | Rempi_Mission | PNG_Lowland | Oceania | Trans-New_Guinea | Korak | 145.45 | -4.55 | Bergstrom_et_al | PASS |
| papuan6278533 | Nobonob | Madang_Lowland | MADANG | Rempi_Mission | PNG_Lowland | Oceania | Trans-New_Guinea | Nobonob | 145.7 | -5.1 | Bergstrom_et_al | PASS |
| papuan6278509 | Nobonob | Madang_Lowland | MADANG | Rempi_Mission | PNG_Lowland | Oceania | Trans-New_Guinea | Nobonob | 145.7 | -5.1 | Bergstrom_et_al | PASS |

| | | | | | | | | | | | | |
|---|---|---|---|---|---|---|---|---|---|---|---|---|
| papuan6278597 | MIXED | Madang_Lowland | MADANG | MIXED | PNG_Lowland | Oceania | Trans-New_Guinea | MIXED | NA | NA | Bergstrom_et_al | PASS |
| papuan6278582 | Pondoma | Madang_Lowland | MADANG | Josephstaal | PNG_Lowland | Oceania | Trans-New_Guinea | Pondoma | 145 | -4.75 | Bergstrom_et_al | PASS |
| papuan6278506 | MIXED | Madang_Lowland | MADANG | MIXED | PNG_Lowland | Oceania | Trans-New_Guinea | MIXED | NA | NA | Bergstrom_et_al | PASS |
| papuan6278606 | Sop | Madang_Lowland | MADANG | Madang | PNG_Lowland | Oceania | Trans-New_Guinea | Sop | 145.45 | -5.5 | Bergstrom_et_al | PASS |
| papuan6278564 | Wagi | Madang_Lowland | MADANG | Rempi_Mission | PNG_Lowland | Oceania | Trans-New_Guinea | Wagi | 145.75 | -5.2 | Bergstrom_et_al | PASS |
| papuan6278572 | Wagi | Madang_Lowland | MADANG | Rempi_Mission | PNG_Lowland | Oceania | Trans-New_Guinea | Wagi | 145.75 | -5.2 | Bergstrom_et_al | PASS |
| papuan6278580 | Wagi | Madang_Lowland | MADANG | Rempi_Mission | PNG_Lowland | Oceania | Trans-New_Guinea | Wagi | 145.75 | -5.2 | Bergstrom_et_al | PASS |
| papuan6278493 | Wagi | Madang_Lowland | MADANG | Rempi_Mission | PNG_Lowland | Oceania | Trans-New_Guinea | Wagi | 145.75 | -5.2 | Bergstrom_et_al | PASS |
| papuan6278501 | Wagi | Madang_Lowland | MADANG | Rempi_Mission | PNG_Lowland | Oceania | Trans-New_Guinea | Wagi | 145.75 | -5.2 | Bergstrom_et_al | PASS |
| papuan6278589 | Wagi | Madang_Lowland | MADANG | Rempi_Mission | PNG_Lowland | Oceania | Trans-New_Guinea | Wagi | 145.75 | -5.2 | Bergstrom_et_al | PASS |
| papuan6278273 | Tulu-Bohuai | Manus_New_Ireland | MANUS | Lorengau | Bismarck_Arch | Oceania | Austronesian | Tulu-Bohuai | 146.85 | -2.1 | Bergstrom_et_al | PASS |
| papuan6278301 | MIXED | MIXED | MANUS | MIXED | Bismarck_Arch | Oceania | MIXED | MIXED | NA | NA | Bergstrom_et_al | Mixed |
| papuan6278320 | MIXED | MIXED | MANUS | MIXED | Bismarck_Arch | Oceania | MIXED | MIXED | NA | NA | Bergstrom_et_al | Mixed |
| papuan6278513 | MIXED | Misima | MILNE_BAY | MIXED | Massim | Oceania | Austronesian | MIXED | NA | NA | Bergstrom_et_al | PASS |
| papuan6278277 | Dobu | Fergusson | MILNE_BAY | Esaala | Massim | Oceania | Austronesian | Dobu | 151.26 | -9.95 | Bergstrom_et_al | PASS |
| papuan6278358 | MIXED | MIXED | MILNE_BAY | MIXED | Massim | Oceania | MIXED | MIXED | NA | NA | Bergstrom_et_al | Mixed |
| papuan6278252 | MIXED | MIXED | MILNE_BAY | MIXED | Massim | Oceania | MIXED | MIXED | NA | NA | Bergstrom_et_al | Mixed |
| papuan6278378 | MIXED | MIXED | MILNE_BAY | MIXED | Massim | Oceania | MIXED | MIXED | NA | NA | Bergstrom_et_al | Mixed |
| papuan6278052 | Umanakaina | Mainland_Eastern_Tip | MILNE_BAY | Rabaraba | Massim | Oceania | Trans-New_Guinea | Umanakaina | 149.6 | -9.9 | Bergstrom_et_al | PASS |
| papuan6278508 | MIXED | MIXED | MIXED | MIXED | MIXED | Oceania | Trans-New_Guinea | MIXED | NA | NA | Bergstrom_et_al | Mixed |
| papuan6277870 | MIXED | MIXED | MIXED | MIXED | MIXED | Oceania | Trans-New_Guinea | MIXED | NA | NA | Bergstrom_et_al | Mixed |
| papuan6278517 | MIXED | MIXED | MIXED | MIXED | MIXED | Oceania | Austronesian | MIXED | NA | NA | Bergstrom_et_al | Mixed |
| papuan6278514 | MIXED | MIXED | MIXED | MIXED | MIXED | Oceania | Trans-New_Guinea | MIXED | NA | NA | Bergstrom_et_al | Mixed |
| papuan6278591 | MIXED | MIXED | MIXED | MIXED | MIXED | Oceania | MIXED | MIXED | NA | NA | Bergstrom_et_al | Mixed |
| papuan6278498 | MIXED | MIXED | MIXED | MIXED | MIXED | Oceania | MIXED | MIXED | NA | NA | Bergstrom_et_al | Mixed |
| papuan6278537 | MIXED | MIXED | MIXED | MIXED | MIXED | Oceania | Austronesian | MIXED | NA | NA | Bergstrom_et_al | Mixed |
| papuan6278595 | MIXED | MIXED | MIXED | MIXED | MIXED | Oceania | MIXED | MIXED | NA | NA | Bergstrom_et_al | Mixed |
| papuan6278588 | MIXED | MIXED | MIXED | MIXED | MIXED | Oceania | Lower_Sepik-Ramu | MIXED | NA | NA | Bergstrom_et_al | Mixed |
| papuan6278059 | MIXED | MIXED | MIXED | MIXED | MIXED | Oceania | Trans-New_Guinea | MIXED | 144.45 | -6.9 | Bergstrom_et_al | Mixed |
| papuan6278350 | Bugawac | Morobe | MOROBE | Lae | PNG_Lowland | Oceania | Austronesian | Bugawac | 147.25 | -6.7 | Bergstrom_et_al | PASS |
| papuan6277843 | Yabem | Morobe | MOROBE | Finschhafen | PNG_Lowland | Oceania | Austronesian | Yabem | 147.85 | -6.65 | Bergstrom_et_al | PASS |
| papuan6278202 | Yabem | Morobe | MOROBE | Finschhafen | PNG_Lowland | Oceania | Austronesian | Yabem | 147.85 | -6.65 | Bergstrom_et_al | PASS |
| papuan6278099 | MIXED | MIXED | MOROBE | MIXED | PNG_Lowland | Oceania | MIXED | MIXED | NA | NA | Bergstrom_et_al | Mixed |
| papuan6278308 | MIXED | MIXED | MOROBE | MIXED | PNG_Lowland | Oceania | MIXED | MIXED | NA | NA | Bergstrom_et_al | Mixed |
| papuan6278370 | MIXED | MIXED | MOROBE | MIXED | PNG_Lowland | Oceania | MIXED | MIXED | NA | NA | Bergstrom_et_al | Mixed |
| papuan6278585 | Kate | Morobe | MOROBE | Finschhafen | PNG_Lowland | Oceania | Trans-New_Guinea | Kate | 147.7 | -6.45 | Bergstrom_et_al | PASS |
| papuan6278573 | Kate | Morobe | MOROBE | Finschhafen | PNG_Lowland | Oceania | Trans-New_Guinea | Kate | 147.7 | -6.45 | Bergstrom_et_al | PASS |
| papuan6278036 | Nabak | Morobe | MOROBE | Lae | PNG_Lowland | Oceania | Trans-New_Guinea | Nabak | 147 | -6.5 | Bergstrom_et_al | PASS |
| papuan6278530 | Patpatar | Manus_New_Ireland | NEW_IRELAND | Namatanai | Bismarck_Arch | Oceania | Austronesian | Patpatar | 152.5 | -3.75 | Bergstrom_et_al | PASS |
| papuan6278067 | Kara | Manus_New_Ireland | NEW_IRELAND | northern_New_Ireland | Bismarck_Arch | Oceania | Austronesian | Kara | 151.1 | -2.8 | Bergstrom_et_al | PASS |
| papuan6278454 | MIXED | MIXED | NORTHERN | MIXED | Massim | Oceania | MIXED | MIXED | NA | NA | Bergstrom_et_al | Mixed |
| papuan6278357 | Orokaiva | Northern | NORTHERN | Popondetta | Massim | Oceania | Trans-New_Guinea | Orokaiva | 148.2 | -8.75 | Bergstrom_et_al | PASS |
| papuan6278384 | Korafe | Northern | NORTHERN | Popondetta | Massim | Oceania | Trans-New_Guinea | Korafe | 149.25 | -9.05 | Bergstrom_et_al | PASS |
| papuan6278313 | Korafe | Northern | NORTHERN | Popondetta | Massim | Oceania | Trans-New_Guinea | Korafe | 149.25 | -9.05 | Bergstrom_et_al | PASS |
| papuan6278284 | Korafe | Northern | NORTHERN | Popondetta | Massim | Oceania | Trans-New_Guinea | Korafe | 149.25 | -9.05 | Bergstrom_et_al | PASS |

| | | | | | | | | | | | | |
|---|---|---|---|---|---|---|---|---|---|---|---|---|
| papuan6278026 | Angal | Southern_Highlands | SOUTHERN_HIGHLANDS | Mendi | PNG_Highland | Oceania | Trans-New_Guinea | Angal | 143.6 | -6 | Bergstrom_et_al | PASS |
| papuan6278039 | Angal | Southern_Highlands | SOUTHERN_HIGHLANDS | Mendi | PNG_Highland | Oceania | Trans-New_Guinea | Angal | 143.6 | -6 | Bergstrom_et_al | PASS |
| papuan6278047 | Angal | Southern_Highlands | SOUTHERN_HIGHLANDS | Mendi | PNG_Highland | Oceania | Trans-New_Guinea | Angal | 143.6 | -6 | Bergstrom_et_al | PASS |
| papuan6278073 | Angal | Southern_Highlands | SOUTHERN_HIGHLANDS | Mendi | PNG_Highland | Oceania | Trans-New_Guinea | Angal | 143.6 | -6 | Bergstrom_et_al | PASS |
| papuan6278097 | Angal | Southern_Highlands | SOUTHERN_HIGHLANDS | Mendi | PNG_Highland | Oceania | Trans-New_Guinea | Angal | 143.6 | -6 | Bergstrom_et_al | PASS |
| papuan6278074 | Angal | Southern_Highlands | SOUTHERN_HIGHLANDS | Mendi | PNG_Highland | Oceania | Trans-New_Guinea | Angal | 143.6 | -6 | Bergstrom_et_al | PASS |
| papuan6278023 | Angal | Southern_Highlands | SOUTHERN_HIGHLANDS | Mendi | PNG_Highland | Oceania | Trans-New_Guinea | Angal | 143.6 | -6 | Bergstrom_et_al | PASS |
| papuan6278209 | Angal | Southern_Highlands | SOUTHERN_HIGHLANDS | Mendi | PNG_Highland | Oceania | Trans-New_Guinea | Angal | 143.6 | -6 | Bergstrom_et_al | PASS |
| papuan6278255 | East_Kewa_Aiya | Southern_Highlands | SOUTHERN_HIGHLANDS | Ialibu | PNG_Highland | Oceania | Trans-New_Guinea | East_Kewa_Aiya | 143.9 | -6.4 | Bergstrom_et_al | PASS |
| papuan6278156 | East_Kewa_Aiya | Southern_Highlands | SOUTHERN_HIGHLANDS | Ialibu | PNG_Highland | Oceania | Trans-New_Guinea | East_Kewa_Aiya | 143.9 | -6.4 | Bergstrom_et_al | PASS |
| papuan6278164 | East_Kewa_Aiya | Southern_Highlands | SOUTHERN_HIGHLANDS | Ialibu | PNG_Highland | Oceania | Trans-New_Guinea | East_Kewa_Aiya | 143.9 | -6.4 | Bergstrom_et_al | PASS |
| papuan6278230 | East_Kewa_Aiya | Southern_Highlands | SOUTHERN_HIGHLANDS | Ialibu | PNG_Highland | Oceania | Trans-New_Guinea | East_Kewa_Aiya | 143.9 | -6.4 | Bergstrom_et_al | PASS |
| papuan6278238 | East_Kewa_Aiya | Southern_Highlands | SOUTHERN_HIGHLANDS | Ialibu | PNG_Highland | Oceania | Trans-New_Guinea | East_Kewa_Aiya | 143.9 | -6.4 | Bergstrom_et_al | PASS |
| papuan6278272 | East_Kewa_Aiya | Southern_Highlands | SOUTHERN_HIGHLANDS | Ialibu | PNG_Highland | Oceania | Trans-New_Guinea | East_Kewa_Aiya | 143.9 | -6.4 | Bergstrom_et_al | PASS |
| papuan6278287 | East_Kewa_Aiya | Southern_Highlands | SOUTHERN_HIGHLANDS | Ialibu | PNG_Highland | Oceania | Trans-New_Guinea | East_Kewa_Aiya | 143.9 | -6.4 | Bergstrom_et_al | PASS |
| papuan6277840 | East_Kewa_Aliya | Southern_Highlands | SOUTHERN_HIGHLANDS | Ialibu | PNG_Highland | Oceania | Trans-New_Guinea | East_Kewa_Aliya | 143.91 | -6.41 | Bergstrom_et_al | PASS |
| papuan6278125 | East_Kewa_Aliya | Southern_Highlands | SOUTHERN_HIGHLANDS | Ialibu | PNG_Highland | Oceania | Trans-New_Guinea | East_Kewa_Aliya | 143.91 | -6.41 | Bergstrom_et_al | PASS |
| papuan6278141 | East_Kewa_Aliya | Southern_Highlands | SOUTHERN_HIGHLANDS | Ialibu | PNG_Highland | Oceania | Trans-New_Guinea | East_Kewa_Aliya | 143.91 | -6.41 | Bergstrom_et_al | PASS |
| papuan6277848 | East_Kewa_Aliya | Southern_Highlands | SOUTHERN_HIGHLANDS | Ialibu | PNG_Highland | Oceania | Trans-New_Guinea | East_Kewa_Aliya | 143.91 | -6.41 | Bergstrom_et_al | PASS |
| papuan6277856 | East_Kewa_Aliya | Southern_Highlands | SOUTHERN_HIGHLANDS | Ialibu | PNG_Highland | Oceania | Trans-New_Guinea | East_Kewa_Aliya | 143.91 | -6.41 | Bergstrom_et_al | PASS |
| papuan6278181 | East_Kewa_Aliya | Southern_Highlands | SOUTHERN_HIGHLANDS | Ialibu | PNG_Highland | Oceania | Trans-New_Guinea | East_Kewa_Aliya | 143.91 | -6.41 | Bergstrom_et_al | PASS |
| papuan6278222 | East_Kewa_Aliya | Southern_Highlands | SOUTHERN_HIGHLANDS | Ialibu | PNG_Highland | Oceania | Trans-New_Guinea | East_Kewa_Aliya | 143.91 | -6.41 | Bergstrom_et_al | PASS |
| papuan6278286 | East_Kewa_Aliya | Southern_Highlands | SOUTHERN_HIGHLANDS | Ialibu | PNG_Highland | Oceania | Trans-New_Guinea | East_Kewa_Aliya | 143.91 | -6.41 | Bergstrom_et_al | PASS |
| papuan6278215 | East_Kewa_Aliya | Southern_Highlands | SOUTHERN_HIGHLANDS | Ialibu | PNG_Highland | Oceania | Trans-New_Guinea | East_Kewa_Aliya | 143.91 | -6.41 | Bergstrom_et_al | PASS |
| papuan6278264 | East_Kewa_Aliya | Southern_Highlands | SOUTHERN_HIGHLANDS | Ialibu | PNG_Highland | Oceania | Trans-New_Guinea | East_Kewa_Aliya | 143.91 | -6.41 | Bergstrom_et_al | PASS |
| papuan6278048 | Erave | Southern_Highlands | SOUTHERN_HIGHLANDS | Mendi | PNG_Highland | Oceania | Trans-New_Guinea | Erave | 143.95 | -6.6 | Bergstrom_et_al | PASS |
| papuan6278138 | Erave | Southern_Highlands | SOUTHERN_HIGHLANDS | Mendi | PNG_Highland | Oceania | Trans-New_Guinea | Erave | 143.95 | -6.6 | Bergstrom_et_al | PASS |
| papuan6278122 | Erave | Southern_Highlands | SOUTHERN_HIGHLANDS | Mendi | PNG_Highland | Oceania | Trans-New_Guinea | Erave | 143.95 | -6.6 | Bergstrom_et_al | PASS |
| papuan6278612 | Erave | Southern_Highlands | SOUTHERN_HIGHLANDS | Mendi | PNG_Highland | Oceania | Trans-New_Guinea | Erave | 143.95 | -6.6 | Bergstrom_et_al | Kin |
| papuan6278123 | Erave | Southern_Highlands | SOUTHERN_HIGHLANDS | Mendi | PNG_Highland | Oceania | Trans-New_Guinea | Erave | 143.95 | -6.6 | Bergstrom_et_al | PASS |
| papuan6278131 | Erave | Southern_Highlands | SOUTHERN_HIGHLANDS | Mendi | PNG_Highland | Oceania | Trans-New_Guinea | Erave | 143.95 | -6.6 | Bergstrom_et_al | PASS |
| papuan6278147 | Erave | Southern_Highlands | SOUTHERN_HIGHLANDS | Mendi | PNG_Highland | Oceania | Trans-New_Guinea | Erave | 143.95 | -6.6 | Bergstrom_et_al | PASS |
| papuan6278187 | Erave | Southern_Highlands | SOUTHERN_HIGHLANDS | Mendi | PNG_Highland | Oceania | Trans-New_Guinea | Erave | 143.95 | -6.6 | Bergstrom_et_al | PASS |

| | | | | | | | | | | | | |
|---|---|---|---|---|---|---|---|---|---|---|---|---|
| papuan6278140 | Erave | Southern_Highlands | SOUTHERN_HIGHLANDS | Mendi | PNG_Highland | Oceania | Trans-New_Guinea | Erave | 143.95 | -6.6 | Bergstrom_et_al | PASS |
| papuan6278124 | Erave | Southern_Highlands | SOUTHERN_HIGHLANDS | Mendi | PNG_Highland | Oceania | Trans-New_Guinea | Erave | 143.95 | -6.6 | Bergstrom_et_al | PASS |
| papuan6278214 | Erave | Southern_Highlands | SOUTHERN_HIGHLANDS | Mendi | PNG_Highland | Oceania | Trans-New_Guinea | Erave | 143.95 | -6.6 | Bergstrom_et_al | PASS |
| papuan6278278 | Erave | Southern_Highlands | SOUTHERN_HIGHLANDS | Mendi | PNG_Highland | Oceania | Trans-New_Guinea | Erave | 143.95 | -6.6 | Bergstrom_et_al | PASS |
| papuan6278246 | Erave | Southern_Highlands | SOUTHERN_HIGHLANDS | Mendi | PNG_Highland | Oceania | Trans-New_Guinea | Erave | 143.95 | -6.6 | Bergstrom_et_al | PASS |
| papuan6278155 | Erave | Southern_Highlands | SOUTHERN_HIGHLANDS | Mendi | PNG_Highland | Oceania | Trans-New_Guinea | Erave | 143.95 | -6.6 | Bergstrom_et_al | PASS |
| papuan6278066 | Foi | Southern_Highlands | SOUTHERN_HIGHLANDS | Mendi | PNG_Highland | Oceania | Trans-New_Guinea | Foi | 143.5 | -6.5 | Bergstrom_et_al | PASS |
| papuan6278078 | Foi | Southern_Highlands | SOUTHERN_HIGHLANDS | Mendi | PNG_Highland | Oceania | Trans-New_Guinea | Foi | 143.5 | -6.5 | Bergstrom_et_al | PASS |
| papuan6278041 | Heneng | Southern_Highlands | SOUTHERN_HIGHLANDS | Mendi | PNG_Highland | Oceania | Trans-New_Guinea | Heneng | 143.5 | -6.1 | Bergstrom_et_al | PASS |
| papuan6278065 | Heneng | Southern_Highlands | SOUTHERN_HIGHLANDS | Mendi | PNG_Highland | Oceania | Trans-New_Guinea | Heneng | 143.5 | -6.1 | Bergstrom_et_al | PASS |
| papuan6278050 | Heneng | Southern_Highlands | SOUTHERN_HIGHLANDS | Mendi | PNG_Highland | Oceania | Trans-New_Guinea | Heneng | 143.5 | -6.1 | Bergstrom_et_al | PASS |
| papuan6278190 | Heneng | Southern_Highlands | SOUTHERN_HIGHLANDS | Mendi | PNG_Highland | Oceania | Trans-New_Guinea | Heneng | 143.5 | -6.1 | Bergstrom_et_al | PASS |
| papuan6278135 | Heneng | Southern_Highlands | SOUTHERN_HIGHLANDS | Mendi | PNG_Highland | Oceania | Trans-New_Guinea | Heneng | 143.5 | -6.1 | Bergstrom_et_al | PASS |
| papuan6278170 | Heneng | Southern_Highlands | SOUTHERN_HIGHLANDS | Mendi | PNG_Highland | Oceania | Trans-New_Guinea | Heneng | 143.5 | -6.1 | Bergstrom_et_al | PASS |
| papuan6278015 | Heneng | Southern_Highlands | SOUTHERN_HIGHLANDS | Mendi | PNG_Highland | Oceania | Trans-New_Guinea | Heneng | 143.5 | -6.1 | Bergstrom_et_al | PASS |
| papuan6278055 | Huli | Southern_Highlands | SOUTHERN_HIGHLANDS | Tauri | PNG_Highland | Oceania | Trans-New_Guinea | Huli | 143 | -5.95 | Bergstrom_et_al | PASS |
| papuan6278009 | Huli | Southern_Highlands | SOUTHERN_HIGHLANDS | Tauri | PNG_Highland | Oceania | Trans-New_Guinea | Huli | 143 | -5.95 | Bergstrom_et_al | PASS |
| papuan6278096 | Huli | Southern_Highlands | SOUTHERN_HIGHLANDS | Tauri | PNG_Highland | Oceania | Trans-New_Guinea | Huli | 143 | -5.95 | Bergstrom_et_al | PASS |
| papuan6278088 | Huli | Southern_Highlands | SOUTHERN_HIGHLANDS | Tauri | PNG_Highland | Oceania | Trans-New_Guinea | Huli | 143 | -5.95 | Bergstrom_et_al | PASS |
| papuan6278080 | Huli | Southern_Highlands | SOUTHERN_HIGHLANDS | Tauri | PNG_Highland | Oceania | Trans-New_Guinea | Huli | 143 | -5.95 | Bergstrom_et_al | PASS |
| papuan6278089 | Huli | Southern_Highlands | SOUTHERN_HIGHLANDS | Tauri | PNG_Highland | Oceania | Trans-New_Guinea | Huli | 143 | -5.95 | Bergstrom_et_al | PASS |
| papuan6278127 | Huli | Southern_Highlands | SOUTHERN_HIGHLANDS | Tauri | PNG_Highland | Oceania | Trans-New_Guinea | Huli | 143 | -5.95 | Bergstrom_et_al | PASS |
| papuan6278151 | Huli | Southern_Highlands | SOUTHERN_HIGHLANDS | Tauri | PNG_Highland | Oceania | Trans-New_Guinea | Huli | 143 | -5.95 | Bergstrom_et_al | PASS |
| papuan6278112 | Huli | Southern_Highlands | SOUTHERN_HIGHLANDS | Tauri | PNG_Highland | Oceania | Trans-New_Guinea | Huli | 143 | -5.95 | Bergstrom_et_al | PASS |
| papuan6278175 | Huli | Southern_Highlands | SOUTHERN_HIGHLANDS | Tauri | PNG_Highland | Oceania | Trans-New_Guinea | Huli | 143 | -5.95 | Bergstrom_et_al | PASS |
| papuan6278103 | Huli | Southern_Highlands | SOUTHERN_HIGHLANDS | Tauri | PNG_Highland | Oceania | Trans-New_Guinea | Huli | 143 | -5.95 | Bergstrom_et_al | PASS |
| papuan6278119 | Huli | Southern_Highlands | SOUTHERN_HIGHLANDS | Tauri | PNG_Highland | Oceania | Trans-New_Guinea | Huli | 143 | -5.95 | Bergstrom_et_al | PASS |
| papuan6278072 | Huli | Southern_Highlands | SOUTHERN_HIGHLANDS | Tauri | PNG_Highland | Oceania | Trans-New_Guinea | Huli | 143 | -5.95 | Bergstrom_et_al | PASS |
| papuan6278120 | Huli | Southern_Highlands | SOUTHERN_HIGHLANDS | Tauri | PNG_Highland | Oceania | Trans-New_Guinea | Huli | 143 | -5.95 | Bergstrom_et_al | PASS |
| papuan6278183 | Huli | Southern_Highlands | SOUTHERN_HIGHLANDS | Tauri | PNG_Highland | Oceania | Trans-New_Guinea | Huli | 143 | -5.95 | Bergstrom_et_al | PASS |
| papuan6278168 | Huli | Southern_Highlands | SOUTHERN_HIGHLANDS | Tauri | PNG_Highland | Oceania | Trans-New_Guinea | Huli | 143 | -5.95 | Bergstrom_et_al | PASS |
| papuan6278105 | Huli | Southern_Highlands | SOUTHERN_HIGHLANDS | Tauri | PNG_Highland | Oceania | Trans-New_Guinea | Huli | 143 | -5.95 | Bergstrom_et_al | PASS |
| papuan6278113 | Huli | Southern_Highlands | SOUTHERN_HIGHLANDS | Tauri | PNG_Highland | Oceania | Trans-New_Guinea | Huli | 143 | -5.95 | Bergstrom_et_al | PASS |

| | | | | | | | | | | | | |
|---|---|---|---|---|---|---|---|---|---|---|---|---|
| papuan6278121 | Huli | Southern_Highlands | SOUTHERN_HIGHLANDS | Tauri | PNG_Highland | Oceania | Trans-New_Guinea | Huli | 143 | -5.95 | Bergstrom_et_al | PASS |
| papuan6278232 | Huli | Southern_Highlands | SOUTHERN_HIGHLANDS | Tauri | PNG_Highland | Oceania | Trans-New_Guinea | Huli | 143 | -5.95 | Bergstrom_et_al | PASS |
| papuan6278090 | Imbongu | Southern_Highlands | SOUTHERN_HIGHLANDS | Ialibu | PNG_Highland | Oceania | Trans-New_Guinea | Imbongu | 144 | -6.15 | Bergstrom_et_al | PASS |
| papuan6278082 | Imbongu | Southern_Highlands | SOUTHERN_HIGHLANDS | Ialibu | PNG_Highland | Oceania | Trans-New_Guinea | Imbongu | 144 | -6.15 | Bergstrom_et_al | PASS |
| papuan6278194 | Imbongu | Southern_Highlands | SOUTHERN_HIGHLANDS | Ialibu | PNG_Highland | Oceania | Trans-New_Guinea | Imbongu | 144 | -6.15 | Bergstrom_et_al | PASS |
| papuan6278008 | Imbongu | Southern_Highlands | SOUTHERN_HIGHLANDS | Ialibu | PNG_Highland | Oceania | Trans-New_Guinea | Imbongu | 144 | -6.15 | Bergstrom_et_al | PASS |
| papuan6278250 | Samberigi | Southern_Highlands | SOUTHERN_HIGHLANDS | Lake_Kutubu | PNG_Highland | Oceania | Trans-New_Guinea | Samberigi | 144 | -6.7 | Bergstrom_et_al | PASS |
| papuan6278017 | West_Kewa | Southern_Highlands | SOUTHERN_HIGHLANDS | Mendi | PNG_Highland | Oceania | Trans-New_Guinea | West_Kewa | 143.75 | -6.3 | Bergstrom_et_al | PASS |
| papuan6278153 | Wiru | Southern_Highlands | SOUTHERN_HIGHLANDS | Ialibu | PNG_Highland | Oceania | Trans-New_Guinea | Wiru | 144.2 | -6.4 | Bergstrom_et_al | PASS |
| papuan6278177 | Wiru | Southern_Highlands | SOUTHERN_HIGHLANDS | Ialibu | PNG_Highland | Oceania | Trans-New_Guinea | Wiru | 144.2 | -6.4 | Bergstrom_et_al | PASS |
| papuan6278193 | Wiru | Southern_Highlands | SOUTHERN_HIGHLANDS | Ialibu | PNG_Highland | Oceania | Trans-New_Guinea | Wiru | 144.2 | -6.4 | Bergstrom_et_al | PASS |
| papuan6278106 | Wiru | Southern_Highlands | SOUTHERN_HIGHLANDS | Ialibu | PNG_Highland | Oceania | Trans-New_Guinea | Wiru | 144.2 | -6.4 | Bergstrom_et_al | PASS |
| papuan6278114 | Wiru | Southern_Highlands | SOUTHERN_HIGHLANDS | Ialibu | PNG_Highland | Oceania | Trans-New_Guinea | Wiru | 144.2 | -6.4 | Bergstrom_et_al | PASS |
| papuan6278107 | Wiru | Southern_Highlands | SOUTHERN_HIGHLANDS | Ialibu | PNG_Highland | Oceania | Trans-New_Guinea | Wiru | 144.2 | -6.4 | Bergstrom_et_al | PASS |
| papuan6278178 | Wiru | Southern_Highlands | SOUTHERN_HIGHLANDS | Ialibu | PNG_Highland | Oceania | Trans-New_Guinea | Wiru | 144.2 | -6.4 | Bergstrom_et_al | Kin |
| papuan6278186 | Wiru | Southern_Highlands | SOUTHERN_HIGHLANDS | Ialibu | PNG_Highland | Oceania | Trans-New_Guinea | Wiru | 144.2 | -6.4 | Bergstrom_et_al | PASS |
| papuan6278162 | Wiru | Southern_Highlands | SOUTHERN_HIGHLANDS | Ialibu | PNG_Highland | Oceania | Trans-New_Guinea | Wiru | 144.2 | -6.4 | Bergstrom_et_al | PASS |
| papuan6278145 | Wiru | Southern_Highlands | SOUTHERN_HIGHLANDS | Ialibu | PNG_Highland | Oceania | Trans-New_Guinea | Wiru | 144.2 | -6.4 | Bergstrom_et_al | PASS |
| papuan6278247 | Wiru | Southern_Highlands | SOUTHERN_HIGHLANDS | Ialibu | PNG_Highland | Oceania | Trans-New_Guinea | Wiru | 144.2 | -6.4 | Bergstrom_et_al | PASS |
| papuan6278271 | Wiru | Southern_Highlands | SOUTHERN_HIGHLANDS | Ialibu | PNG_Highland | Oceania | Trans-New_Guinea | Wiru | 144.2 | -6.4 | Bergstrom_et_al | PASS |
| papuan6277815 | Kyaka | Western_Highlands | WESTERN_HIGHLANDS | Wabag_Koles | PNG_Highland | Oceania | Trans-New_Guinea | Kyaka | 144.1 | -5.5 | Bergstrom_et_al | PASS |
| papuan6277875 | Maring | Western_Highlands | WESTERN_HIGHLANDS | Hagen | PNG_Highland | Oceania | Trans-New_Guinea | Maring | 144.6 | -5.5 | Bergstrom_et_al | PASS |
| papuan6278043 | Mbo-Ung | Western_Highlands | WESTERN_HIGHLANDS | Hagen | PNG_Highland | Oceania | Trans-New_Guinea | Mbo-Ung | 144.1 | -5.9 | Bergstrom_et_al | PASS |
| papuan6277863 | Mbo-Ung | Western_Highlands | WESTERN_HIGHLANDS | Hagen | PNG_Highland | Oceania | Trans-New_Guinea | Mbo-Ung | 144.1 | -5.9 | Bergstrom_et_al | PASS |
| papuan6277862 | Melpa | Western_Highlands | WESTERN_HIGHLANDS | Hagen | PNG_Highland | Oceania | Trans-New_Guinea | Melpa | 144.3 | -5.65 | Bergstrom_et_al | PASS |
| papuan6277903 | Melpa | Western_Highlands | WESTERN_HIGHLANDS | Hagen | PNG_Highland | Oceania | Trans-New_Guinea | Melpa | 144.3 | -5.65 | Bergstrom_et_al | PASS |
| papuan6277839 | Melpa | Western_Highlands | WESTERN_HIGHLANDS | Hagen | PNG_Highland | Oceania | Trans-New_Guinea | Melpa | 144.3 | -5.65 | Bergstrom_et_al | PASS |
| papuan6277887 | MIXED | Western_Highlands | WESTERN_HIGHLANDS | MIXED | PNG_Highland | Oceania | Trans-New_Guinea | MIXED | NA | NA | Bergstrom_et_al | PASS |
| papuan6277895 | MIXED | Western_Highlands | WESTERN_HIGHLANDS | MIXED | PNG_Highland | Oceania | Trans-New_Guinea | MIXED | NA | NA | Bergstrom_et_al | PASS |
| papuan6278225 | Nii | Western_Highlands | WESTERN_HIGHLANDS | Hagen | PNG_Highland | Oceania | Trans-New_Guinea | Nii | 144.5 | -5.8 | Bergstrom_et_al | PASS |
| papuan6277942 | Nii | Western_Highlands | WESTERN_HIGHLANDS | Hagen | PNG_Highland | Oceania | Trans-New_Guinea | Nii | 144.5 | -5.8 | Bergstrom_et_al | PASS |
| papuan6278098 | Nii | Western_Highlands | WESTERN_HIGHLANDS | Hagen | PNG_Highland | Oceania | Trans-New_Guinea | Nii | 144.5 | -5.8 | Bergstrom_et_al | PASS |
| papuan6277831 | Nii | Western_Highlands | WESTERN_HIGHLANDS | Hagen | PNG_Highland | Oceania | Trans-New_Guinea | Nii | 144.5 | -5.8 | Bergstrom_et_al | PASS |

| | | | | | | | | | | | | |
|---|---|---|---|---|---|---|---|---|---|---|---|---|
| papuan6277855 | MIXED | Western_Highlands | WESTERN_HIGHLANDS | MIXED | PNG_Highland | Oceania | Trans-New_Guinea | MIXED | NA | NA | Bergstrom_et_al | PASS |
| papuan6278233 | MIXED | Western_Highlands | WESTERN_HIGHLANDS | MIXED | PNG_Highland | Oceania | Trans-New_Guinea | MIXED | NA | NA | Bergstrom_et_al | PASS |
| papuan6277838 | Umbu-Ungu | Western_Highlands | WESTERN_HIGHLANDS | Tambul-Kaugel | PNG_Highland | Oceania | Trans-New_Guinea | Umbu-Ungu | 143.9 | -5.9 | Bergstrom_et_al | PASS |
| papuan6277894 | Umbu-Ungu | Western_Highlands | WESTERN_HIGHLANDS | Tambul-Kaugel | PNG_Highland | Oceania | Trans-New_Guinea | Umbu-Ungu | 143.9 | -5.9 | Bergstrom_et_al | PASS |
| papuan6277878 | Umbu-Ungu | Western_Highlands | WESTERN_HIGHLANDS | Tambul-Kaugel | PNG_Highland | Oceania | Trans-New_Guinea | Umbu-Ungu | 143.9 | -5.9 | Bergstrom_et_al | PASS |
| papuan6278013 | Umbu-Ungu | Western_Highlands | WESTERN_HIGHLANDS | Tambul-Kaugel | PNG_Highland | Oceania | Trans-New_Guinea | Umbu-Ungu | 143.9 | -5.9 | Bergstrom_et_al | PASS |
| papuan6277912 | Wahgi | Western_Highlands | WESTERN_HIGHLANDS | Minj | PNG_Highland | Oceania | Trans-New_Guinea | Wahgi | 144.7 | -5.9 | Bergstrom_et_al | PASS |
| papuan6277961 | Wahgi | Western_Highlands | WESTERN_HIGHLANDS | Minj | PNG_Highland | Oceania | Trans-New_Guinea | Wahgi | 144.7 | -5.9 | Bergstrom_et_al | PASS |
| papuan6277914 | Wahgi | Western_Highlands | WESTERN_HIGHLANDS | Minj | PNG_Highland | Oceania | Trans-New_Guinea | Wahgi | 144.7 | -5.9 | Bergstrom_et_al | PASS |
| papuan6277932 | Wahgi | Western_Highlands | WESTERN_HIGHLANDS | Minj | PNG_Highland | Oceania | Trans-New_Guinea | Wahgi | 144.7 | -5.9 | Bergstrom_et_al | PASS |
| papuan6278608 | Wahgi | Western_Highlands | WESTERN_HIGHLANDS | Minj | PNG_Highland | Oceania | Trans-New_Guinea | Wahgi | 144.7 | -5.9 | Bergstrom_et_al | PASS |
| papuan6277830 | Wahgi | Western_Highlands | WESTERN_HIGHLANDS | Minj | PNG_Highland | Oceania | Trans-New_Guinea | Wahgi | 144.7 | -5.9 | Bergstrom_et_al | PASS |
| papuan6277913 | Wahgi | Western_Highlands | WESTERN_HIGHLANDS | Minj | PNG_Highland | Oceania | Trans-New_Guinea | Wahgi | 144.7 | -5.9 | Bergstrom_et_al | PASS |
| papuan6278323 | MIXED | MIXED | WESTERN | MIXED | PNG_Lowland | Oceania | MIXED | MIXED | NA | NA | Bergstrom_et_al | Mixed |
| papuan6278282 | Southern_Kiwai | Western | WESTERN | Daru | PNG_Lowland | Oceania | Trans-New_Guinea | Southern_Kiwai | 143.3 | -8.7 | Bergstrom_et_al | PASS |
| papuan6278305 | Southern_Kiwai | Western | WESTERN | Daru | PNG_Lowland | Oceania | Trans-New_Guinea | Southern_Kiwai | 143.3 | -8.7 | Bergstrom_et_al | PASS |
| papuan6278326 | Southern_Kiwai | Western | WESTERN | Daru | PNG_Lowland | Oceania | Trans-New_Guinea | Southern_Kiwai | 143.3 | -8.7 | Bergstrom_et_al | PASS |
| papuan6278361 | Southern_Kiwai | Western | WESTERN | Daru | PNG_Lowland | Oceania | Trans-New_Guinea | Southern_Kiwai | 143.3 | -8.7 | Bergstrom_et_al | PASS |
| papuan6278345 | Southern_Kiwai | Western | WESTERN | Daru | PNG_Lowland | Oceania | Trans-New_Guinea | Southern_Kiwai | 143.3 | -8.7 | Bergstrom_et_al | PASS |
| LP6005592-DNA_C03 | Mbuti | Africa | NA | NA | Non_Oceania | Africa | Central_Sudanic | Efe | 29 | 1 | Mallick_et_al | PASS |
| LP6005441-DNA_B08 | Mbuti | Africa | NA | NA | Non_Oceania | Africa | Central_Sudanic | Efe | 29 | 1 | Mallick_et_al | PASS |
| LP6005441-DNA_A08 | Mbuti | Africa | NA | NA | Non_Oceania | Africa | Central_Sudanic | Efe | 29 | 1 | Mallick_et_al | PASS |
| SS6004471 | Mbuti | Africa | NA | NA | Non_Oceania | Africa | Central_Sudanic | Efe | 29 | 1 | Mallick_et_al | PASS |
| LP6005442-DNA_B02 | Yoruba | Africa | NA | NA | Non_Oceania | Africa | Niger-Congo | Yoruba | 3.9 | 7.4 | Mallick_et_al | PASS |
| LP6005442-DNA_A02 | Yoruba | Africa | NA | NA | Non_Oceania | Africa | Niger-Congo | Yoruba | 3.9 | 7.4 | Mallick_et_al | PASS |
| SS6004475 | Yoruba | Africa | NA | NA | Non_Oceania | Africa | Niger-Congo | Yoruba | 3.9 | 7.4 | Mallick_et_al | PASS |
| LP6005441-DNA_A05 | French | WestEurasia | NA | NA | Non_Oceania | WestEurasia | Indo-European | French | 2 | 46 | Mallick_et_al | PASS |
| LP6005441-DNA_B05 | French | WestEurasia | NA | NA | Non_Oceania | WestEurasia | Indo-European | French | 2 | 46 | Mallick_et_al | PASS |
| SS6004468 | French | WestEurasia | NA | NA | Non_Oceania | WestEurasia | Indo-European | French | 2 | 46 | Mallick_et_al | PASS |
| LP6005442-DNA_A11 | Spanish | WestEurasia | NA | NA | Non_Oceania | WestEurasia | Indo-European | Spanish | -4 | 39.9 | Mallick_et_al | PASS |
| LP6005442-DNA_B11 | Spanish | WestEurasia | NA | NA | Non_Oceania | WestEurasia | Indo-European | Spanish | -4 | 39.9 | Mallick_et_al | PASS |
| LP6005443-DNA_H01 | Tu | EastAsia | NA | NA | Non_Oceania | EastAsia | Mongolian | Tu | 101 | 36 | Mallick_et_al | PASS |
| LP6005441-DNA_D12 | Tu | EastAsia | NA | NA | Non_Oceania | EastAsia | Mongolian | Tu | 101 | 36 | Mallick_et_al | PASS |
| LP6005441-DNA_D05 | Han | EastAsia | NA | NA | Non_Oceania | EastAsia | Sino-Tibetan | Mandarin | 114 | 32.3 | Mallick_et_al | PASS |
| LP6005441-DNA_C05 | Han | EastAsia | NA | NA | Non_Oceania | EastAsia | Sino-Tibetan | Mandarin | 114 | 32.3 | Mallick_et_al | PASS |
| LP6005443-DNA_A02 | Tujia | EastAsia | NA | NA | Non_Oceania | EastAsia | Sino-Tibetan | Tujia | 109 | 29 | Mallick_et_al | PASS |

| | | | | | | | | | | | | |
|---|---|---|---|---|---|---|---|---|---|---|---|---|
| LP6005441-DNA_F12 | Tujia | EastAsia | NA | NA | Non_Oceania | EastAsia | Sino-Tibetan | Tujia | 109 | 29 | Mallick_et_al | PASS |
| LP6005443-DNA_E09 | Naxi | EastAsia | NA | NA | Non_Oceania | EastAsia | Sino-Tibetan | Naxi | 100 | 26 | Mallick_et_al | PASS |
| LP6005441-DNA_B09 | Naxi | EastAsia | NA | NA | Non_Oceania | EastAsia | Sino-Tibetan | Naxi | 100 | 26 | Mallick_et_al | PASS |
| LP6005442-DNA_H01 | Yi | EastAsia | NA | NA | Non_Oceania | EastAsia | Sino-Tibetan | Yi | 103 | 28 | Mallick_et_al | PASS |
| LP6005441-DNA_B07 | Lahu | EastAsia | NA | NA | Non_Oceania | EastAsia | Sino-Tibetan | Lahu | 100 | 22 | Mallick_et_al | PASS |
| LP6005443-DNA_E01 | Lahu | EastAsia | NA | NA | Non_Oceania | EastAsia | Sino-Tibetan | Lahu | 100 | 22 | Mallick_et_al | PASS |
| LP6005441-DNA_C08 | Miao | EastAsia | NA | NA | Non_Oceania | EastAsia | Hmong-Mien | Miao | 109 | 28 | Mallick_et_al | PASS |
| LP6005441-DNA_D08 | Miao | EastAsia | NA | NA | Non_Oceania | EastAsia | Hmong-Mien | Miao | 109 | 28 | Mallick_et_al | PASS |
| LP6005443-DNA_G01 | She | EastAsia | NA | NA | Non_Oceania | EastAsia | Hmong-Mien | She | 119 | 27 | Mallick_et_al | PASS |
| LP6005443-DNA_F01 | She | EastAsia | NA | NA | Non_Oceania | EastAsia | Hmong-Mien | She | 119 | 27 | Mallick_et_al | PASS |
| LP6005592-DNA_D03 | Dai | EastAsia | NA | NA | Non_Oceania | EastAsia | Tai-Kadai | Central_Tai | 100 | 21 | Mallick_et_al | PASS |
| LP6005443-DNA_B01 | Dai | EastAsia | NA | NA | Non_Oceania | EastAsia | Tai-Kadai | Central_Tai | 100 | 21 | Mallick_et_al | PASS |
| LP6005441-DNA_D04 | Dai | EastAsia | NA | NA | Non_Oceania | EastAsia | Tai-Kadai | Central_Tai | 100 | 21 | Mallick_et_al | PASS |
| SS6004467 | Dai | EastAsia | NA | NA | Non_Oceania | EastAsia | Tai-Kadai | Central_Tai | 100 | 21 | Mallick_et_al | PASS |
| LP6005443-DNA_A07 | Thai | EastAsia | NA | NA | Non_Oceania | EastAsia | Tai-Kadai | Central_Tai | 100.5 | 13.8 | Mallick_et_al | PASS |
| LP6005443-DNA_B07 | Thai | EastAsia | NA | NA | Non_Oceania | EastAsia | Tai-Kadai | Central_Tai | 100.5 | 13.8 | Mallick_et_al | PASS |
| LP6005442-DNA_D11 | Kinh | EastAsia | NA | NA | Non_Oceania | EastAsia | Austro-Asiatic | Vietnamese | 105.9 | 21 | Mallick_et_al | PASS |
| LP6005442-DNA_C11 | Kinh | EastAsia | NA | NA | Non_Oceania | EastAsia | Austro-Asiatic | Vietnamese | 105.9 | 21 | Mallick_et_al | PASS |
| LP6005519-DNA_B06 | Burmese | EastAsia | NA | NA | Non_Oceania | EastAsia | Sino-Tibetan | Burmese | 96.7 | 17 | Mallick_et_al | PASS |
| LP6005519-DNA_A06 | Burmese | EastAsia | NA | NA | Non_Oceania | EastAsia | Sino-Tibetan | Burmese | 96.7 | 17 | Mallick_et_al | PASS |
| LP6005442-DNA_C07 | Ami | EastAsia_AN | NA | NA | Non_Oceania | EastAsia | Austronesian | Amis | 121.19 | 22.84 | Mallick_et_al | PASS |
| LP6005443-DNA_G05 | Ami | EastAsia_AN | NA | NA | Non_Oceania | EastAsia | Austronesian | Amis | 121.19 | 22.84 | Mallick_et_al | PASS |
| LP6005442-DNA_E07 | Atayal | EastAsia_AN | NA | NA | Non_Oceania | EastAsia | Austronesian | Atayal | 121.3 | 24.61 | Mallick_et_al | PASS |
| LP6005519-DNA_C06 | Igorot | EastAsia_AN | NA | NA | Non_Oceania | EastAsia | Austronesian | Northern_Luzon | 121 | 17.1 | Mallick_et_al | PASS |
| LP6005519-DNA_D06 | Igorot | EastAsia_AN | NA | NA | Non_Oceania | EastAsia | Austronesian | Northern_Luzon | 121 | 17.1 | Mallick_et_al | PASS |
| LP6005519-DNA_F06 | Dusun | EastAsia_AN | NA | NA | Non_Oceania | EastAsia | Austronesian | Dusun | 114.7 | 4.7 | Mallick_et_al | PASS |
| LP6005519-DNA_E06 | Dusun | EastAsia_AN | NA | NA | Non_Oceania | EastAsia | Austronesian | Dusun | 114.7 | 4.7 | Mallick_et_al | PASS |
| SS6004478 | Australian | Australia | NA | NA | Australia | Oceania | Pama-Nyungan | Paman | 143 | -13 | Mallick_et_al | PASS |
| SS6004477 | Australian | Australia | NA | NA | Australia | Oceania | Pama-Nyungan | Paman | 143 | -13 | Mallick_et_al | PASS |
| LP6005441-DNA_B03 | Bougainville | Bougainville | BOUGAINVILLE | Bougainville | Solomon_Arch | Oceania | Austronesian | Halia | 155 | -6 | Mallick_et_al | PASS |
| LP6005441-DNA_A03 | Bougainville | Bougainville | BOUGAINVILLE | Bougainville | Solomon_Arch | Oceania | Austronesian | Halia | 155 | -6 | Mallick_et_al | PASS |
| LP6005441-DNA_B10 | Papuan | S_Papuan | EAST_SEPIK | Maprik | PNG_Highland | Oceania | Sepik | Ambulas | 143 | -4 | Mallick_et_al | PASS |
| LP6005443-DNA_F07 | Papuan | S_Papuan | EAST_SEPIK | Maprik | PNG_Highland | Oceania | Sepik | Ambulas | 143 | -4 | Mallick_et_al | PASS |
| LP6005443-DNA_A08 | Papuan | S_Papuan | EAST_SEPIK | Maprik | PNG_Highland | Oceania | Sepik | Ambulas | 143 | -4 | Mallick_et_al | PASS |

| | | | | | | | | | | | | |
|---|---|---|---|---|---|---|---|---|---|---|---|---|
| LP6005443-DNA_B08 | Papuan | S_Papuan | EAST_SEPIK | Maprik | PNG_Highland | Oceania | Sepik | Ambulas | 143 | -4 | Mallick_et_al | PASS |
| LP6005443-DNA_C07 | Papuan | S_Papuan | EAST_SEPIK | Maprik | PNG_Highland | Oceania | Sepik | Ambulas | 143 | -4 | Mallick_et_al | PASS |
| LP6005443-DNA_G07 | Papuan | S_Papuan | EAST_SEPIK | Maprik | PNG_Highland | Oceania | Sepik | Ambulas | 143 | -4 | Mallick_et_al | PASS |
| LP6005443-DNA_D08 | Papuan | S_Papuan | EAST_SEPIK | Maprik | PNG_Highland | Oceania | Sepik | Ambulas | 143 | -4 | Mallick_et_al | PASS |
| LP6005443-DNA_E08 | Papuan | S_Papuan | EAST_SEPIK | Maprik | PNG_Highland | Oceania | Sepik | Ambulas | 143 | -4 | Mallick_et_al | PASS |
| LP6005443-DNA_H07 | Papuan | S_Papuan | EAST_SEPIK | Maprik | PNG_Lowland | Oceania | Sepik | Ambulas | 143 | -4 | Mallick_et_al | PASS |
| LP6005441-DNA_A10 | Papuan | S_Papuan | EAST_SEPIK | Maprik | PNG_Lowland | Oceania | Sepik | Ambulas | 143 | -4 | Mallick_et_al | PASS |
| LP6005443-DNA_E07 | Papuan | S_Papuan | EAST_SEPIK | Maprik | PNG_Lowland | Oceania | Sepik | Ambulas | 143 | -4 | Mallick_et_al | PASS |
| LP6005443-DNA_D07 | Papuan | S_Papuan | EAST_SEPIK | Maprik | PNG_Lowland | Oceania | Sepik | Ambulas | 143 | -4 | Mallick_et_al | PASS |
| LP6005443-DNA_C08 | Papuan | S_Papuan | EAST_SEPIK | Maprik | PNG_Lowland | Oceania | Sepik | Ambulas | 143 | -4 | Mallick_et_al | PASS |
| SS6004472 | Papuan | S_Papuan | EAST_SEPIK | Maprik | PNG_Lowland | Oceania | Sepik | Ambulas | 143 | -4 | Mallick_et_al | PASS |
| UV500 | Lavongai | Manus_New_Ireland | NEW_IRELAND | North_Lavongai | Bismarck_Arch | Oceania | Austronesian | Lavongai | 150.27 | -2.53 | Vernot_et_al | PASS |
| UV518 | Mussau | Manus_New_Ireland | NEW_IRELAND | Kaupgu | Bismarck_Arch | Oceania | Austronesian | Mussau | 149.73 | -1.58 | Vernot_et_al | PASS |
| UV573 | Nailik | Manus_New_Ireland | NEW_IRELAND | Nailik | Bismarck_Arch | Oceania | Austronesian | Nalik | 151.3 | -2.94 | Vernot_et_al | PASS |
| UV580 | Nailik | Manus_New_Ireland | NEW_IRELAND | Nailik | Bismarck_Arch | Oceania | Austronesian | Nalik | 151.3 | -2.94 | Vernot_et_al | Kin |
| UV043 | Baining | East_New_Britain | EAST_NEW_BRITAIN | Mali | Bismarck_Arch | Oceania | East_New_Britain | Mali | 152 | -4.52 | Vernot_et_al | PASS |
| UV305 | Baining | East_New_Britain | EAST_NEW_BRITAIN | Kaket | Bismarck_Arch | Oceania | East_New_Britain | Qaqet | 152 | -4.52 | Vernot_et_al | PASS |
| UV1134 | Ata | West_New_Britain | WEST_NEW_BRITAIN | Luge | Bismarck_Arch | Oceania | Yele-West_New_Britain | Pele-Ata | 151.03 | -5.57 | Vernot_et_al | PASS |
| UV1230 | Ata | West_New_Britain | WEST_NEW_BRITAIN | Uasilau | Bismarck_Arch | Oceania | Yele-West_New_Britain | Pele-Ata | 151.03 | -5.57 | Vernot_et_al | PASS |
| UV1042 | Mamusi | West_New_Britain | WEST_NEW_BRITAIN | Kisiluvi | Bismarck_Arch | Oceania | Austronesian | Mamusi | 150.97 | -5.87 | Vernot_et_al | PASS |
| UV1224 | Mamusi | West_New_Britain | WEST_NEW_BRITAIN | Paleabu | Bismarck_Arch | Oceania | Austronesian | Mamusi | 150.97 | -5.87 | Vernot_et_al | PASS |
| UV1196 | Melamala | West_New_Britain | WEST_NEW_BRITAIN | Ubili | Bismarck_Arch | Oceania | Austronesian | Meramera | 151.33 | -5.02 | Vernot_et_al | PASS |
| UV1263 | Pasismanua | West_New_Britain | WEST_NEW_BRITAIN | Poronga | Bismarck_Arch | Oceania | Austronesian | Pasismanua | 150.09 | -6.27 | Vernot_et_al | PASS |
| UV1266 | Pasismanua | West_New_Britain | WEST_NEW_BRITAIN | Poronga | Bismarck_Arch | Oceania | Austronesian | Pasismanua | 150.09 | -6.27 | Vernot_et_al | Kin |
| UV886 | Nakanai_bileki | West_New_Britain | WEST_NEW_BRITAIN | Bileki | Bismarck_Arch | Oceania | Austronesian | Nakanai | 150.66 | -5.67 | Vernot_et_al | PASS |
| UV897 | Nakanai_bileki | West_New_Britain | WEST_NEW_BRITAIN | Bileki | Bismarck_Arch | Oceania | Austronesian | Nakanai | 150.66 | -5.67 | Vernot_et_al | PASS |
| UV910 | Nakanai_bileki | West_New_Britain | WEST_NEW_BRITAIN | Bileki | Bismarck_Arch | Oceania | Austronesian | Nakanai | 150.66 | -5.67 | Vernot_et_al | PASS |
| UV919 | Nakanai_bileki | West_New_Britain | WEST_NEW_BRITAIN | Bileki | Bismarck_Arch | Oceania | Austronesian | Nakanai | 150.66 | -5.67 | Vernot_et_al | PASS |
| UV923 | Nakanai_bileki | West_New_Britain | WEST_NEW_BRITAIN | Bileki | Bismarck_Arch | Oceania | Austronesian | Nakanai | 150.66 | -5.67 | Vernot_et_al | PASS |
| UV925 | Nakanai_bileki | West_New_Britain | WEST_NEW_BRITAIN | Bileki | Bismarck_Arch | Oceania | Austronesian | Nakanai | 150.66 | -5.67 | Vernot_et_al | PASS |
| UV927 | Nakanai_bileki | West_New_Britain | WEST_NEW_BRITAIN | Bileki | Bismarck_Arch | Oceania | Austronesian | Nakanai | 150.66 | -5.67 | Vernot_et_al | Kin |
| UV929 | Nakanai_bileki | West_New_Britain | WEST_NEW_BRITAIN | Bileki | Bismarck_Arch | Oceania | Austronesian | Nakanai | 150.66 | -5.67 | Vernot_et_al | PASS |
| UV931 | Nakanai_bileki | West_New_Britain | WEST_NEW_BRITAIN | Bileki | Bismarck_Arch | Oceania | Austronesian | Nakanai | 150.66 | -5.67 | Vernot_et_al | PASS |

| | | | | | | | | | | | | |
|---|---|---|---|---|---|---|---|---|---|---|---|---|
| UV940 | Nakanai_bileki | West_New_Britain | WEST_NEW_BRITAIN | Bileki | Bismarck_Arch | Oceania | Austronesian | Nakanai | 150.66 | -5.67 | Vernot_et_al | PASS |
| UV944 | Nakanai_bileki | West_New_Britain | WEST_NEW_BRITAIN | Bileki | Bismarck_Arch | Oceania | Austronesian | Nakanai | 150.66 | -5.67 | Vernot_et_al | PASS |
| UV946 | Nakanai | West_New_Britain | WEST_NEW_BRITAIN | Bileki | Bismarck_Arch | Oceania | Austronesian | Nakanai | 150.66 | -5.67 | Vernot_et_al | PASS |
| UV952 | Nakanai | West_New_Britain | WEST_NEW_BRITAIN | Bileki | Bismarck_Arch | Oceania | Austronesian | Nakanai | 150.66 | -5.67 | Vernot_et_al | Kin |
| UV956 | Nakanai | West_New_Britain | WEST_NEW_BRITAIN | Bileki | Bismarck_Arch | Oceania | Austronesian | Nakanai | 150.66 | -5.67 | Vernot_et_al | PASS |
| UV958 | Nakanai | West_New_Britain | WEST_NEW_BRITAIN | Bileki | Bismarck_Arch | Oceania | Austronesian | Nakanai | 150.66 | -5.67 | Vernot_et_al | PASS |
| UV964 | Nakanai | West_New_Britain | WEST_NEW_BRITAIN | Bileki | Bismarck_Arch | Oceania | Austronesian | Nakanai | 150.66 | -5.67 | Vernot_et_al | PASS |
| UV971 | Nakanai | West_New_Britain | WEST_NEW_BRITAIN | Bileki | Bismarck_Arch | Oceania | Austronesian | Nakanai | 150.66 | -5.67 | Vernot_et_al | PASS |
| UV979 | Nakanai | West_New_Britain | WEST_NEW_BRITAIN | Bileki | Bismarck_Arch | Oceania | Austronesian | Nakanai | 150.66 | -5.67 | Vernot_et_al | Kin |
| UV986 | Nakanai | West_New_Britain | WEST_NEW_BRITAIN | Bileki | Bismarck_Arch | Oceania | Austronesian | Nakanai | 150.66 | -5.67 | Vernot_et_al | PASS |
| B00FLGA | Atayal | EastAsia_AN | NA | NA | Non_Oceania | EastAsia | Austronesian | Atayal | 121.3 | 24.61 | Choin_et_al | PASS |
| B00FLGM | Atayal | EastAsia_AN | NA | NA | Non_Oceania | EastAsia | Austronesian | Atayal | 121.3 | 24.61 | Choin_et_al | PASS |
| B00FLGQ | Atayal | EastAsia_AN | NA | NA | Non_Oceania | EastAsia | Austronesian | Atayal | 121.3 | 24.61 | Choin_et_al | PASS |
| B00FLGU | Atayal | EastAsia_AN | NA | NA | Non_Oceania | EastAsia | Austronesian | Atayal | 121.3 | 24.61 | Choin_et_al | PASS |
| B00FLH2 | Atayal | EastAsia_AN | NA | NA | Non_Oceania | EastAsia | Austronesian | Atayal | 121.3 | 24.61 | Choin_et_al | PASS |
| B00FLH6 | Atayal | EastAsia_AN | NA | NA | Non_Oceania | EastAsia | Austronesian | Atayal | 121.3 | 24.61 | Choin_et_al | PASS |
| B00FLHA | Atayal | EastAsia_AN | NA | NA | Non_Oceania | EastAsia | Austronesian | Atayal | 121.3 | 24.61 | Choin_et_al | PASS |
| B00FLGE | Atayal | EastAsia_AN | NA | NA | Non_Oceania | EastAsia | Austronesian | Atayal | 121.3 | 24.61 | Choin_et_al | PASS |
| B00FLGI | Atayal | EastAsia_AN | NA | NA | Non_Oceania | EastAsia | Austronesian | Atayal | 121.3 | 24.61 | Choin_et_al | PASS |
| B00FLGY | Atayal | EastAsia_AN | NA | NA | Non_Oceania | EastAsia | Austronesian | Atayal | 121.3 | 24.61 | Choin_et_al | PASS |
| B00FLHE | Paiwan | EastAsia_AN | NA | NA | Non_Oceania | EastAsia | Austronesian | Paiwan | 120.69 | 22.68 | Choin_et_al | PASS |
| B00FLHM | Paiwan | EastAsia_AN | NA | NA | Non_Oceania | EastAsia | Austronesian | Paiwan | 120.69 | 22.68 | Choin_et_al | Kin |
| B00FLHQ | Paiwan | EastAsia_AN | NA | NA | Non_Oceania | EastAsia | Austronesian | Paiwan | 120.69 | 22.68 | Choin_et_al | PASS |
| B00FLHU | Paiwan | EastAsia_AN | NA | NA | Non_Oceania | EastAsia | Austronesian | Paiwan | 120.69 | 22.68 | Choin_et_al | PASS |
| B00FLI2 | Paiwan | EastAsia_AN | NA | NA | Non_Oceania | EastAsia | Austronesian | Paiwan | 120.69 | 22.68 | Choin_et_al | PASS |
| B00FLIA | Paiwan | EastAsia_AN | NA | NA | Non_Oceania | EastAsia | Austronesian | Paiwan | 120.69 | 22.68 | Choin_et_al | PASS |
| B00FLHI | Paiwan | EastAsia_AN | NA | NA | Non_Oceania | EastAsia | Austronesian | Paiwan | 120.69 | 22.68 | Choin_et_al | PASS |
| B00FLHY | Paiwan | EastAsia_AN | NA | NA | Non_Oceania | EastAsia | Austronesian | Paiwan | 120.69 | 22.68 | Choin_et_al | PASS |
| B00FLIE | Paiwan | EastAsia_AN | NA | NA | Non_Oceania | EastAsia | Austronesian | Paiwan | 120.69 | 22.68 | Choin_et_al | PASS |
| B00I2BM | Vella_Lavella | Vella_Lavella | NA | Vella_Lavella | Solomon_Arch | Oceania | Central_Solomons | Bilua | 156.37 | -7.76 | Choin_et_al | PASS |
| B00I2BN | Vella_Lavella | Vella_Lavella | NA | Vella_Lavella | Solomon_Arch | Oceania | Central_Solomons | Bilua | 156.37 | -7.76 | Choin_et_al | PASS |
| B00I2BP | Vella_Lavella | Vella_Lavella | NA | Vella_Lavella | Solomon_Arch | Oceania | Central_Solomons | Bilua | 156.37 | -7.76 | Choin_et_al | PASS |
| B00I2BQ | Vella_Lavella | Vella_Lavella | NA | Vella_Lavella | Solomon_Arch | Oceania | Central_Solomons | Bilua | 156.37 | -7.76 | Choin_et_al | PASS |
| B00I2BR | Vella_Lavella | Vella_Lavella | NA | Vella_Lavella | Solomon_Arch | Oceania | Central_Solomons | Bilua | 156.37 | -7.76 | Choin_et_al | PASS |
| B00I2BS | Vella_Lavella | Vella_Lavella | NA | Vella_Lavella | Solomon_Arch | Oceania | Central_Solomons | Bilua | 156.37 | -7.76 | Choin_et_al | PASS |
| B00I2BT | Vella_Lavella | Vella_Lavella | NA | Vella_Lavella | Solomon_Arch | Oceania | Central_Solomons | Bilua | 156.37 | -7.76 | Choin_et_al | PASS |
| B00I2BU | Vella_Lavella | Vella_Lavella | NA | Vella_Lavella | Solomon_Arch | Oceania | Central_Solomons | Bilua | 156.37 | -7.76 | Choin_et_al | PASS |
| B00I2BV | Vella_Lavella | Vella_Lavella | NA | Vella_Lavella | Solomon_Arch | Oceania | Central_Solomons | Bilua | 156.37 | -7.76 | Choin_et_al | PASS |
| B00I2BW | Vella_Lavella | Vella_Lavella | NA | Vella_Lavella | Solomon_Arch | Oceania | Central_Solomons | Bilua | 156.37 | -7.76 | Choin_et_al | PASS |
| B00I2BX | Vella_Lavella | Vella_Lavella | NA | Vella_Lavella | Solomon_Arch | Oceania | Central_Solomons | Bilua | 156.37 | -7.76 | Choin_et_al | PASS |
| B00I2BY | Vella_Lavella | Vella_Lavella | NA | Vella_Lavella | Solomon_Arch | Oceania | Central_Solomons | Bilua | 156.37 | -7.76 | Choin_et_al | PASS |
| B00I2BZ | Vella_Lavella | Vella_Lavella | NA | Vella_Lavella | Solomon_Arch | Oceania | Central_Solomons | Bilua | 156.37 | -7.76 | Choin_et_al | PASS |
| B00I2C0 | Vella_Lavella | Vella_Lavella | NA | Vella_Lavella | Solomon_Arch | Oceania | Central_Solomons | Bilua | 156.37 | -7.76 | Choin_et_al | PASS |
| B00I2C1 | Vella_Lavella | Vella_Lavella | NA | Vella_Lavella | Solomon_Arch | Oceania | Central_Solomons | Bilua | 156.37 | -7.76 | Choin_et_al | PASS |

| | | | | | | | | | | | | |
|---|---|---|---|---|---|---|---|---|---|---|---|---|
| B00I2C2 | Vella_Lavella | Vella_Lavella | NA | Vella_Lavella | Solomon_Arch | Oceania | Central_Solomons | Bilua | 156.37 | -7.76 | Choin_et_al | PASS |
| B00I2C3 | Vella_Lavella | Vella_Lavella | NA | Vella_Lavella | Solomon_Arch | Oceania | Central_Solomons | Bilua | 156.37 | -7.76 | Choin_et_al | PASS |
| B00I2C4 | Vella_Lavella | Vella_Lavella | NA | Vella_Lavella | Solomon_Arch | Oceania | Central_Solomons | Bilua | 156.37 | -7.76 | Choin_et_al | PASS |
| B00I2B2 | Malaita | Malaita | NA | Malaita | Solomon_Arch | Oceania | Austronesian | Malaitan | 159.95 | -9.02 | Choin_et_al | PASS |
| B00I2B8 | Malaita | Malaita | NA | Malaita | Solomon_Arch | Oceania | Austronesian | Malaitan | 159.95 | -9.02 | Choin_et_al | PASS |
| B00I2B3 | Malaita | Malaita | NA | Malaita | Solomon_Arch | Oceania | Austronesian | Malaitan | 159.95 | -9.02 | Choin_et_al | PASS |
| B00I2B4 | Malaita | Malaita | NA | Malaita | Solomon_Arch | Oceania | Austronesian | Malaitan | 159.95 | -9.02 | Choin_et_al | PASS |
| B00I2B5 | Malaita | Malaita | NA | Malaita | Solomon_Arch | Oceania | Austronesian | Malaitan | 159.95 | -9.02 | Choin_et_al | PASS |
| B00I2B6 | Malaita | Malaita | NA | Malaita | Solomon_Arch | Oceania | Austronesian | Malaitan | 159.95 | -9.02 | Choin_et_al | PASS |
| B00I2B7 | Malaita | Malaita | NA | Malaita | Solomon_Arch | Oceania | Austronesian | Malaitan | 159.95 | -9.02 | Choin_et_al | PASS |
| B00I2B9 | Malaita | Malaita | NA | Malaita | Solomon_Arch | Oceania | Austronesian | Malaitan | 159.95 | -9.02 | Choin_et_al | PASS |
| B00I2BA | Malaita | Malaita | NA | Malaita | Solomon_Arch | Oceania | Austronesian | Malaitan | 159.95 | -9.02 | Choin_et_al | PASS |
| B00I2BB | Malaita | Malaita | NA | Malaita | Solomon_Arch | Oceania | Austronesian | Malaitan | 159.95 | -9.02 | Choin_et_al | PASS |
| B00I2BC | Malaita | Malaita | NA | Malaita | Solomon_Arch | Oceania | Austronesian | Malaitan | 159.95 | -9.02 | Choin_et_al | PASS |
| B00I2BD | Malaita | Malaita | NA | Malaita | Solomon_Arch | Oceania | Austronesian | Malaitan | 159.95 | -9.02 | Choin_et_al | PASS |
| B00I2BE | Malaita | Malaita | NA | Malaita | Solomon_Arch | Oceania | Austronesian | Malaitan | 159.95 | -9.02 | Choin_et_al | PASS |
| B00I2BF | Malaita | Malaita | NA | Malaita | Solomon_Arch | Oceania | Austronesian | Malaitan | 159.95 | -9.02 | Choin_et_al | PASS |
| B00I2BG | Malaita | Malaita | NA | Malaita | Solomon_Arch | Oceania | Austronesian | Malaitan | 159.95 | -9.02 | Choin_et_al | PASS |
| B00I2BH | Malaita | Malaita | NA | Malaita | Solomon_Arch | Oceania | Austronesian | Malaitan | 159.95 | -9.02 | Choin_et_al | PASS |
| B00I2BJ | Malaita | Malaita | NA | Malaita | Solomon_Arch | Oceania | Austronesian | Malaitan | 159.95 | -9.02 | Choin_et_al | PASS |
| B00I2BL | Malaita | Malaita | NA | Malaita | Solomon_Arch | Oceania | Austronesian | Malaitan | 159.95 | -9.02 | Choin_et_al | PASS |
| B00I2AI | Santa_Cruz | Santa_Cruz | NA | Santa_Cruz | Solomon_Arch | Oceania | Austronesian | Reefs_Santa_Cruz | 165.85 | -10.76 | Choin_et_al | PASS |
| B00I2AJ | Santa_Cruz | Santa_Cruz | NA | Santa_Cruz | Solomon_Arch | Oceania | Austronesian | Reefs_Santa_Cruz | 165.85 | -10.76 | Choin_et_al | PASS |
| B00I2AL | Santa_Cruz | Santa_Cruz | NA | Santa_Cruz | Solomon_Arch | Oceania | Austronesian | Reefs_Santa_Cruz | 165.85 | -10.76 | Choin_et_al | PASS |
| B00I2AM | Santa_Cruz | Santa_Cruz | NA | Santa_Cruz | Solomon_Arch | Oceania | Austronesian | Reefs_Santa_Cruz | 165.85 | -10.76 | Choin_et_al | PASS |
| B00I2AN | Santa_Cruz | Santa_Cruz | NA | Santa_Cruz | Solomon_Arch | Oceania | Austronesian | Reefs_Santa_Cruz | 165.85 | -10.76 | Choin_et_al | PASS |
| B00I2AO | Santa_Cruz | Santa_Cruz | NA | Santa_Cruz | Solomon_Arch | Oceania | Austronesian | Reefs_Santa_Cruz | 165.85 | -10.76 | Choin_et_al | PASS |
| B00I2AP | Santa_Cruz | Santa_Cruz | NA | Santa_Cruz | Solomon_Arch | Oceania | Austronesian | Reefs_Santa_Cruz | 165.85 | -10.76 | Choin_et_al | PASS |
| B00I2AQ | Santa_Cruz | Santa_Cruz | NA | Santa_Cruz | Solomon_Arch | Oceania | Austronesian | Reefs_Santa_Cruz | 165.85 | -10.76 | Choin_et_al | PASS |
| B00I2AR | Santa_Cruz | Santa_Cruz | NA | Santa_Cruz | Solomon_Arch | Oceania | Austronesian | Reefs_Santa_Cruz | 165.85 | -10.76 | Choin_et_al | PASS |
| B00I2AS | Santa_Cruz | Santa_Cruz | NA | Santa_Cruz | Solomon_Arch | Oceania | Austronesian | Reefs_Santa_Cruz | 165.85 | -10.76 | Choin_et_al | PASS |
| B00I2AU | Santa_Cruz | Santa_Cruz | NA | Santa_Cruz | Solomon_Arch | Oceania | Austronesian | Reefs_Santa_Cruz | 165.85 | -10.76 | Choin_et_al | PASS |
| B00I2AV | Santa_Cruz | Santa_Cruz | NA | Santa_Cruz | Solomon_Arch | Oceania | Austronesian | Reefs_Santa_Cruz | 165.85 | -10.76 | Choin_et_al | PASS |
| B00I2AX | Santa_Cruz | Santa_Cruz | NA | Santa_Cruz | Solomon_Arch | Oceania | Austronesian | Reefs_Santa_Cruz | 165.85 | -10.76 | Choin_et_al | PASS |
| B00I2AY | Santa_Cruz | Santa_Cruz | NA | Santa_Cruz | Solomon_Arch | Oceania | Austronesian | Reefs_Santa_Cruz | 165.85 | -10.76 | Choin_et_al | PASS |
| B00FLID | Bellona | Bellona_Rennell | NA | Bellona | Solomon_Arch | Oceania | Austronesian | Rennell_Bellona | 159.79 | -11.3 | Choin_et_al | PASS |
| B00FLIH | Bellona | Bellona_Rennell | NA | Bellona | Solomon_Arch | Oceania | Austronesian | Rennell_Bellona | 159.79 | -11.3 | Choin_et_al | PASS |
| B00FLS1 | Bellona | Bellona_Rennell | NA | Bellona | Solomon_Arch | Oceania | Austronesian | Rennell_Bellona | 159.79 | -11.3 | Choin_et_al | PASS |
| B00FLS5 | Bellona | Bellona_Rennell | NA | Bellona | Solomon_Arch | Oceania | Austronesian | Rennell_Bellona | 159.79 | -11.3 | Choin_et_al | PASS |
| B00FLIL | Bellona | Bellona_Rennell | NA | Bellona | Solomon_Arch | Oceania | Austronesian | Rennell_Bellona | 159.79 | -11.3 | Choin_et_al | PASS |
| B00FLIP | Bellona | Bellona_Rennell | NA | Bellona | Solomon_Arch | Oceania | Austronesian | Rennell_Bellona | 159.79 | -11.3 | Choin_et_al | PASS |
| B00FLI1 | Bellona | Bellona_Rennell | NA | Bellona | Solomon_Arch | Oceania | Austronesian | Rennell_Bellona | 159.79 | -11.3 | Choin_et_al | PASS |
| B00FLI5 | Bellona | Bellona_Rennell | NA | Bellona | Solomon_Arch | Oceania | Austronesian | Rennell_Bellona | 159.79 | -11.3 | Choin_et_al | PASS |
| B00FLI9 | Bellona | Bellona_Rennell | NA | Bellona | Solomon_Arch | Oceania | Austronesian | Rennell_Bellona | 159.79 | -11.3 | Choin_et_al | PASS |
| B00FLIT | Bellona | Bellona_Rennell | NA | Bellona | Solomon_Arch | Oceania | Austronesian | Rennell_Bellona | 159.79 | -11.3 | Choin_et_al | PASS |
| B00FLS9 | Bellona | Bellona_Rennell | NA | Bellona | Solomon_Arch | Oceania | Austronesian | Rennell_Bellona | 159.79 | -11.3 | Choin_et_al | PASS |
| B00FLHX | Bellona | Bellona_Rennell | NA | Bellona | Solomon_Arch | Oceania | Austronesian | Rennell_Bellona | 159.79 | -11.3 | Choin_et_al | PASS |
| B00FLIX | Bellona | Bellona_Rennell | NA | Bellona | Solomon_Arch | Oceania | Austronesian | Rennell_Bellona | 159.79 | -11.3 | Choin_et_al | PASS |
| B00FLJ1 | Bellona | Bellona_Rennell | NA | Bellona | Solomon_Arch | Oceania | Austronesian | Rennell_Bellona | 159.79 | -11.3 | Choin_et_al | PASS |

| | | | | | | | | | | | | |
|---|---|---|---|---|---|---|---|---|---|---|---|---|
| B00FLJ5 | Bellona | Bellona_Rennell | NA | Bellona | Solomon_Arch | Oceania | Austronesian | Rennell_Bellona | 159.79 | -11.3 | Choin_et_al | PASS |
| B00FLHH | Rennell | Bellona_Rennell | NA | Rennell | Solomon_Arch | Oceania | Austronesian | Rennell_Bellona | 160.29 | -11.64 | Choin_et_al | PASS |
| B00FLHP | Rennell | Bellona_Rennell | NA | Rennell | Solomon_Arch | Oceania | Austronesian | Rennell_Bellona | 160.29 | -11.64 | Choin_et_al | PASS |
| B00FLHD | Rennell | Bellona_Rennell | NA | Rennell | Solomon_Arch | Oceania | Austronesian | Rennell_Bellona | 160.29 | -11.64 | Choin_et_al | PASS |
| B00FLHL | Rennell | Bellona_Rennell | NA | Rennell | Solomon_Arch | Oceania | Austronesian | Rennell_Bellona | 160.29 | -11.64 | Choin_et_al | PASS |
| B00FLG9 | Tikopia | Tikopia | NA | Tikopia | Solomon_Arch | Oceania | Austronesian | Tikopia | 168.3 | -12.29 | Choin_et_al | PASS |
| B00FLGL | Tikopia | Tikopia | NA | Tikopia | Solomon_Arch | Oceania | Austronesian | Tikopia | 168.3 | -12.29 | Choin_et_al | PASS |
| B00FLGT | Tikopia | Tikopia | NA | Tikopia | Solomon_Arch | Oceania | Austronesian | Tikopia | 168.3 | -12.29 | Choin_et_al | PASS |
| B00FLH5 | Tikopia | Tikopia | NA | Tikopia | Solomon_Arch | Oceania | Austronesian | Tikopia | 168.3 | -12.29 | Choin_et_al | PASS |
| B00FLGD | Tikopia | Tikopia | NA | Tikopia | Solomon_Arch | Oceania | Austronesian | Tikopia | 168.3 | -12.29 | Choin_et_al | PASS |
| B00FLGH | Tikopia | Tikopia | NA | Tikopia | Solomon_Arch | Oceania | Austronesian | Tikopia | 168.3 | -12.29 | Choin_et_al | PASS |
| B00FLGP | Tikopia | Tikopia | NA | Tikopia | Solomon_Arch | Oceania | Austronesian | Tikopia | 168.3 | -12.29 | Choin_et_al | PASS |
| B00FLGX | Tikopia | Tikopia | NA | Tikopia | Solomon_Arch | Oceania | Austronesian | Tikopia | 168.3 | -12.29 | Choin_et_al | PASS |
| B00FLH9 | Tikopia | Tikopia | NA | Tikopia | Solomon_Arch | Oceania | Austronesian | Tikopia | 168.3 | -12.29 | Choin_et_al | PASS |
| B00FLH1 | Tikopia | Tikopia | NA | Tikopia | Solomon_Arch | Oceania | Austronesian | Tikopia | 168.3 | -12.29 | Choin_et_al | PASS |

### Table S2.

### Austronesian- and Papuan-related ancestry proportions for the Massim groups, estimation by ADMIXTURE, GLOBETROTTER, and RFMix.

| | | ADMIXTURE | | GLOBETROTTER | | RFMix | | |
|---|---|---|---|---|---|---|---|---|
| Massim region | Group | Austronesian | Papuan | Austronesian | Papuan | Austronesian | Papuan | Uncertain |
| Collingwood Bay | Northern | 0.3 | 0.7 | 0.28 | 0.72 | 0.23 | 0.67 | 0.1 |
| | Wanigela | 0.33 | 0.68 | 0.32 | 0.68 | 0.25 | 0.65 | 0.1 |
| | Airara | 0.3 | 0.7 | 0.28 | 0.72 | 0.23 | 0.67 | 0.09 |
| Western Massim | Mainland eastern tip | 0.38 | 0.62 | 0.33 | 0.67 | 0.27 | 0.61 | 0.11 |
| | Normanby | 0.37 | 0.63 | 0.33 | 0.67 | 0.27 | 0.61 | 0.11 |
| | Fergusson | 0.36 | 0.64 | 0.34 | 0.66 | 0.26 | 0.63 | 0.11 |
| Northern Massim | Trobriand | 0.52 | 0.48 | 0.42 | 0.58 | 0.43 | 0.45 | 0.12 |
| | Gawa | 0.52 | 0.48 | 0.42 | 0.58 | 0.43 | 0.45 | 0.12 |
| | Woodlark | 0.51 | 0.49 | 0.42 | 0.58 | 0.42 | 0.46 | 0.12 |
| | Laughlan | 0.52 | 0.49 | 0.41 | 0.59 | 0.41 | 0.46 | 0.13 |
| Southern Massim | Misima | 0.43 | 0.57 | 0.37 | 0.63 | 0.31 | 0.57 | 0.12 |
| | Western Calvados | 0.42 | 0.58 | 0.38 | 0.62 | 0.32 | 0.56 | 0.12 |
| | Eastern Calvados | 0.4 | 0.6 | 0.35 | 0.65 | 0.26 | 0.63 | 0.11 |
| | Sudest | 0.27 | 0.73 | 0.31 | 0.69 | 0.17 | 0.73 | 0.1 |
| | Rossel | 0.2 | 0.8 | 0.27 | 0.73 | 0.13 | 0.8 | 0.08 |
